# Beyond G1/S regulation: How cell size homeostasis is tightly controlled throughout the cell cycle?

**DOI:** 10.1101/2022.02.03.478996

**Authors:** Xili Liu, Jiawei Yan, Marc W. Kirschner

## Abstract

To achieve a stable mass distribution over multiple generations, proliferating cells require some means of counteracting stochastic noise in the rate of growth, the time spent in the cell cycle, and the imprecision of the equality of cell division. In the most widely accepted model, cell size is thought to be regulated at the G1/S transition, such that cells smaller than a critical size pause at the end of G1 phase until they have accumulated mass to a predetermined size threshold, at which point the cells proceed through the rest of the cell cycle. However, a model, based solely on a specific size checkpoint at G1/S, cannot readily explain why cells with deficient G1/S control mechanisms are still able to maintain a very stable cell mass distribution. Furthermore, such a model would not easily account for how stochastic variation in cell mass during the subsequent phases of the cell cycle can be anticipated at G1/S. To address such questions, we applied computationally enhanced Quantitative Phase Microscopy (ceQPM) to populations of proliferating cells, which enables highly accurate measurement of cell dry mass of individual cells throughout the cell cycle. From these measurements we can evaluate the factors that contribute to cell mass homeostasis at any point in the cell cycle. Our findings reveal that cell mass homeostasis is accurately maintained, despite disruptions to the normal G1/S machinery or perturbations in the rate of cell growth. Control of cell mass accumulation is clearly not confined to the G1/S transition but is instead exerted throughout the cell cycle. Using several mammalian cell types, we find that the coefficient of variation in dry mass of cells in the population begins to decline well before the G1/S transition and continues to decline throughout S and G2 phases. Among the different cell types tested, the detailed response of cell growth rate to cell mass differs. However, in general, when it falls below that for exponential growth, the natural increase in the coefficient of variation of cell mass is effectively constrained. We find that both size-dependent cell cycle regulation and size-dependent growth rate modulation contribute to reducing cell mass variation within the population. Through the interplay and coordination of these two processes, accurate cell mass homeostasis emerges. Such findings reveal previously unappreciated and very general principles of cell size control in proliferating cells. These same regulatory processes might also be operative in terminally differentiated cells. Further quantitative dynamical studies should lead to a better understanding of the underlying molecular mechanisms of cell size control.

## Introduction

The size distribution of a population of proliferating cells is accurately maintained over many generations, despite variability in the growth rate and the duration of the cell cycle in individual cells, as well as the imprecision in the equipartition of daughter cells at mitosis. Each of these factors is known to contribute to dispersion in cell size within a in a population(1). It has long been evident that there has to be some “correction” mechanism that would act within individual cells to counteract the combined effects of the sources of random variation and thereby ensure a stable size distribution in the population. Such a mechanism would need to operate precisely over many generations(2). Studies on mammalian and yeast cell size have focused on one attractive and plausible mechanism for size homeostasis: the dependence of the G1 length inversely with size. Theoretically such a mechanism would allow small cells to spend a longer time growing in the G1 phase, thus allowing them to catch up to larger cells. Such a process would reduce cell size variation by normalizing size at the point of S phase entry(2–9). Several molecular players in this process have been suggested, such as the dilution of retinoblastoma (Rb) protein(6,9,10) and the activation of p38 MAPK kinase(11,12). However, such a mechanism, while simple, cannot in principle fully explain the constancy in the cell size distribution over many generations. In particular, if G1 length regulation were the only operative mechanism, cells would not be able to anticipate random variation introduced during the subsequent nonG1 cell cycle phases, which collectively represent a period longer than G1 in most proliferating cell types. Nevertheless, most proliferating cell populations, regardless of their surrounding environment and genetic background, manage to achieve highly accurate size homeostasis (13).

In 1985, Zetterberg et al. reported that the variation of G1 length in mouse fibroblasts accounted for most of the variation in cell cycle length when cells switched from quiescence to proliferation (14). However, a later study in several cell lines found the G1, S, and G2 phase lengths had comparable variability and were all positively correlated with the cell cycle length in normal cycling populations (15), implying a dependency of cell cycle phase lengths on cell size outside of G1. Regulation of the S and G2 lengths is known to make a contribution to size homeostasis in lower eukaryotic organisms, such as budding and fission yeasts(16–18). However, evidence of size-dependent regulation outside of G1 has seldom been reported in mammalian cells(4,7). Little is known about whether the nonG1 phases play an appreciable role in maintaining mammalian cell size homeostasis or whether variation in cell size introduced in the nonG1 phases is somehow fully compensated at the next G1/S transition.

Conceptually, an alternative approach to regulating cell size at S phase entry or in any other cell cycle phases would be to regulate *cell growth*(1,19). A few previous studies have suggested various types of size-dependent growth rate modulation in cultured cells. For instance, Cadart et al. found that the slope of volume growth rate vs. cell volume decreases for large cells at birth(7); Neurohr et al. found that volume growth rate slows down in excessively large senescent cells(20); and Ginzberg et al. found that nuclear area is negatively correlated with its subsequent growth rate at two points during the cell cycle(8). Though such observations have been noted, there is little said about their quantitative importance, and nothing is known about underlying control mechanisms. Furthermore, these various types of growth modulation were discovered in different systems using different physical proxies for cell size, such as cell volume and nuclear area. Hence, little can be said about whether these processes coexist in the same cell, are specific to certain cell types, or are only reflected in certain types of cell measurement. Furthermore, compared to studies on cell cycle control, cell growth control has received little attention; hence today the collective results are much less conclusive.

In keeping with a preceding study in bacteria(21), we distinguish between “size control” and “size homeostasis”. We use the term “size control” to refer to the regulation of the mean size, such as when the mean size in a population of cells responds to a change of environment or when cells differentiate into a different cell type; whereas we reserve the term “size homeostasis” for the control of the variance around the mean size of a population in a defined steady-state condition. Though these two processes may be related, we cannot assume that they share the same mechanism. In this study, our focus is on the less well understood and perhaps more general concept of *size homeostasis*. We have employed cultured cell lines because primary cells can take a very long time to reach a stable cell size in culture, whereas cell lines are much more stable and reproducible. Furthermore, cell lines have been well characterized; hence observations from different laboratories can be compared, and experiments can be easily replicated. Finally, we expect that size regulation occurs in all cell types, normal and transformed, embryonic and differentiated. Like other general cellular mechanisms, such as mitosis, DNA replication, and protein secretion, it is highly likely that underlying general mechanisms are conserved. To test this generality, we have studied size regulation during the cell cycle in several mammalian cell lines of diverse origins, cultured under different conditions, and compared their behaviors.

Cell size can be represented by either mass or volume. Cell volume tends to be a more passive response than mass to the mechanical and osmotic conditions occurring during the cell cycle and differentiation(22–25). Hence, we have chosen to focus on cell mass homeostasis. There are excellent experimental means to measure cell mass in suspension culture (27), but it is much harder to measure cell mass accurately when cells are attached to a substratum, which is the physiological context for most mammalian cell types. This single experimental limitation has thwarted the study of cell mass homeostasis and growth rate control in the most well studied systems. Measuring the mass of a single cell on a culture dish accurately is surprisingly difficult. Furthemore determining the growth rate from the time derivative of the mass is even more challenging(26,27). The study of cell mass growth rate regulation in attached cells with sufficient precision to distinguish between different models of growth control has required the development of new methods. To this end, we recently developed computationally enhanced Quantitative Phase Microscopy (ceQPM), which measures cell dry mass (the cell’s mass excluding water) by the refractive index difference between cell and medium to a precision of better than 2% (28). To describe statistically significant features of cell mass and growth rate regulation, we tracked single-cell growth and the timing of cell cycle events at a scale of thousands of cells per experiment. Using this improved technology, we could investigate the process of cell mass accumulation relative to cell cycle progression throughout the cell cycle and derive important new understandings of cell mass homeostasis during the cell cycle in several cultured cell lines. The results challenge existing theories of cell mass (or, more colloquially, cell size) homeostasis and suggest further mechanistic experiments.

## Results

### Cell mass variation is tightly controlled and largely independent of the state of the G1/S circuitry

“Cell mass homeostasis” can be strictly defined as the maintenance of a stable distribution of cell mass over generations in a population of proliferating cells. Expressed mathematically, at homeostasis, the coefficient of variation (CV) of cell mass at division, *CV*_*d*_, should be lower than the CV of cell mass at birth, *CV*_*b*_. And, the two should fulfill the equation adapted from Huh et al. (29):

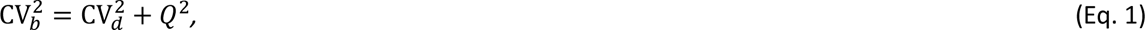

where *Q* denotes the partition error, with 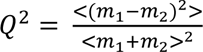; *m*_1_ and *m*_2_ are the birth masses of the two sister cells, respectively. By monitoring the proliferation and growth of HeLa cells by ceQPM, we found that the cells were indeed at such a homeostatic state, as the difference between the left- and right-hand sides of Equation 1 was negligible (Fig. S1).

To explore this further, we considered an abstract model of how the cell mass variation of a cell population evolves with cell cycle progression (SI Text, Section 1). If there were no operative controls and cell mass grew exponentially 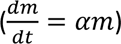 (Fig. 1A), the cell mass CV would be expected to increase super-exponentially as the cells traverse the cell cycle due to the variation of the growth exponent, *α*, among cells (Fig. 1B). Furthermore, the variation in cell cycle length and the partition error would further contribute to the cell mass CV at each generation (Fig. 1C). To maintain cell mass homeostasis, these accumulated discrepancies must be offset by a *reduction* of variability by some process during the cell cycle. If, as suggested in both in vitro and in vivo systems(4,6), the G1/S checkpoint were the principal “size control checkpoint” (Fig. 1A), we would expect the reduction in cell mass variation to occur before or at the G1/S transition. The cell mass CV would then be expected to increase super-exponentially after G1/S due to the lack of any operable size control processes in the nonG1 phases. Therefore, the CV reduction before G1/S would have to greatly undershoot the birth mass CV to anticipate and compensate for the cell mass variability that would accumulate during the nonG1 phases (Fig. 1B). If the G1/S control were weakened by genetic mutation or pharmacological perturbation (Fig. 1A), the reduction in cell mass CV before G1/S would be expected to decrease, and the uncorrected error would cause an increase in the division mass CV (Fig. 1B). Such a population would eventually reach a new homeostatic state with higher birth and division mass CVs in order for Eq. 1 to be fulfilled (Fig. 1C). Therefore, the birth mass CV at homeostasis can be used to indicate the stringency of the control on cell mass homeostasis.

**Figure 1.**
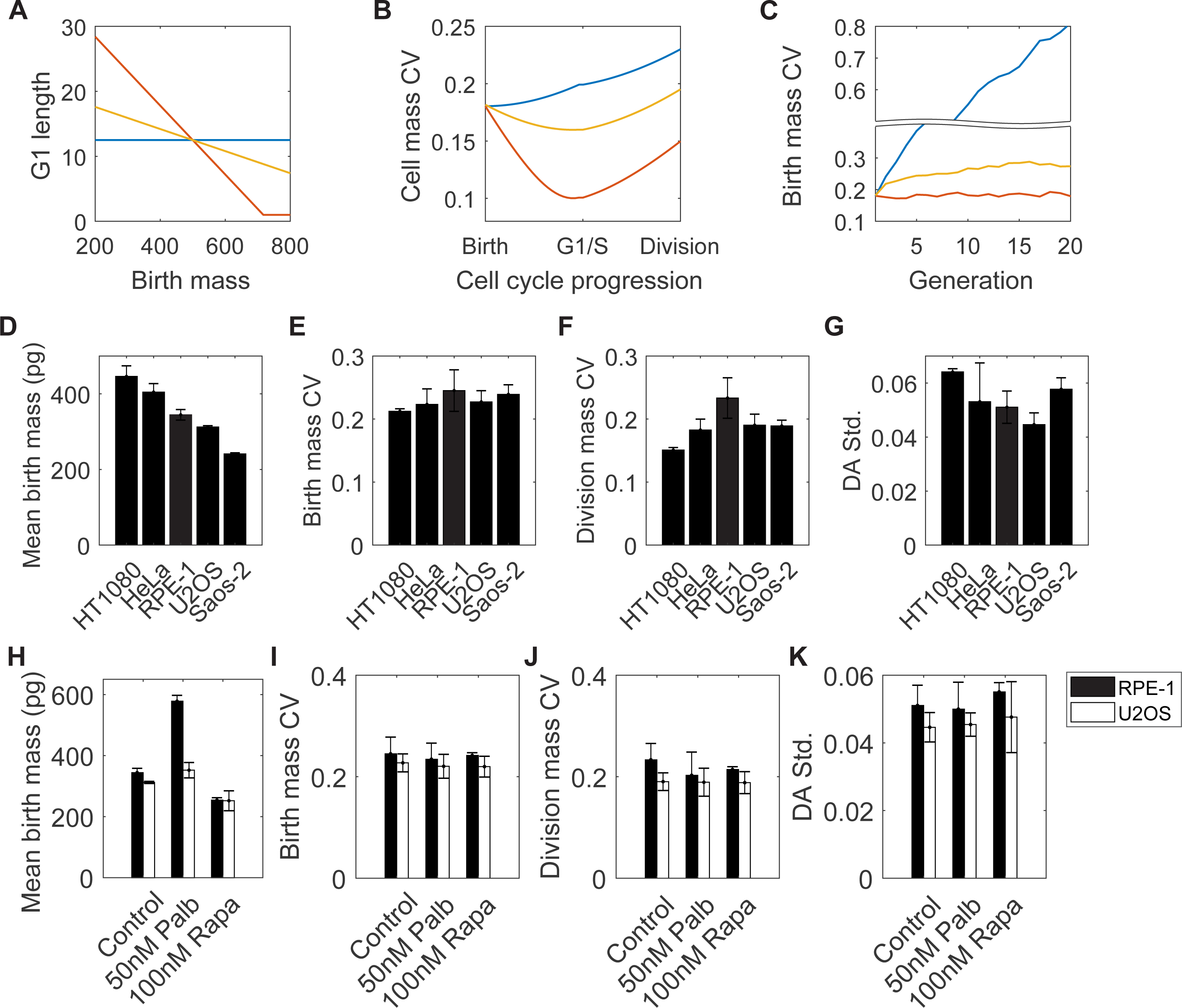
Cell mass variation is tightly controlled in mammalian cell lines and is robust to perturbations in G1/S regulation or growth rate. (A-C) An abstract model of cell mass homeostasis at different G1 regulation strengths, represented by the slope of G1 length vs. birth mass correlation. The corresponding model and simulation parameters are in the SI Text, Section 1. In the model, we assume cells grow exponentially, and the G1 length control is the only mechanism to reduce cell mass variation. (A) Correlations between G1 length and birth mass. Blue: no G1 length control; Red: with strong G1 length control; Yellow: with weak G1 length control. (B) Cell mass CV changes with cell cycle progression during one cell cycle with the corresponding G1 length regulation in (A). (C) Birth mass CV changes across generations with the corresponding G1 length regulation in (A). (D-G) The mean birth mass (D), birth mass CV (E), division mass CV (F), and standard deviation of Division Asymmetry (DA std.) (G) for different cell lines. (H-K) The mean birth mass (H), birth mass CV (I), division mass CV (J), and DA std. (K) of RPE-1 and U2OS cells in normal culture medium, medium with 50 nM palbociclib, and medium with 100 nM rapamycin at cell mass homeostasis. Error bars in (D-K) indicate the standard deviation of three or more measurements.

To investigate how different forms of G1/S control affect cell mass homeostasis, we compared various human cancer cell lines, each with different G1/S deficiencies, and RPE-1, a cell line with a wild-type G1/S transition(7,12,30) (Table S1). To evaluate the stringency of the control mechanism on cell mass homeostasis, we measured the birth and division mass CVs of live cell populations from short-term videos using ceQPM. We define the Division Asymmetry, 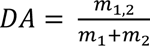, where *m*_1_ and *m*_2_ represent the birth masses of the two daughter cells, and *m*_1,2_ denotes the mass of either of the daughter cells. For a population that divides with perfect symmetry, the distribution of *DA* should be precisely at 0.5 without any dispersion. But if either daughter cell were larger or smaller than half the original size, its *DA* would deviate from 0.5. The standard deviation of *DA* (DA std.) quantitatively represents the fidelity of cytokinesis, and it is more commonly used than the partition error *Q* in Eq. 1(17,31). Despite the considerable variation in cell mass across the different cell lines (the mean birth mass of the largest cell line, HT1080, is 1.85-fold greater than the smallest cell line, Saos-2) (Fig. 1D), the difference in birth mass CV is less than 15% for each cell line (Fig. 1E); the division mass CV and DA std. for these cell lines were also comparable (Fig. 1F, G). Note that the measurement error of ceQPM is negligible (less than 2%) compared to the birth and division mass CVs.

To assess the robustness of the birth mass CV to perturbations in the G1/S transition, we perturbed G1/S regulation in both RPE-1 and U2OS cells using a well characterized CDK4/6 inhibitor, palbociclib(32). Although U2OS cells have intact Rb proteins, which have been reported to govern the G1/S transition(4,6,33), they carry deficiencies in other G1/S regulators (Table S1) and are much less sensitive to palbociclib than RPE-1, which has intact G1/S circuitry (Fig. S2A, B). Both cell lines were examined at a low dose of palbociclib, where there was a delay in G1/S but no arrest of the cell cycle (11). We measured the dry mass of RPE-1 and U2OS cells after being cultured for more than one week in palbociclib, at which point the mass distribution of each cell line had reached a new steady state. It had been shown previously that a low dose of palbociclib weakens the negative correlation between birth size and G1 length (like the yellow curve in Fig. 1A)(11). Thus, if G1 regulation were essential for cell mass homeostasis, we would expect the birth mass CV to increase with palbociclib treatment (like the yellow curve in Fig. 1C). Surprisingly, although the mean mass at birth had increased by 1.68 fold and 1.13 fold, respectively in RPE-1 and U2OS cells (Fig. 1H), the birth mass CV for either cell line hardly changed and in fact slightly decreased (a 4% and 3% reduction for RPE-1 for U2OS cells, respectively) (Fig. 1I). Similarly, the division mass CV and the standard deviation of Division Asymmetry, DA std., also hardly changed after exposure of both cell lines to palbociclib (Fig. 1J, K). These very small changes in mass CVs indicate that the control of mass homeostasis still operates accurately, despite strong perturbation of the G1/S transition.

Since disruption and delay of the cell cycle at G1/S did not appear to affect size homeostasis, we examined the inhibition of cell growth for effects on cell size variability. We used rapamycin, a specific inhibitor of mTOR(34), which has pervasive knock-on effects on protein synthesis and degradation(35). When RPE-1 and U2OS cultures were exposed to rapamycin, the steady-state birth mass decreased by 27% and 20%, respectively (Fig. 1H). However, there were no significant changes in the birth mass CV, division mass CV, or Division Asymmetry (changes less than 8% were observed) (Fig. 1I-K). Therefore, it appears that mass homeostasis is strongly buffered, even when mass is greatly perturbed.

### Cell mass variation is regulated throughout the cell cycle

Using ceQPM, we can now ask at what points during the cell cycle does variation in cell size occur and at what points is it suppressed. We used the Coefficient of Variation as a metric of cell mass variation and measured it *throughout* the cell cycle in live RPE-1 and HeLa cells (Fig. S3A). To correlate the CV with the state of the cell cycle, we utilized fluorescently tagged geminin degron as the cell cycle marker. Geminin is a protein that regulates DNA replication. Possessing a destruction sequence like cyclin B, geminin is degraded precisely at mitosis and starts its accumulation precisely at the G1/S transition (Fig. S3B)(36). We aligned individual cell mass trajectories by normalizing the length of the G1 segment to 0-0.5 and that of the nonG1 segment to 0.5-1 and then calculated the CV of these normalized cell mass trajectories with cell cycle progression. In RPE-1 cells, the cell mass CV was found to be reduced throughout the cell cycle (Fig. 2A), whereas in HeLa cells, the cell mass CV increased in the G1 phase before declining in the nonG1 phases (Fig. 2B). Neither cell line exhibited its lowest cell mass CV at the G1/S transition, as would be predicted by conventional G1 length control models (Fig. 1B).

**Figure 2.**
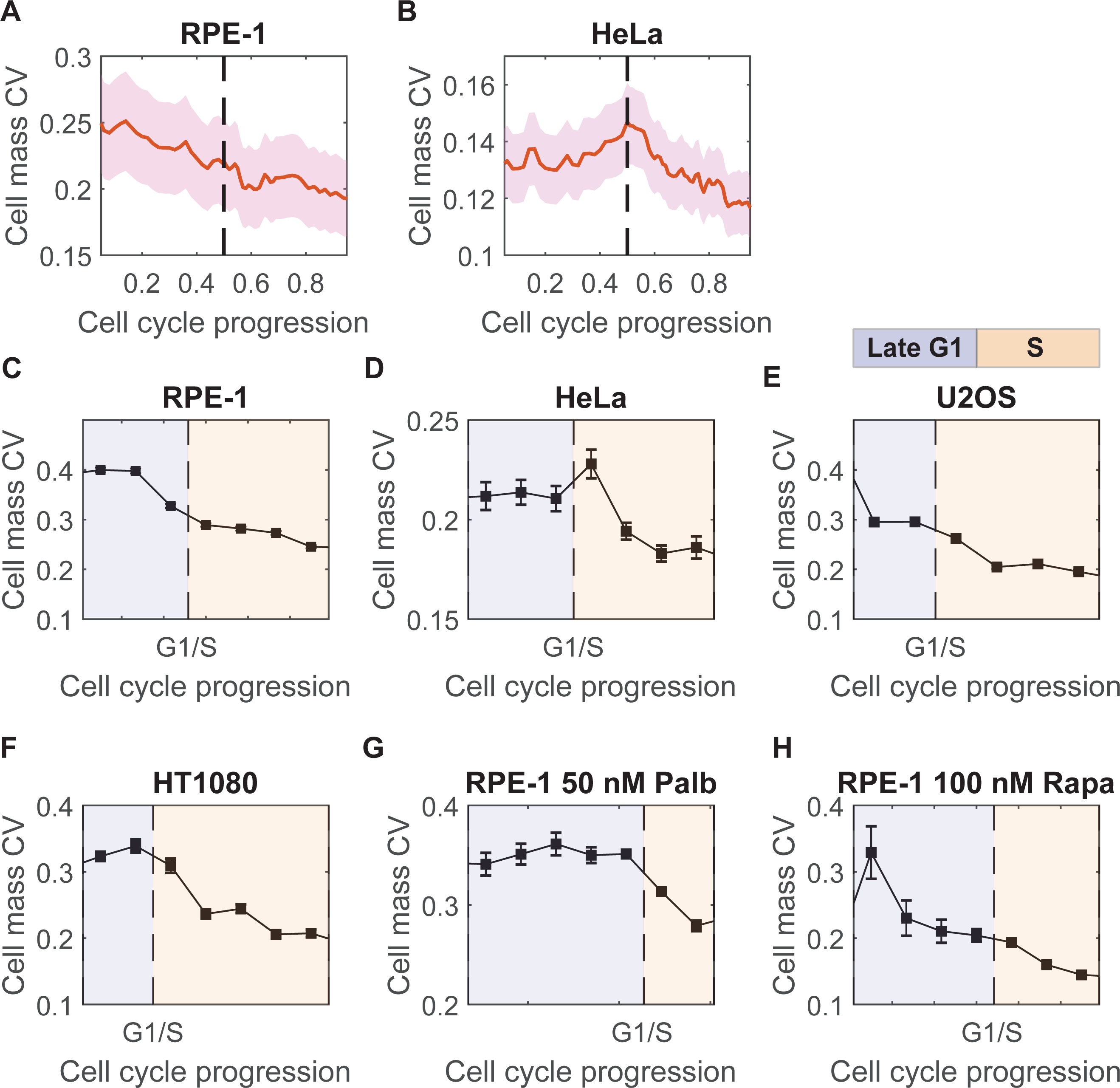
Cell mass variation is regulated throughout the cell cycle. (A, B) Cell mass CV change with cell cycle progression measured in live RPE-1 (n= 89) (A) and HeLa cells (n = 223) (B). The red solid lines denote the cell mass CV of the population; the pink shadows show the 95% confidence interval; the dashed line indicates the G1/S transition. (C-H) The profile of how cell mass CV changes with cell cycle progression at cell mass homeostasis measured in fixed RPE-1 (C), HeLa (D), U2OS (E), and HT1080 (F) cells, as well as RPE-1 cells that had reached the new cell mass homeostasis with 50 nM palbociclib (G) or 100 nM rapamycin (H). The cell cycle stages were identified by DNA content and log(mAG-hGeminin) as illustrated in Fig. S4 B-F, H, and J; the late G1 and S phases are indicated by areas shaded in purple and orange, respectively; error bars are the standard error of CV, 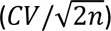, where *n* is the cell number at the corresponding cell cycle stage (n>135 for all conditions).

To examine the regulation of cell mass variation in various cell lines and under different conditions, we calculated the cell mass CV profile as a function of cell cycle progression from fixed cells, which provided much higher throughput than our live cell measurements. Using Ergodic Rate Analysis (ERA)(37), we defined a cell cycle mean path and divided it into 13-14 segments evenly spaced in time, using the measurements of DNA content and the fluorescently-tagged geminin degron. We applied this approach to hundreds of thousands of fixed cells (Fig. S4A). DNA replication occurs exclusively in the S phase, whereas geminin accumulation starts at the G1/S transition (Fig, S4B-E, H, J)(37). Though these two markers provide good resolution in late G1 and S phases, they have poor temporal resolution in the early G1 and G2-M phases due to inaccuracy in cell cycle stage identification (Fig. S4F). Therefore, we focused our analyses exclusively on the cell mass CV in the late G1 and S phases, employing large numbers of fixed cells.

We applied this approach to four cell lines: RPE-1, HeLa, U2OS, and HT1080. The cell mass CV profiles in fixed RPE-1 and HeLa cells (Fig. 2C, D) were similar to what we had previously found in the live cell trajectories (Fig. 2A, B), further validating the use of fixed cells to extract cell mass CV profiles. We found that in RPE-1 and U2OS cells, the cell mass CV declined in late G1 (Fig. 2C, E), as would be expected from conventional models where regulation of the G1/S transition is thought to be the sole means for normalizing cell size. However, we were surprised to find that the CV of cell mass then decreased progressively through S phase. Most strikingly, in HeLa and HT1080 cells, there was virtually no reduction in cell mass CV in late G1; the major decrease only took place in S phase (Fig. 2 D, F). These quantitative differences in cell mass CV profiles may depend on the status of the G1/S circuitry in these cell lines (Table S1). These observations are completely at odds with the G1/S transition playing the dominant role in cell size control, although it may remain a critical point for cell cycle regulation(1,19,33). Note that the decrease in cell mass CV cannot be explained by a reduction in noise because even if noise went to zero at some point, the CV would remain at its previous value. We believe that very strong conclusions can be drawn from these phenomenological measurements: They suggest that there must be feedback between cell size and cell growth rate or between cell size and cell cycle outside of the G1/S transition. The effect of this feedback would be to effectively reduce existing variation in the population in nonG1 phases of the cell cycle.

Since palbociclib and rapamycin had little or no effect on the birth and division mass CVs (Fig. 1I, J), we wondered whether they affected the timing of mass CV regulation during the cell cycle. Consequently, we carefully measured the cell mass CV profiles in fixed RPE-1 cells that had reached new cell mass homeostasis with either drug. Both drugs altered the duration of the cell cycle phases and particularly extended the G1 phase (Table S2). As we had done above with untreated cells, we computed the cell cycle mean path of treated cells and examined their cell mass CV as a function of cell cycle progression (Fig. S4G-J). Strikingly, we found that disrupting the G1/S transition with palbociclib led to a slight increase in cell mass CV in late G1, followed by a much greater reduction in cell mass CV during the S phase (Fig. 2G). Conversely, inhibiting cell growth with rapamycin caused a greater reduction of cell mass CV in late G1, and the reduction in S phase became smaller (Fig. 2H). These results suggest that the regulation of mass CV during S phase compensates for the mass CV reduction in late G1. Thus, when there is an insufficient or excessive reduction in mass CV in late G1 due to the inhibition of the G1/S transition or growth, respectively, there is a corresponding change in the mass CV in S phase, which acts to maintain the mass CV at division at the same level.

### Feedback by cell mass on the duration of G1, but also on the duration of S and G2 phases

To investigate further cell size regulation outside of the G1 phase, we needed to better optimize the resolution of the cell cycle markers we had employed. We therefore adopted two cell cycle markers for live cells that bracketed S phase: mAG-hGeminin(36) and mTurquoise2-SLBP(38). The APC^Cdh1^ substrate, geminin, starts to accumulate in the nucleus at S phase entry(39), whereas the histone mRNA stem-loop binding protein, SLBP, is rapidly degraded at the end of the S phase(40)(Fig. S5A). Unlike the conventional PCNA or DNA ligase I markers, which label replication foci during the S phase (41,42), geminin and SLBP are diffusive in the nucleus and more suitable for the relatively low spatial resolution of the QPM camera. With these two markers, we could accurately quantify the duration of G1, S, and G2-M phases. Since the duration of M phase is remarkably constant (15), we attributed most of the variation in G2-M duration to the G2 phase itself. We verified that the timing of S phase, as identified by geminin and SLBP, was consistent with the timing of S phase identified by the DNA ligase I foci (Fig. S5B-C). None of the markers affected the length of any of the cell cycle phases, nor did they affect the size-dependent regulation of the duration of the cell cycle phases (Table S3). Moreover, the identification of the cell cycle phases (G1, S, and G2-M) using geminin and SLBP exhibited a similar variability in their lengths as those shown using PCNA as a marker of S-phase by Araujo et al. (15) (Table S2). Therefore, we were confident that the geminin and SLBP markers faithfully reported the cell cycle phase durations and did not change the physiology of these processes.

We confirmed the well-established existence of cell size-dependent regulation of G1 length using ceQPM. Consistent with previous findings(3,4,6–8,12), we found that the G1 length was negatively correlated with birth mass in both non-transformed and transformed cell lines, RPE-1 (Fig. 3A) and HeLa (Fig. 3E), respectively. The correlation was stronger in RPE-1 than in HeLa cells (Fig. 3A, E, Table S4). We also investigated the size-dependent regulation of the durations of both S and G2 phases. S and G2-M phase lengths negatively correlated with the initial mass of the corresponding periods in both RPE-1 and HeLa cells (Fig. 3B-C, F-G). For RPE-1 cells, the correlations of cell cycle phase length with initial mass in S and G2 were weaker than that in G1, yet they were significant (Fig. 3A-C, Table S4), demonstrating that S and G2 regulation of cell mass variation can occur in a non-transformed cell with an intact cell cycle network, including an intact G1/S transition. In contrast to the conventional models that would have predicted G1 length to vary inversely with mass while leaving other phases unaffected, we found that a negative correlation of cell cycle phase length with mass was much stronger in the S phase in HeLa cells. This negative correlation of the length of S phase with mass was significantly stronger than its effect on the G1 phase, with a correlation coefficient of −0.29 for the S phase versus −0.20 for the G1 phase (Fig. 3E-F, Table S4). It is worth noting that although RPE-1 has more stringent G1/S control than HeLa, the overall dependency of cell cycle length on cell mass was not stronger (Fig. 3D, H, Table S4). These studies challenge the G1/S checkpoint model, as size-dependent cell cycle regulation is not restricted to the change in the length of G1 phase as predicted (2,43,44), but rather it is accompanied by changes in the lengths of the other phases of the cell cycle.

**Figure 3.**
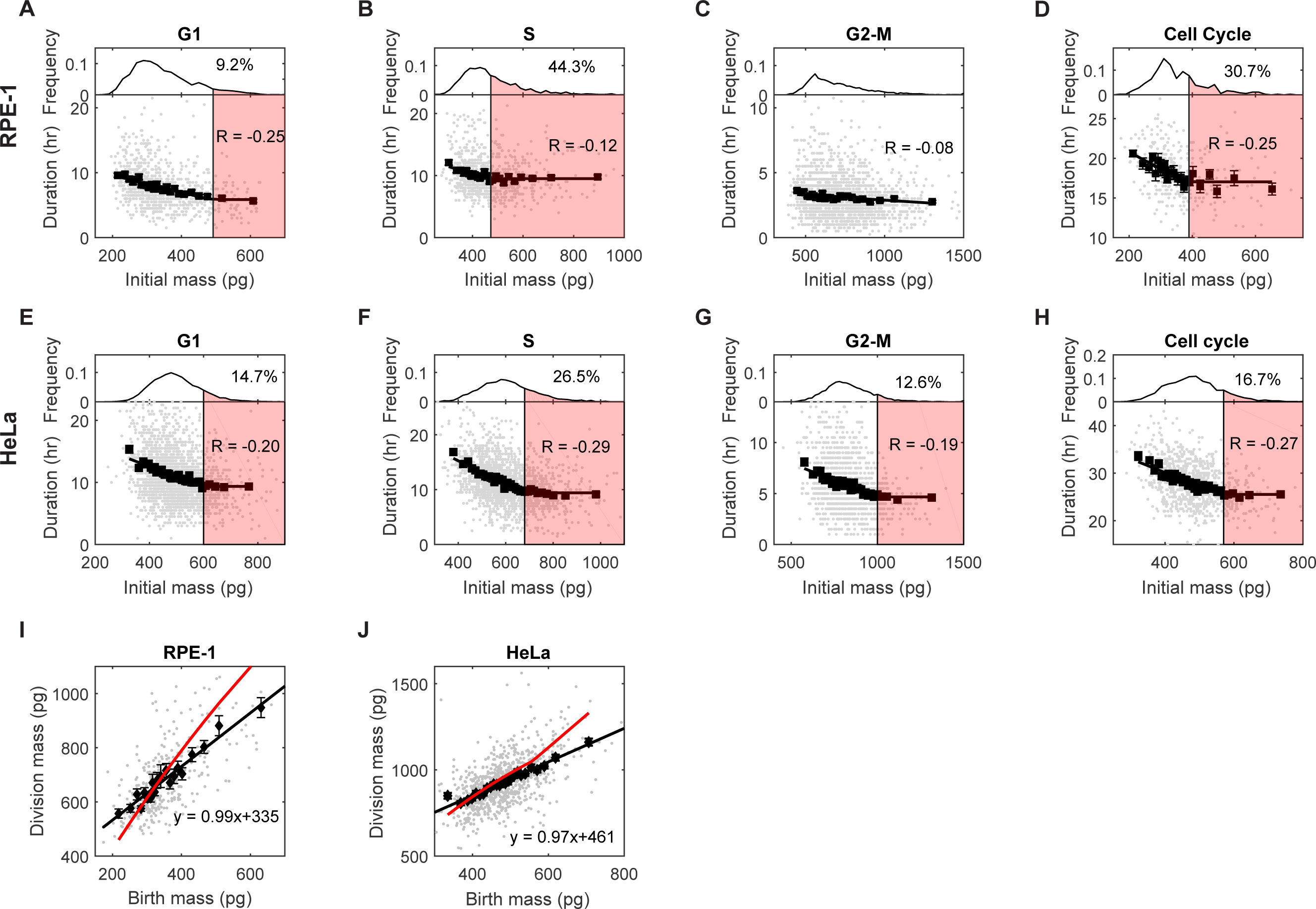
The negative regulation of the durations of the G1, S, and G2 phases by cell mass. (A-D) The correlation between G1 (A), S (B), G2-M (C), and full cell cycle (D) length and the initial mass of the corresponding period in RPE-1 cells. The bottom panels indicate the correlation; the top panels are the distributions of the initial mass. Each gray dot in the bottom panels is an observation; R is the correlation coefficient of the gray dots; black squares indicate the average of each cell mass bin; error bars are the standard error of the mean (SEM); solid black line is the best fit of the black squares (Table S4). The red shaded area in the top panel indicates the cell mass range that affected by the minimal cell cycle phase length limit, with the text indicating the percentage of affected cells in the distribution. (E-H) The correlation between G1 (E), S (F), G2-M (G), and full cell cycle (H) length and the initial mass of the corresponding period in HeLa cells. (I-J) The correlation between birth and division masses in RPE-1 (I) and HeLa (J) cells. Each gray dot is an observation; black squares are the average of each cell mass bin; error bars are SEM. The solid black line is the best linear fit of the gray dots; the text indicates the function of the best fit; the red line is the prediction of the best fit in (D) or (H), respectively, assuming that cells grow exponentially (Materials and Methods).

Upon closer examination of the binned correlations, we observed a fixed minimum limit for the length of nearly every phase of the cell cycle, as well as the length of the entire cell cycle in RPE-1 and HeLa (Fig. 3A-B, D-H). These limits are not further reduced in large cells. To summarize these findings we employ two graphical representations for these correlations: a linear model and a bilinear model, comprised of two line segments. With these we fit the binned correlations of mass and cell cycle progression. We found that a bilinear model provided a better fit for all phases of RPE-1 and HeLa cells, with the exception of the G2-M phase in RPE-1 cells (Fig. 3A-H)(Table S4). This graphical relationship implies that regulating the duration of cell cycle phases cannot effectively control the mass of large cells. To illustrate the impact of the minimal cell cycle length on cell mass variation, we conducted simulations to observe the mean and CV of cell mass within a cell population across generations, while varying the fraction of cells affected by the minimal length limit (SI Text, Section 2). These simulations show that as the minimal cell cycle length applies to more and more cells, the homeostatic birth size CV increases. The system eventually loses homeostasis when the minimal cell cycle length is imposed on more than 40 percent of the cell population (Fig. S6).

In these experiments, we found that the slope of a graph of birth masses versus division masses was close to 1 in both RPE-1 (Fig. 3I) and HeLa cells (Fig. 3J), consistent with the adder-like behavior seen previously (7). The adder model predicts that cells add a constant amount of mass during the cell cycle, regardless of their birth mass. In our measurements, each cell cycle phase exhibited an adder-like behavior (Fig. S7), making the full cell cycle a sequential adder. Such sequential adder behaviors challenge the interpretation that, in mammalian cells mass regulation arises from a combination of a G1 sizer and a nonG1 timer (19). Rather, the present findings strongly suggest that each cell cycle phase, except for M phase, contributes to cell mass homeostasis. Moreover, the fitted function of birth mass and cell cycle length correlation cannot fully explain the adder behavior. This is particularly the case for large cells, under the assumption of exponential growth (Fig. 3I-J). This discrepancy is at least partially due to the existence of a minimal cell cycle phase length. These new results underscore the need for a process of non-exponential growth (or what we term “growth rate modulation) to maintain cell mass homeostasis in the mammalian cells we have studied, rather than relying solely on processes of cell cycle regulation.

### How size-dependent growth rate modulation reduces the CV of cell mass during cell cycle progression

The simplest mathematical model for cell growth kinetics, which requires no mass sensing or feedbacks, is an exponential model where the growth rate is proportional to mass. This has been particularly successful in describing growth in bacteria and can be rationalized by a process of ribosome-dependent ribosome biosynthesis (45,46). This simple exponential model, however, causes variation in cell size to amplify as cells progress through the cell cycle (Fig. 1B, SI Text, Section 3.1). Contradicting this model, several studies have found that although large cells generally grow faster than small cells, growth is not precisely exponential in mammalian cells(7,28,45), and this might lead to a reduction in cell size variation. Various previous studies suggested the dependency of growth rate on cell mass changes with cell mass and cell cycle stage(7,8,20,47–50). Recent studies by us and others have found growth rate oscillations(28,51), where a cell alternates between increases and decreases in growth rate.

To explore the dependence of growth rate on cell mass in proliferating cells, we measured the growth rate in a 3-hour time window and computed its correlation with cell mass at 0 hours. We examined how growth rate correlated with cell mass in 25,000 HeLa cells and found that the relation of mass to growth was close to exponential, except for a slight depression for large cells (Fig. S8A). Nevertheless, when we segregated the cells into four cell cycle phases, we uncovered distinct cell cycle dependencies in such correlations, which were originally masked by pooling all cells for analysis (Fig. S8B). An even closer look at the data, with cells categorized into 14 equally divided cell cycle stages, revealed positive-to-negative correlation transitions at various points in the cell cycle (Fig. S8C). The slope of the linear relation between cell mass and growth rate for cells in different stages of the cell cycle indicated stronger modulation (greater deviation from the expected slope of exponential growth) in the late G1 and G2-M phases (Fig. S8D), consistent with Fig. S8B and previous studies (8,37). However, the proportionality is sub-exponential in most of the cell cycle stages (Fig. S8D), suggesting a global process that inherently limits the growth of large cells.

When we investigated the mass vs. growth correlations in finely divided cell cycle stages we found subtle features. Yet, such studies require very large numbers of cells and very accurate growth rate measurements. Coarser cell cycle discrimination leads to a loss of this kind of information on subtle changes in the growth rate, it nevertheless adds greater statistical power to conclusions about overarching aspects of size-dependent growth regulation. Therefore, there is a practical tradeoff between high cell cycle resolution of the growth analyses and the statistical reliability of the findings. In the following analyses, we aimed for stronger statistical significance and therefore partitioned cells more crudely into the G1 and nonG1 phases, focusing on the most salient features of growth rate modulation. This level of resolution was sufficient to reveal previously undiscovered features, which serve to correct our current understandings.

Measuring the correlation between cell mass and growth rate in five different cell lines, we found that each cell line behaved somewhat differently. In RPE-1 cells, growth was proportional to cell mass, but the proportionality was much less than exponential, with a significant nonzero intercept (Fig. 4A). In HeLa cells, the proportionality between growth and cell mass is much closer to, but slightly less than exponential in both G1 and nonG1 phases (Fig. 4B). The observed mass vs. growth correlations in short-term measurements in RPE-1 and HeLa cells were consistent with their long-term growth trajectories (Fig. S9), showing nearly linear growth in RPE-1 and a slight deviation from exponential growth in HeLa cells. Therefore, we can confirm that the observed deviation from exponential growth is not due to inspection or sampling bias caused by the short-term measurement(52), but truly signifies the inherent growth law of the cells. In U2OS cells, the correlation was close to exponential for all cells in nonG1 and most cells in G1 phase, but it was abruptly negative for the 15% largest cells in G1 phase (Fig. 4C). In HT1080 cells, growth was close to exponential for small cells but transitioned to nearly linear growth in large cells during both G1 and nonG1 phases (Fig. 4D). A bilinear model provided a significantly better fit than a simple linear model for cells in the nonG1 phase, indicating the significance of the transition in mass vs. growth correlations as cells became larger (Table S5). In Saos-2 cells, growth was exponential except for a slight deviation for large cells in nonG1 phase (Fig. 4E). Taken together, these results indicate that the mathematical description of growth rate is not simply exponential in the cell lines we have investigated, and that different cell lines display different characteristics of mass dependency at different phases of the cell cycle.

**Figure 4.**
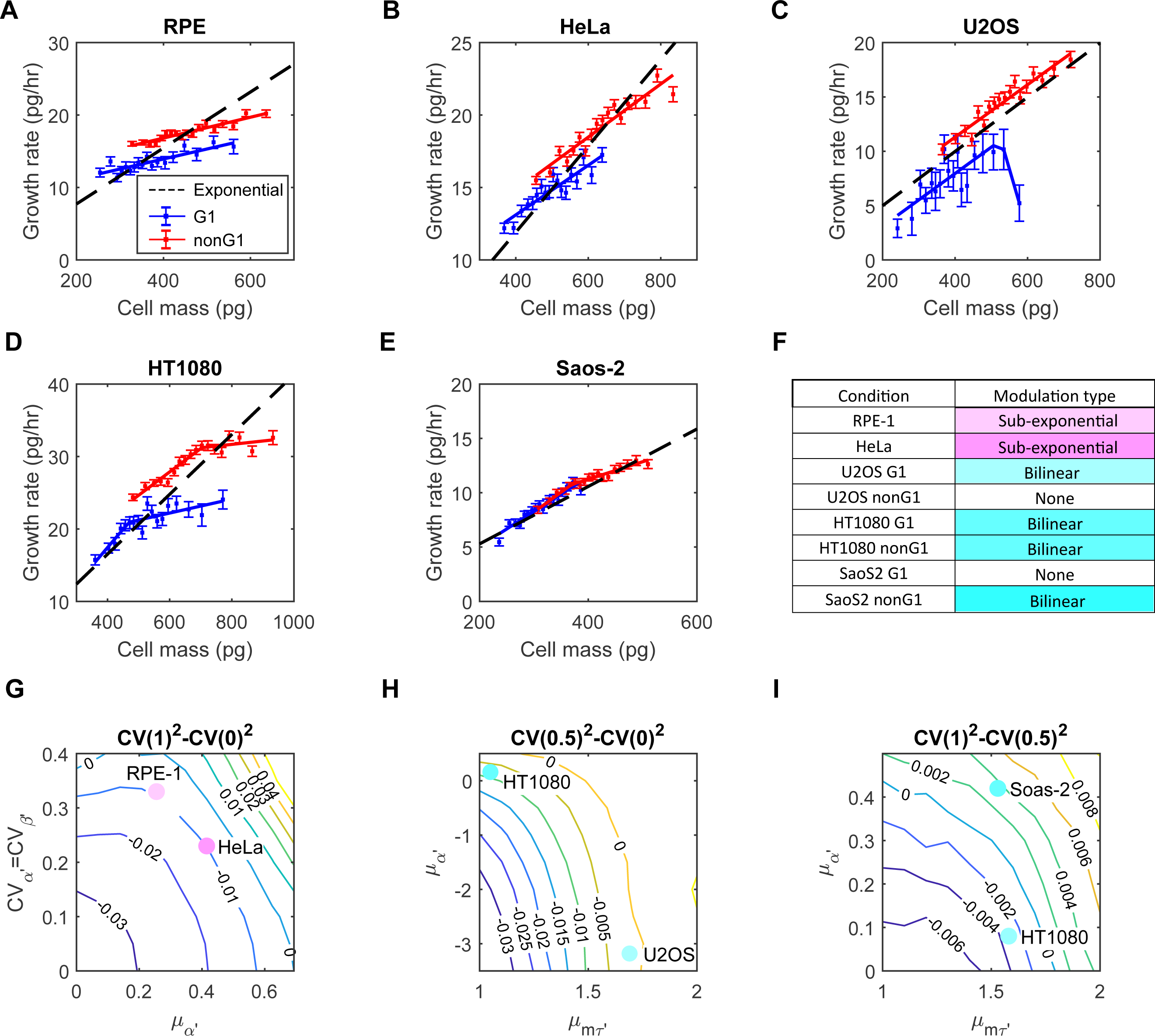
Growth rate dependence on mass differs in different cell lines, and growth rate modulation can effectively reduce cell mass CV during the cell cycle. (A-E) Correlations between cell mass and growth rate in the G1 (blue) and nonG1 (red) phases for RPE-1 (A), HeLa (B), U2OS (C), HT1080 (D), and Saos-2 (E) cells. Filled squares represent the median growth rate of each bin; error bars show SEM. The black dashed lines indicate the expected behavior for exponential growth. The solid blue and red lines are the best fit of the filled squares (Table S5). (F) The observed conditions were categorized into three types: Sub-exponential, Bilinear, and no modulation. (G) Contour plot illustrating the change in cell mass CV during the entire cell cycle for Sub-exponential growth rate modulation (SE), as a function of the mean and CV of *α*^′^ and *β*^′^, obtained from numerical simulations (SI Text, Section 3.1). (H-I) Contour plots illustrating the change in cell mass CV during the G1 (H) and nonG1 phases (I) for Bilinear growth rate modulation (BI), as a function of the means of *α*^′^ and *m*^′^, obtained from numerical simulations (SI Text, Section 3.2). These simulations assumed a 20% CV in *α*^′^. Solid circles in (G-I) indicate the corresponding positions in the contour plots when adopting parameter values from the experimental data.

To better compare the behaviors of different cell lines, we normalized the mass vs. growth correlations, using the means of birth mass and cell cycle length (Table S6). Since DNA copy number affects the correlation intercepts (Fig. 4A-D), we focused solely on the slope of the correlations. We can distinguish two general types of growth rate modulation (Fig. 4F, Table S6). In the first type, growth is linearly related to cell mass, but with a slope lower than exponential growth (RPE-1 and HeLa). We refer to this as Sub-exponential (SE) Modulation. In the second type, the slope of the mass vs. growth correlation is close to exponential for small cells but becomes less positive or even negative for large cells (U2OS G1, HT1080, and SaoS-2 nonG1). We refer to this as Bilinear (Bl) Modulation. For U2OS cells in the nonG1 phases and SaoS-2 cells in the G1 phase, the correlation slope is not significantly different from exponential growth, suggesting minimal regulation.

Other studies had proposed that growth rate modulation contributes to cell mass homeostasis(1,7,8,19,37). However, most of these claims were speculative and lacked sufficient quantitative support. The work by Cadart et al. in 2018 stands out as an exception, as it quantitated the correlation between birth mass and growth rate(7). Accurate and quantitative correlations between growth rate and cell mass are essential for a thorough assessment of the impact of growth rate regulation. Nevertheless, due to the scarcity of high-quality experimental data, most theoretical investigations into cell mass homeostasis have disregarded growth rate regulation completely and focused solely on the regulation of cell cycle length, often assuming exponential growth(53–56). In this study, we address this gap in previous studies by investigating theoretically whether the types of growth rate modulation we observed could effectively reduce cell mass variation. Using stochastic models and simulations, we focused on the influence of growth rate modulation and growth rate noise on the cell mass CV over one cell cycle. Initially for convenience, we assumed that all cells divided at the same cell cycle length. Subsequently in more comprehensive models, we incorporated cell cycle regulation and noise, as discussed in a later section.

In the absence of any growth rate modulation, we might imagine that cell mass should accumulate exponentially, as has been found in bacteria(45). This would cause the cell mass CV to increase super-exponentially due to stochastic variation in growth rate (SI Text, Section 3.1). When growth rate modulation is in the Sub-exponential (SE) form (Fig. 4F, Table S6), the slope of the correlation between cell mass and growth rate is lower than that of exponential growth. This can be described by the equation: 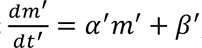, where the growth rate is composed of two terms: *α*^′^*m*^′^ represents the part of growth rate proportional to cell mass, whereas *β*^′^ represents the part independent of cell mass. Here, *m*^′^ and *t*^′^ are the cell mass and cell cycle progression time normalized by the means of birth mass and cell cycle length, respectively (SI Text, Section 3.1). According to the definition of sub-exponential growth, the mean of *α*^′^ is smaller than *ln*2 and greater than 0, and the mean of *β*^′^is determined by *α*^′^when assuming that the mean division mass is twice the mean birth mass, a requirement for maintaining mass homeostasis. For simplicity, we first assumed that *α*^′^and *β*^′^ have the same CV, but we also examined how the CV of either parameter affected the results in the supplementary information (Fig. S10).

During the initial stages of the cell cycle, the cell mass CV consistently decreases, with the rate of decrease negatively correlated with the mean of *α*^′^ and independent of the CVs of *α*^′^ and *β*^′^ (Fig. S10A, C, F, SI Text, Section 3.1). As the cell cycle progresses, the rate of mass CV reduction slows down, and the mass CV may even increase during the later period of the cell cycle (Fig. S10B, D, G, SI Text, Section 3.1). The overall change in the cell mass CV throughout the cell cycle depends on both the mean of *α*^′^ and the CVs of *α*^′^ and *β*^′^. The smaller mean of *α*^′^and lower CV of *α*^′^ and *β*^′^ result in a more significant reduction in the cell mass CV (Fig. 4G, Fig. S10E, H). In summary, growth rate variability (characterized by the CVs of *α*^′^and *β*^′^) amplifies cell mass variation, while strong growth rate modulation (small *α*^′^) can reduce cell mass variation throughout the cell cycle.

To assess whether growth rate modulation in RPE-1 and HeLa cells can cause cell mass CV reduction throughout the cell cycle, we derived the parameters from the experimental data. The mean of *α*^′^ was determined based on the mean correlations in Fig. 3A and B (Table S6). To estimate the variation in *α*^′^, we used long-term live-cell growth trajectories. The CV of *α*^′^ was found to be independent of cell mass (Fig. S11A-B). The variability in *α*^′^arises from two sources: stochastic partitioning of cellular contents during cell division (extrinsic variability) and intrinsic fluctuations in biochemical reactions (intrinsic variability) (57). The former, determined at birth, is a major contributor to cell mass variation, while the effect of the latter gradually cancels out over time, exerting minimal impact on cell mass variation.

Therefore, we focused on the extrinsic variability and estimated it by calculating the variation among the means of individual growth trajectories (Fig. S11C). The CV of *α*^′^ was estimated to be 0.33 for RPE-1 and 0.23 for HeLa cells, respectively. It is challenging to isolate the variation in *β*^′^from measurement error, thus we conducted simulations with *β*^′^ having the same CV as *α*^′^ or with the CV of *β*^′^ being equal to zero. Using these parameters, we found that both RPE-1 and HeLa cells could reduce the cell mass CV after one cell cycle (Fig. 4G, Fig. S10E, H). Since the minimal requirement for cell mass homeostasis is to have a lower cell mass CV at division than that at birth, we concluded that growth rate modulation is sufficient to maintain cell mass homeostasis in RPE-1 and HeLa cells.

When a plot of growth rate versus mass is in a Bilinear (BI) form (Fig. 4F, Table S6), the slope of the mass vs. growth correlation is close to exponential for small cells and becomes less positive or even negative in large cells. This can be described by the equation: 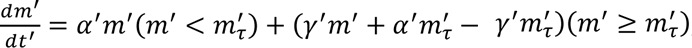, where the mean of *α*^′^ is close to *ln*2 and the mean of *γ*^′^ is smaller than *ln*2 (SI Text, Section 3.2). The first term on the right side of the equation represents the exponential portion of the mass vs. growth rate correlation, while the second term describes the part where growth rate modulation takes effect. Here, *γ*^′^indicates the strength of modulation and *m*_*τ*_′ signifies the cell mass at which this modulation begins to take effect. Both *γ*^′^and *m*_*τ*_′ are normalized by the means of cell cycle length and birth mass, respectively. Our findings indicate that the increase in cell mass CV is primarily driven by the CV of *α*^′^(Fig. S12A-D). Additionally, we investigated the impact of the means of *γ*′, and *m*_*τ*_′ on the change in cell mass CV throughout the cell cycle. We found that the smaller the means of *γ*^′^ and *m*_*τ*_′, which means the stronger the modulation on growth rate and the more cells it affects, the greater the cell mass CV reduction (Fig. 4H-I, Fig. S12).

To assess whether the growth rate modulation on its own in U2OS, HT1080, and SaoS-2 cells can also lead to a reduction in cell mass CV, we simulated the changes in cell mass CV during the G1 or nonG1 phase using values of *γ*^′^ and *m*_*τ*_′ obtained from the experimental data. When assuming a 20% CV for *α*^′^, growth rate modulation was found to decrease the cell mass CV in the G1 phase for U2OS and HT1080 cells (Fig. 4H), as well as in the nonG1 phase in HT1080 cells (Fig. 4I). However, it was not sufficient to reduce the cell mass CV in the nonG1 phase in SaoS-2 cells (Fig. 4I). As the CV of *α*^′^ increases, the reduction in cell mass CV becomes less pronounced (Fig. S12E-H). Eventually, all three cell lines failed to reduce cell mass CV at a 40% CV for *α*^′^ (Fig. S12G-H). Notably, despite U2OS G1 cells exhibiting a greatly negative *γ*^′^value, which indicated an exceptionally strong growth rate modulation, its effectiveness in reducing cell mass CV was lower than that of HT1080 G1 cells due to a smaller proportion of affected cells in U2OS, represented by a larger *m*_*τ*_′.

In summary, we found diverse patterns of correlation between cell mass and growth rate in different cell lines, and within the same cell line measured at different cell cycle stages. We developed stochastic models to explore the impact of different mass vs. growth correlations on the change in cell mass CV throughout the cell cycle. These models are representations of the data itself and not contrived schemes. They suggest strongly that in many cases sub-exponential growth, either for all cells or even for a subset of cells, can be an effective means of reducing cell mass CV and can ensure cell mass homeostasis (a stable mass variation over generations).

### Regulation of the cell cycle and regulation of growth rate compensate for each other to maintain cell mass homeostasis

Both size-dependent regulation of the progression through the cell cycle and size-dependent regulation of growth rate are used by cells to reduce cell mass variation. To evaluate the relative importance of these processes in maintaining cell mass homeostasis, we have tried to perturb each mechanism individually in RPE-1 cells.

To disrupt the size-dependent regulation of G1 length, we tried to slow the entry into S phase using palbociclib, an 6 inhibitor that specifically blocks the activation of Cdk4,6, which is required for entry into S phase(32). As discussed, low concentrations of palbociclib increased the mean cell mass and, as expected, prolonged the cell cycle length by elongating the G1 phase (Fig. 1H, Table S2). However, once treated cells had reached the new homeostatic state, the CV of birth mass remained unchanged compared to untreated cells (Fig. 1I). When we analyzed the duration of each cell cycle phase as a function of cell mass, we found a reduced impact of cell mass on G1 phase length coupled with an enhanced impact on S phase length, characterized by the slopes and the correlation coefficients of the correlations between cell mass and the durations of these phases (Fig. 5A-B, K). Additionally, the size-dependent regulation of G2 phase was diminished, yet still statistically significant (p = 0.0057) (Fig. 5C, K). These opposite changes in G1 and S phase regulation suggest that the size-dependent regulation of S phase had compensated for a weakened impact of cell mass on G1 length regulation. Hence, in specific circumstances such as palbociclib treatment, S phase can become the primary period responsible for reducing cell mass variation (Fig. 2G). Nevertheless, such compensation ultimately proves insufficient, resulting in a diminished cell mass-dependent regulation of the entire cell cycle length (Fig. 5D, K). To maintain the birth mass CV at the same level as untreated cells, additional regulation of growth rate is required to further reduce cell mass variation during the cell cycle. Indeed, we found that the correlations between cell mass and growth rate in palbociclib-treated cells were even closer to linear growth compared to untreated cells (Fig. 5E), implying a stronger growth rate modulation and a greater reduction in cell mass variation through growth rate regulation. The unchanged CV of birth mass when cells are treated with the G1/S inhibitor, palbociclib (Fig. 1I), is collectively a result of the interplay between size-dependent cell cycle regulation and size-dependent growth rate regulation. Thus the cell mass CV is maintained despite a significant increase in the mean birth mass (Fig. 1H).

**Figure 5.**
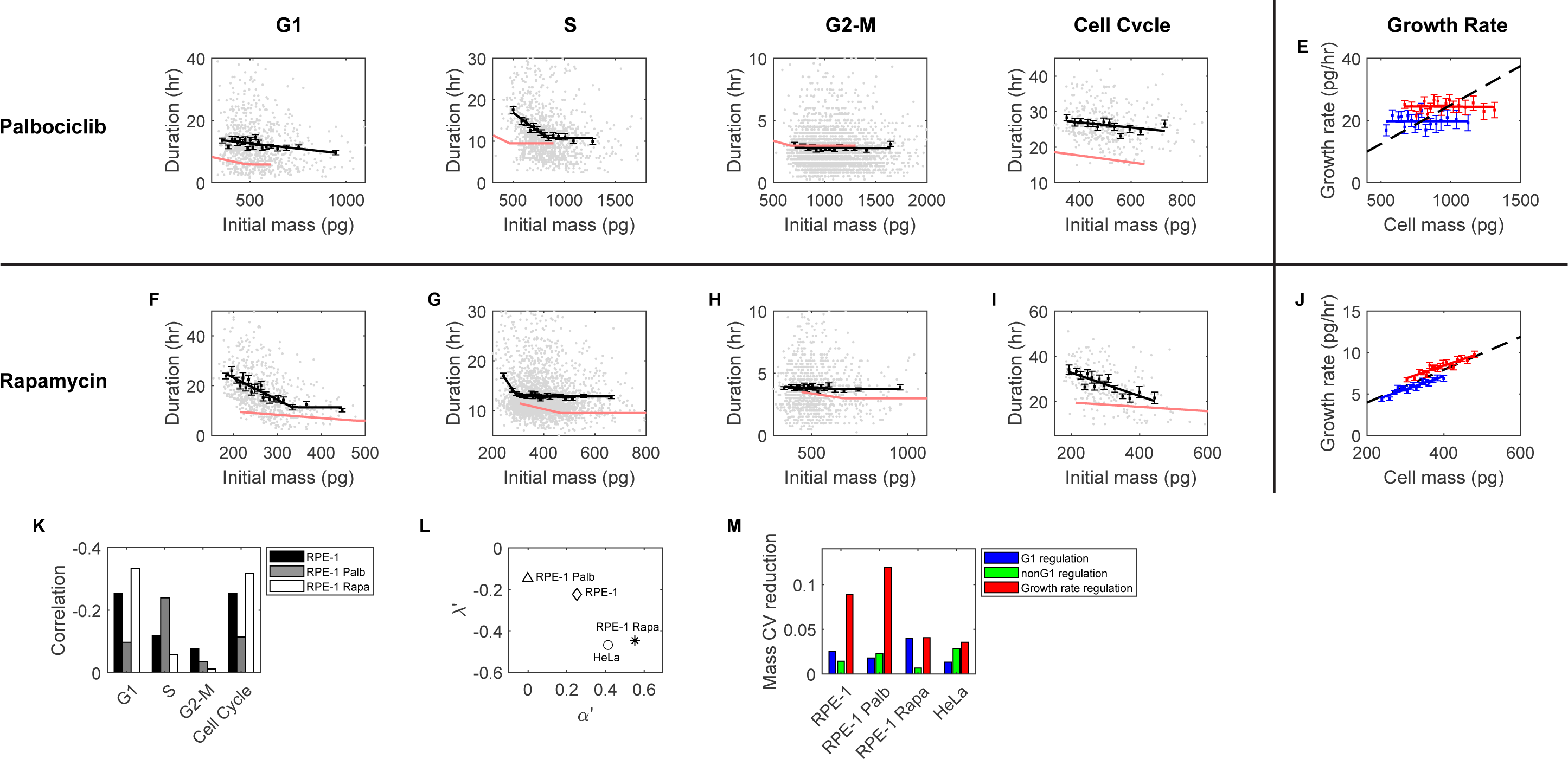
The compensatory roles of size-dependent cell cycle regulation and size-dependent growth rate regulation in maintaining cell mass homeostasis. (A-D) The correlation between G1 (A), S (B), G2-M (C), and full cell cycle (D) length and the initial mass of the corresponding period in RPE-1 cells treated with 50 nM palbociclib. Each gray dot is an observation; black squares indicate the average of each cell mass bin; error bars are SEM; solid black line is the best fit of the black squares; solid red lines are the corresponding correlations in untreated RPE-1 cells. (E) Correlations between cell mass and growth rate in the G1 (blue) and nonG1 (red) phases for RPE-1 cells treated with 50 nM palbociclib. Filled squares represent the median growth rate of each bin; error bars show SEM. The black dashed lines indicate the expected behavior for exponential growth. The solid blue and red lines are the best fit of the filled squares. (F-I) The correlation between G1 (F), S (G), G2-M (H), and full cell cycle (I) length and the initial mass of the corresponding period in RPE-1 cells treated with 100 nM rapamycin. (J) Correlations between cell mass and growth rate in the G1 (blue) and nonG1 (red) phases for RPE-1 cells treated with 100 nM rapamycin. (K) Kendall rank correlations between the duration of indicated cell cycle phase and cell mass at the initiation of the respective phase, in untreated RPE-1 cells, RPE-1 treated with 50 nM palbociclib, and RPE-1 treated with 100 nM rapamycin. (L) The correlation between the normalized slope of birth mass vs. cell cycle length correlation, *λ*′, and the normalized slope of cell mass vs. growth rate correlation, *α*′, depicted for untreated HeLa and RPE-1 cells, as well as RPE-1 cells treated with palbociclib or rapamycin. The values of *λ*′ and *α*′ used in this plot are listed in Table S7. (M) The contribution of each control mechanism shown as the reduction in the simulated division mass CV when the respective control mechanism is included compared to that without any control mechanisms. Simulation parameters were obtained from experimental data measured in untreated HeLa and RPE-1 cells, as well as RPE-1 cells treated with palbociclib or rapamycin.

In a converse experiment, we specifically perturbed cell growth rate. We treated cells with rapamycin to inhibit mTOR activity. Treatment with rapamycin resulted in an elongation of the cell cycle (Table S2) and a decrease in mean cell mass (Fig. 1H). Similar to the results with palbociclib treatment, rapamycin treatment left the birth mass CV unchanged (Fig. 1I). Cell mass-dependent feedback on G1 length was enhanced in the presence of rapamycin (Fig. 5F, K), while feedbacks on the S and G2-M phases were weakened (Fig. 5G-H, K). Additionally, the minimal lengths of all cell cycle phases were slightly increased compared to untreated cells (Fig. 5F-H). In the presence of rapamycin, the cell mass fed back more strongly on the entire cell cycle length, as indicated by the more negative slope and correlation coefficient of the mass vs. cell cycle length correlation (Fig. 5I, K). Furthermore, the relative strengths of correlations between cell mass and cell cycle phase length aligned with the reduced cell mass CV in the corresponding phases: for instance, cell mass CV was primarily reduced in the G1 phase with rapamycin treatment (Fig. 2H), consistent with the strongest cell cycle regulation in the G1 phase. On the other hand, we found that the slopes of the mass vs. growth correlations in both the G1 and nonG1 phases closely resembled that of exponential growth (Fig. 5J), suggesting a weaker role of growth rate regulation in maintaining cell mass homeostasis when growth rate is inhibited by rapamycin.

As should be clear from the experiments described above, size-dependent cell cycle regulation and size-dependent growth rate modulation interact with each other to maintain the birth mass CV at a consistent level even when the G1/S transition or cell growth rate is perturbed, resulting in significant changes in the mean birth mass. As we studied the feedbacks of cell mass on cell cycle length and growth rate in many different circumstances, we felt a need to develop new ways for comparing the response of each under different conditions. We have found it convenient to define a new parameter to represent the strength of this linkage. We utilized the normalized slope of birth mass vs. cell cycle length correlation as the parameter *λ*′, which quantifies the strength of *size-dependent cell cycle regulation*.

The value of *λ*^′^is always negative. A more negative value of *λ*^′^ indicates a stronger regulation. Additionally, since the slopes of the cell mass vs. growth rate correlations in the G1 and nonG1 phases were similar, we found it useful to calculate the average slope of these phases and normalized it by the mean doubling time to represent the strength of *size-dependent growth rate regulation*, which we denoted as *α*′. The value of *α*′ is smaller than or equal to *ln*2, which represents exponential growth. A smaller value of *α*′ indicates a greater deviation from exponential growth and thus a stronger modulation of growth rate. We found an inverse correlation between *λ*′ and *α*′ across all the conditions we have investigated (Fig. 5L, Table S7), suggesting a compensatory effect between the regulation of cell cycle and growth rate (i.e., the strengths of these regulatory processes tend to change in opposite directions). For instance, when cell cycle regulation was inhibited (e.g., by palbociclib), the modulation of growth rate became stronger, and conversely, when growth rate regulation was inhibited (e.g., by rapamycin), the modulation of cell cycle length became stronger. These findings highlight the compensatory roles played by these two processes in maintaining cell mass homeostasis.

To illustrate further the compensatory roles of regulation on cell cycle and growth rate, we developed a stochastic model to simulate changes in cell mass variation throughout the cell cycle (SI Text, Section 4). In this model, we considered three factors that could contribute to the increase of cell mass variation: variability in cell cycle length, variability in growth rate, and noise in cell mass partition during mitotic division. For simplicity, we only considered extrinsic noise as the source of growth rate variability, which is due to stochasticity in the partitioning of cellular contents during cell division, as previously discussed (Fig. S11C, SI Text, Section 3.1). As control mechanisms, we considered size-dependent regulation of the duration of G1 and nonG1 phases separately, and we also considered size-dependent growth modulation throughout the entire cell cycle. We chose all the parameters in this model from our actual experimental data and evaluated the impact of each control mechanism by comparing the cell mass CV at division with and without these control mechanisms. Notably, we observed some discrepancies between the simulated division mass CV, incorporating all three control mechanisms, and the values measured in experiments (Table S8). These may arise from the simplification of variability in growth rate (Fig. S11C, SI Text, Section 4), which effectively influences cell mass variation (Fig. S13) but is challenging to estimate accurately from experimental data. Nevertheless, these simulations largely reflect the relative significance of each control mechanism in maintaining cell mass homeostasis.

The model results indicate that in RPE-1 cells, the regulation of G1 length plays a slightly greater role compared to nonG1 length regulation, but both are overshadowed by the modulation of growth rate (Fig. 5M). When the G1/S control is inhibited by palbociclib, the contribution of G1 length regulation slightly decreases, the contribution of nonG1 regulation slightly increases, and the role of growth rate modulation becomes even more dominant (Fig. 5M). On the other hand, inhibiting growth with rapamycin leads to an increase in the dominance of G1 length regulation, with its contribution now comparable to that of growth rate modulation, while the impact of nonG1 regulation becomes smaller (Fig. 5M). In HeLa cells, the cell mass variation is considerably smaller than that in RPE-1 cells (Table S8) without considering any control mechanisms, due to the lower variation in growth rate in HeLa cells. It is worth noting that HeLa cells possess a mutated G1/S network. Its ranking of contributions from the three mechanisms is similar to the scenario observed in RPE-1 cells treated with palbociclib, which disrupts the G1/S transition. Specifically, in HeLa cells, the contribution of growth rate modulation outweighs that of nonG1 length regulation, which, in turn, outweighs that of G1 length regulation (Fig. 5M).

These findings collectively reveal compensatory roles of cell cycle and growth rate regulation in reducing cell mass variation, particularly distinguishing the regulation of G1 length and the regulation of growth rate. Generally, growth rate modulation is the more dominant mechanism. When one feedback process is hindered, other mechanisms become stronger to maintain cell mass variation at a similar level. Growth rate modulation, rather than cell cycle regulation, consistently plays the predominant role in reducing cell mass CV, regardless of whether or not the cells possess an intact G1/S circuit. Even when the growth rate is inhibited by rapamycin, the contribution of growth rate modulation remains on par with that of G1 length regulation. These observations contradict the conventional size control models (1,14,19,43,58–62), which predict that G1/S control is the primary contributor to size homeostasis in mammalian cells.

### Other explanations for how a population of cells might reduce its cell size variation

We explored additional processes that could potentially contribute to the reduction in cell mass CV but were not accounted for in our stochastic model. In principle, any process that affects the likelihood of cell division or cell viability differentially in large and small cells could influence the distribution of cell mass within a population. To estimate the importance of such effects, we examined the rate of cell death and cell cycle arrest through long-term measurements of cell growth and proliferation. During the 48 to 72-hour duration of our cell measurements, we defined cell cycle arrest events as instances where a cell remained in the same cell cycle phase while its mass continued to increase throughout the experiment. Furthermore, cell death was identified by a sudden and drastic decrease in cell dry mass, indicating cell membrane permeabilization.

We found events of cell cycle arrest or cell death in the culture affected no more than 2% of cells in all the conditions that were studied (Table S9). In particular, neither cell cycle arrest nor cell death occurred frequently enough to contribute significantly to cell mass homeostasis in any of the experiments that we have described. It is worth noting that the remarkably low frequency of cell cycle arrest in cells treated with rapamycin and palbociclib at the drug concentrations used in this study suggests that these drugs at low concentrations do not induce quiescence or senescence at the population level (Table S9).

Furthermore, the concentrations of these drugs did not appear to be toxic enough to cause significant cell death (Table S9). One intriguing observation was that some large RPE-1 cells treated with palbociclib experienced a partial loss of cytoplasm during mitosis (Table S9, Movie S1). This cytoplasmic loss could be attributed to incomplete cortical contraction during mitotic rounding(63). The amount of mass loss appeared to be random. Notably, these rare events, accounting for approximately 0.5% of cells, did not have a significant impact at the population level on cell mass homeostasis in the presence of palbociclib.

It is worth noting that although these mechanisms were of negligible importance in the specific experimental setting of our study, they might still play a significant role in a tissue setting, for example during wound healing, regeneration, aging, and/or disease.

### A picture of cell mass homeostasis in proliferating cells

Homeostasis refers to a process of maintaining a balance between the inherent noise in cellular processes and the feedback control mechanisms that correct for them. In proliferating cells, this noise arises from stochastic variation in growth rate, cell cycle length, and cell mass partitioning during mitosis. To reduce cell mass variation, size-dependent regulation can occur through the control of cell cycle progression, growth rate, or both.

To illustrate size regulation graphically as a balance between noise and control mechanisms, we have depicted the concept of cell mass homeostasis as a “teeter-totter” (Fig. 6). Stochastic noise and feedback control mechanisms are represented as opposing forces on either side of the lever’s fulcrum; the size of the icon represents the importance of the process, as determined from the stochastic models (Table S8). When these effects are balanced, the system reaches a steady state. In cell lines like RPE-1, where the G1/S circuit is intact, the relative importance of the control mechanisms can be ranked from greatest (heaviest on the teeter-totter) to smallest (lightest on the teeter-totter) as follows: growth rate modulation, G1 length regulation, and nonG1 length regulation. A perturbation of the system leads to changes in the stochastic nature of the processes and affects the operation of specific control mechanisms. When this happens, other control mechanisms compensate for these changes and restore the balance. For example, when G1/S control is inhibited, either through pharmacological inhibitors, such as palbociclib, or genetic mutations in the G1/S circuitry, as seen in HeLa cells, the contribution of G1 length regulation is reduced. In response, nonG1 length regulation and growth rate modulation become more significant. Conversely, when growth rate modulation is inhibited, such as by rapamycin, G1 length regulation becomes more important, and growth rate modulation contributes less.

**Figure 6.**
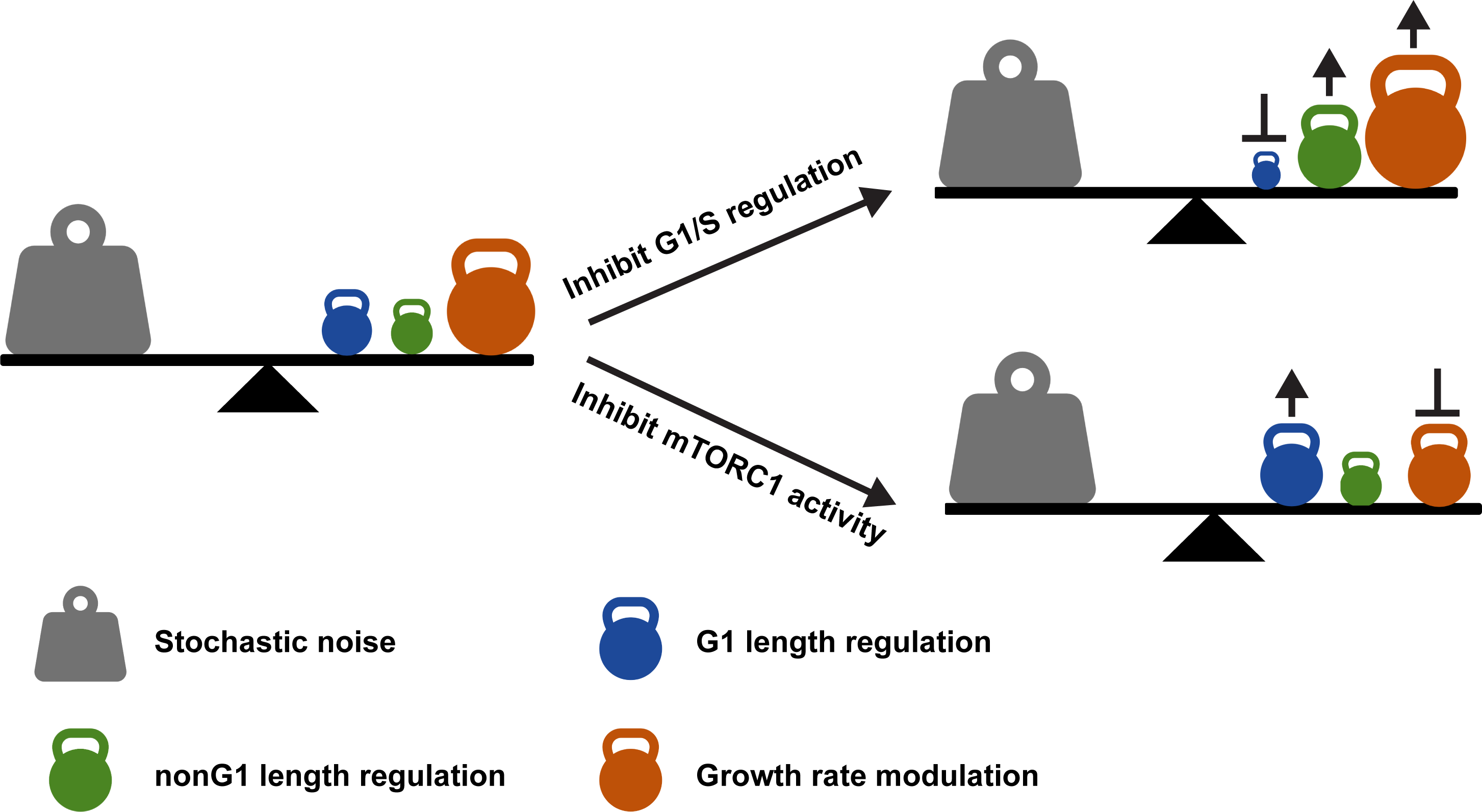
The teeter-totter model of cell mass homeostasis. Cell mass homeostasis requires a balance between stochastic noise and control mechanisms. In unperturbed cells with an intact G1/S circuitry, the weights of control mechanisms from the heaviest to the lightest are the growth rate modulation, G1 length regulation, and nonG1 length regulation. When G1/S control is perturbed, the impact of the G1 length regulation becomes smaller, and the nonG1 length regulation and growth rate modulation become larger to compensate. When the growth rate modulation is suppressed, the G1 length regulation plays a more prominent role in compensating for the reduced impact of growth rate modulation.

Overall, the teeter-totter of cell mass homeostasis is robustly balanced through the compensatory interactions of these different control processes within the cell. It is likely that the coordination and adjustment of these compensatory mechanisms at the molecular level are crucial for cellular survival under changing conditions. While our understanding of how these mechanisms achieve balance has advanced, further study is needed to elucidate how they coordinate and adapt their compensation at the molecular level to maintain balance under the changed conditions and how this plays out in health, disease, aging, etc.

## Discussion

To summarize: in examining cell mass homeostasis, we found that stochastic variation in cell mass in proliferating cells is tightly controlled *throughout the cell cycle* (Fig. 2) via size-dependent regulation of cell growth rate (Fig. 4) and size-dependent regulation of cell cycle progression (Fig. 3). Generally speaking, among the cell lines and cell cycle and cell growth inhibitors that we have employed (including those previously studied and analyzed), we conclude that the G1/S transition does not appear to be a privileged place where cell size regulation is imposed. Rather size regulation occurs throughout the cell cycle phases. The compensation that keeps stochastic variation of size in check emerges from an interplay of these mechanisms and results in effective cell mass regulation. Not only is homeostasis maintained, but it is also maintained at high stringency, as indicated by the narrow distribution of cell mass at birth (Fig. 1). Furthermore, cell mass homeostasis is robust to changes in genetic background and is resistant to manipulations of the G1/S transition or perturbation of mTOR activity (Fig. 1). The birth mass CV measured in many proliferating bacterial, yeast, mammalian, and plant cells falls in a relatively small range (from 11% to 25%) (Table S10), which is comparable to the birth weight CV of a human fetus(64). Although it is not clear whether such strict control is explicitly selected for during evolution or merely a by-product of some other selection (65,66), cell mass homeostasis appears to be highly regulated and presumably important. Though we focused on cultured human cell lines in this study, the mechanisms underlying cell size homeostasis, just as the mechanisms underlying the cell cycle itself, are likely to be conserved.

In this study, we utilized ceQPM(28) as a means of measuring cell dry mass, providing a complementary approach to previous studies that focused on cell volume as an indicator of cell size(3,7). We found that many aspects of the behavior of cell mass, as directly measured by ceQPM, were consistent with studies of cell volume, particularly those reported by Cadart et al., who obtained high-quality cell volume data(7). For example, in line with their observations, we also identified inverse correlations between initial mass and cell cycle phase duration in both the G1 and nonG1 phases in HeLa cells (Fig. 3), the existence of a minimal duration of the G1 phase (Fig. 3), the “adder”-like correlation between the birth and division masses (Fig. 3), and the coordination between size-dependent cell cycle regulation and growth rate modulation in maintaining cell size homeostasis (Fig. 5). This consistency is further supported by our recent findings that cell volume usually changes proportionally with cell mass in cultured proliferating cells, except during mitosis, resulting in a narrow distribution of cell mass density(67). However, we were able to observe more detailed discrepancies in the regulation of mass and volume growth. For instance, while Cadart et al. reported that volume growth rate is dependent on cell volume at birth(7), we found that mass growth rate is related to cell mass at any point of the cell cycle, and this relationship varies across different cell cycle stages (Fig. 4, S8). Moreover, the noise in mass growth rate appears to affect the slope of the correlation (Fig. S11), in contrast to Cadart et al.’s findings of noise primarily impacting the intercept in volume growth rate(68). These discrepancies may be attributed to inherent differences in the factors affecting mass or volume and the speed and mechanisms by which cells respond to perturbations or fluctuations in mass or volume(69).

Aside from confirming previous discoveries, our findings took a significant step forward in exploring mechanisms underlying cell mass homeostasis. Extensive data collection on large populations of cells was possible thanks to the high throughput of ceQPM(28). From these extensive measurements, we derived reliable correlations between cell mass, the duration of cell cycle phases, and the growth rate. We studied these across multiple cell lines and under various pharmacologic perturbations. We were able to fit such data to simple functions (Fig. 3, 4, 5), which facilitated our ability to derive quantitative models. These models, in turn, facilitated our interpretation of the underlying cellular responses. For example, we showed how G1, S, and G2 phases are each under negative regulation by cell mass in both transformed and non-transformed cells (Fig. 3). A particularly noteworthy discovery was the identification of a minimum length for each phase of the cell cycle in large cells, which explains the limited impact of cell cycle regulation on very large cells, leaving the underlying process to growth rate modulation. (Fig. 3). We further demonstrated that growth rate is modulated differently in different cell types or cell lines (Fig. 4). Such comprehensive characterization of growth regulation was not previously possible without the extensive and precise measurements of cell mass and growth rate by ceQPM(28). When we perturbed cells by inhibiting the G1/S transition or suppressing the growth rate (Fig. 5), ceQPM enabled us to go beyond the qualitative phenomena observed in previous studies (8,11,12). It allowed us not only to determine the average changes in cell size, cell cycle phase duration, and growth rate but also to measure these qualities at the single-cell level, tracking the individual cells over time. This enabled us to derive important quantitative correlation functions. These functions in turn allowed us to write deterministic equations, incorporate stochastic noise, and ultimately develop a stochastic model. With this model, we could estimate the relative weight of each of the regulatory mechanisms employed in maintaining cell mass homeostasis and finally deduce how the weights of these separate mechanisms depend on each other (Fig. 5).

One simple finding stands out. It has been generally assumed, and widely cited in review articles and textbooks of biology, that G1 length regulation is the predominant or even the sole mechanism controlling cell size during the cell cycle (1,14,19,43,58–62). There was always an appeal of this simple mechanism, as it made perturbation of the cell cycle at G1/S the whole process for cell mass control. This is clearly not the case. Our current highly quantitative studies involving at least hundreds of cells per condition demonstrated unequivocally that, at least for the cell lines we employed, the impact of G1 length regulation on constraining cell mass CV within a proliferating cell population is much less significant than the modulation of cells’ mass accumulation (growth) rate (Fig. 5). This holds true for non-transformed cells with intact G1/S control. Furthermore, even in the presence of growth inhibition induced by rapamycin, the contribution of growth rate modulation to cell mass CV reduction is no less than that of G1 length regulation.

Why would there be size-dependent growth rate regulation if regulation of cell cycle progression were sufficient to control cell size? With so many essential genes in the genome, it seems like a weak argument to claim that having two separate mechanisms provides increased security for survival. We propose instead that they serve two separate functions. Control of G1/S might be used primarily to set the cell size for a given cell *type*. In this view, the G1/S transition is hard-wired into developmental pathways like the MAP kinase pathway or the BMP pathway through proteins like TGFβ. By contrast, control of cell growth might be primarily used for a different purpose: *maintaining* cell size homeostasis of any given cell type against environmental or stochastic variation. It makes more sense that the targeted *mean* size of a given cell type is controlled by a few key molecular players downstream of specific hormonal or nutrient signals or cellular differentiation. Those molecular players (such as CDK4/6 or other CDK inhibitors) were described as a cell size “dial” in a previous model by Tan et al.(11).

However, once cells are programmed to adopt a defined size in their new state, they would still require a mechanism to maintain size homeostasis around that new mean by buffering against environmental or internal stochastic fluctuation. Consistent with the work presented here (Fig. 1) and studies in budding and fission yeasts(13,17), deletion or overexpression of the G1/S inhibitors change the mean size dramatically but have only limited effects on the variation of cell size. Furthermore, systems that only act at a single gate for size variation would fail to provide continuous feedback on size variation and would have difficulty correcting noise introduced after that gate operates, which in this case is early in the cell cycle (70). By contrast, growth rate regulation, particularly sub-exponential growth, where growth rate is proportional to cell mass but exhibits a slope smaller than that of exponential growth, proves to be a highly effective means for reducing cell mass variation throughout the cell cycle (Fig. 4, SI Text, Section 3). The effectiveness of this mechanism is bolstered by its operation throughout the entire cell cycle and in all cells. This form of regulation would be more effective than growth rate modulation restricted to short periods of the cell cycle and only in large cells, as suggested by previous studies(1,8,20,37,49). Unraveling the determinant factors that underlie the sub-exponential scaling between growth rate and cell mass will likely shed light on the coordination between size-dependent biomass synthesis, nutrient transportation, and macromolecule destruction(71). We can imagine that pathological conditions, such as aging related diseases, may target growth rate regulation and therefore affect cells at different stages of cell cycle or even non-growing cells.

Aside from the size-dependent regulation on the G1 length and cell growth rate, the regulation of nonG1 phase lengths also contributes significantly to the reduction of cell mass variation (Fig. 5). This is presumably due to the fact that cell cycle phases outside of G1 have non-negligible negative correlations with cell mass (Fig. 3) and often occupy a larger portion of the cell cycle than the G1 phase (Table S2). The mechanisms regulating G2 length have been mainly studied in fission yeast, where the G2/M transition acts as the major size control checkpoint(17,72–74). Mammalian cells share homologous components of this G2/M regulation with fission yeast(75,76), suggesting that similar mechanisms might function during this stage in mammalian cells. However, further investigation beyond simple homology will be needed to confirm this possibility. The regulation of S phase length as a means for controlling cell mass in mammalian cells has rarely been explored. One potential mechanism of size-dependent S phase length regulation could involve the control of the number of replication complexes. If the number of forks were proportional to the total cell mass, so that small cells made fewer forks, this could lengthen S phase (77). If cell mass were to affect the number of active origins or DNA replication speed, it might also affect the level of DNA damage due to the under-replicated regions(77–79). Replication stress is not uncommon in normal cycling populations, as evidenced by the presence of DNA lesions in more than 20% of G1 cells in non-transformed cell lines (80). If the occurrence of replication stress is influenced by cell size and leads to a form of DNA damage that cannot be resolved, it could potentially drive tumorigenesis or senescence in a cell size-dependent manner, resulting in heterogeneous behavior in a genetically uniform population. This scenario might hold clinical significance and thus deserves further investigation. Additional research may be needed to establish the relationship between the probability of replication stress and cell mass during S phase. Furthermore, the actual mechanism of S phase length regulation could be more complicated than the size-dependent replication fork number. The negative correlation between cell mass and S phase length is strengthened in palbociclib-treated RPE-1 cells compared to untreated cells (Fig. 5), suggesting more complex crosstalk between the G1 and S phase regulation that cannot be fully explained by the size-dependent replication fork number.

The data presented here makes the case that regulation of cell mass, which occurs throughout the entire cell cycle, is achieved by continuously monitoring cell size and adjusting the rate of mass accumulation. Such a mechanism would require that a cell continually “know” how large it is and how large it should be. How might cells sense their size relative to a changing standard that changes with cell types, nutrient conditions, and pharmacological perturbations? A proposed mechanism of cell size sensing relies on some form of disproportionality of molecular components or signals to size. For instance, cells might sense size through the sub-scaling of inhibitors or super-scaling of activators to regulate their cell cycle length(6,10,43,81). Cell mass accumulation requires nutrient provision, transcription, translation, and degradation; any rate-limiting step might serve as a size sensor. It has also been proposed that cells may sense size and modulate growth rate by DNA limitation, cytoplasmic dilution, surface-to-volume ratio, sublinear proportionality between metabolic rate and cell mass, transport efficiency, and other such mechanisms(20,70,82–84). We have found that different cell lines modulate growth rates in different manners. It is of course plausible that each cell line we investigated employs a distinct size-sensing mechanism and a distinct mode of response of mass accumulation.

However, it is more likely that all cell lines share a universal mechanism that allows various forms of growth rate modulation under particular conditions. One potential candidate for this universal mechanism would be the mTOR pathway, which governs biomass synthesis and responds to various upstream signals(35,85). Investigating how the mTOR pathway responds to cell size warrants further exploration. Additionally, growth rate regulation exhibits cell cycle-specific patterns and even intrinsic oscillations (28,48,50,51). The coexistence of multiple forms of regulation complicates any investigation. It would be helpful if we could isolate each mechanism for study, perhaps by identifying conditions where only one of the processes is dominant. Situations such as cell cycle arrest and size enlargement (so called cellular senescence) triggered by DNA damage or other stresses are of particular interest(43,86), which may disentangle size-dependent growth regulation from the cell cycle-dependent growth regulation allowing us to focus on the effects of cell size on growth rate using the methods we employed in this study.

In summary, the use of ceQPM to quantify single-cell dry mass, mass growth rate, and cell cycle progression has provided, as yet, the most accurate, complete, and quantitative description of cell mass homeostasis in mammalian cells. In a sense we have in this paper showcased the seldomly appreciated power of phenomenological descriptions. Such descriptions have been proven to be inherently powerful in physics and chemistry. The simple and unequivocal phenomenon of the reduction in the coefficient variation of cell mass within a proliferating population *throughout* the cell cycle, itself directly ruled out the possibility that cells control mass solely or principally at the G1/S transition by controlling the length of the G1 phase. While this result is far from a complete answer to the problem of cell size homeostasis and does not yet provide specific molecular mechanisms, it nevertheless serves as a guide for future investigation. It redirected our focus away from the G1/S transition in cell size homeostasis. Our next step will concentrate on understanding the molecular-level mechanisms governing size-dependent regulation of growth rate, as this aspect, being the predominant player, holds greater promise in elucidating how cells maintain a stable size distribution. The compensatory responses to perturbing size suggest that there are regulatory pathways in cell size regulation that we have not appreciated. It suggest that we need to explore how cell size feeds back on the anabolic or proteostatic machinery, thus posing the next big questions to address in our ongoing research.

As of now, we are still in the early stage of defining the phenomenon of size homeostasis and have much to learn about the regulatory circuits that tell a cell how large it is and how large it should be at any given time or in any given circumstance. Studying cell size homeostasis in cultured cells can lay the groundwork for future investigations into size control in vivo and its implications for disease, thereby expanding our understanding of cell physiology.

## Materials and Methods

### Cell Culture and Chemical Treatment

HeLa mAG-hGem, RPE-1 mAG-hGem, HT1080 mAG-hGem mKO2-hCdt1, and HeLa mAG-hGEM DNA-ligase-dsRed cells were made in previous studies by our laboratory(37,87). U2OS mAG-hGem and Saos-2 mAG-hGEM cells were generated by lentivirus infection in this study. Lentivirus carrying mTurquoise2-SLBP was purchased from Addgene (83842-LV) to make HeLa mAG-hGem mTurquoise2-SLBP, RPE-1 mAG-hGem mTurquoise2-SLBP, and HeLa mAG-hGEM DNA-ligase-dsRed mTurquoise2-SLBP. Single clones of stable expression were selected for each cell line. Cells were incubated at 37 °C with 5% CO2 in Dulbecco’s Modified Eagle’s Medium (DMEM) (11965; Thermo Fisher Scientific) with 25 mM HEPES (15630080; Thermo Fisher Scientific) and 10 mM sodium pyruvate (11360070; Thermo Fisher Scientific), or McCoy’s 5A Medium (16600082; Thermo Fisher Scientific). Both media were supplemented with 10% fetal bovine serum (FBS) (16000044; Thermo Fisher Scientific) and 1% penicillin/streptomycin (15140122; Thermo Fisher Scientific). Palbociclib and MG132 were purchased from Selleckchem (PD-0332991 and S2619), rapamycin was purchased from LC Laboratories (R-5000), and cycloheximide was purchased from Millipore Sigma (C4859).

### Live Cell Imaging

Cells were imaged at 10X magnification by an Eclipse Ti microscope with the Perfect Focus System (PFS) (Nikon, Japan) and an SID4BIO camera (Phasics, France). Nikon NIS-Elements AR ver. 4.13.0.1 software with the WellPlate plugin was used to acquire images. A home-made incubation chamber was used to maintain a constant environment of 36 °C and 5% CO2 during imaging. Cells were seeded on 6 well glass bottom plates (P06G-1.5-14-F; MatTek) at a density of 1500 cells/cm^2^ 3 h before long-term imaging or 3500 cells/cm^2^ 16 h before short-term imaging. Before time-lapse imaging was started, mineral oil (M8410; Millipore Sigma) was added into each well to prevent media evaporation. In the long-term experiments studying the cell cycle regulation, cells were monitored for 48 or 72 h. In the short-term experiments studying growth rate modulation, cells were monitored for 3 h. For all experiments, the phase images were acquired every 30 min, and the fluorescence images were acquired every 1 h.

### Cell fixation and cell cycle identification

After the short-term time-lapse imaging, the mineral oil was gently removed by aspiration. Cells were fixed with 4% paraformaldehyde (RT 157-8; Electron Microscopy Sciences) and stained with Hoechst 33342 (62249; Thermo Fisher Scientific) at a final concentration of 1 µM. The cells were then imaged by QPM again to identify their cell cycle stages.

### QPM Image Processing and Data Analysis

The QPM images were processed by the ceQPM method developed previously(28) and conducted on the O2 high-performance computing cluster at Harvard Medical School.

To test the significance of the minimal cell cycle phase length, we fitted the binned correlations between the initial mass and cell cycle phase duration in Fig. 3 and Fig. 5 with two alternative models. a linear model *y* = *a*_1_*x* + *b*_1_, and a bilinear model *y* = *a*_2_*x* + *b*_2_ (*x* ≤ *x*_0_), *y* = *a*_2_*x*_0_ + *b*_2_ (*x* > *x*_0_), where *y* is the cell cycle phase length, *x* is the initial mass, *a*_1_, *b*_1_, *a*_2_, *b*_2_, and *x*_0_ are the fitting parameters. We used the Akaike Information Criterion (AIC) to compare the goodness of fits. A smaller AIC indicates a better fit, and the relative likelihood *p*_*linear*_ or *p*_*bilinear*_ predicts the probability that the alternative model is a better fit when the linear or bilinear model has the smaller AIC (88). Since the correlations between the initial mass and cell cycle phase duration were not linear, we utilized the Kendall’s rank correlation coefficient to represent the correlation strength. This coefficient is more suitable for our data as it does not assume a linear relationship, unlike the widely used Pearson correlation coefficient(89).

To evaluate whether the cell cycle control could explain the adder behavior in Fig. 3I-J, we assumed cells grew exponentially at the rate of *α* = ln(2) /*DT*, where *DT* is the averaged cell cycle length. The division mass could be predicted by *m*_*d*_ = *m*_*b*_*e*^*αT*^, *T* = *f*(*m*_*b*_), where *f* is the best fitted function in the alternative models of cell cycle length vs. birth mass.

To fit the binned correlation between growth rate and cell mass in Fig. 4 and Fig. 5, we employed two alternative models: a linear model 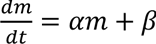 and a bilinear model 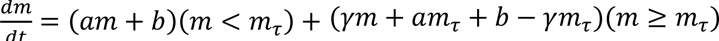, where *y* is the growth rate, *x* is the cell mass, *α*, *β*, *a*, *b*, *γ*, and *m*_*τ*_ are the fitting parameters. We used the Akaike Information Criterion (AIC) to estimate the goodnesses of fits.

## Data Availability

Cell lines are available upon request. All data are included in the manuscript and supporting source data files.

## Acknowledgements

We thank the National Institute of General Medical Sciences (5RO1GM26875-42, 5R35GM145248) and National Institute on Aging (1R56AG073341, 5R01AG073341) for grant support. We thank the Nikon Imaging Center at Harvard Medical School for sharing its resources. We thank the Research Computing Group at Harvard Medical School for providing support for the O2 Computing Cluster for imaging processing and data storage. We thank Johan Paulsson, Ariel Amir, Prathitha Kar, Ethan Levien, Ahmed Rattani, Wenzhe Ma, Gabriel Neurohr, Simon Gemble, and Renata Basto for insightful suggestions. We thank William Ratzan for proof-reading the manuscript.

## References

1. Ginzberg MB, Kafri R, Kirschner M. On being the right (cell) size. Science [Internet]. 2015;348(6236):1245075–1245075. Available from: http://www.sciencemag.org/cgi/doi/10.1126/science.1245075

2. Killander D, Zetterberg a. A quantitative cytochemical investigation of the relationship between cell mass and initiation of DNA synthesis in mouse fibroblasts in vitro. Exp Cell Res [Internet]. 1965 Oct;40(1):12–20. Available from: http://www.ncbi.nlm.nih.gov/pubmed/5838935

3. Varsano G, Wang Y, Wu M, Varsano G, Wang Y, Wu M. Probing Mammalian Cell Size Homeostasis by Channel-Assisted Cell Reshaping. CellReports [Internet]. 2017;20(2):397–410. Available from: 10.1016/j.celrep.2017.06.057

4. Xie S, Skotheim JM. A G1 Sizer Coordinates Growth and Division in the Mouse Epidermis. Curr Biol [Internet]. 2020;30(5):916–924.e2. Available from: 10.1016/j.cub.2019.12.062

5. Dolznig H, Grebien F, Sauer T, Beug H, Müllner EW. Evidence for a size-sensing mechanism in animal cells. Nat Cell Biol [Internet]. 2004 Sep [cited 2012 Dec 5];6(9):899–905. Available from: http://www.ncbi.nlm.nih.gov/pubmed/15322555

6. Zatulovskiy E, Zhang S, Berenson DF, Topacio BR, Skotheim JM. Cell growth dilutes the cell cycle inhibitor Rb to trigger cell division. Science (80-) [Internet]. 2020;369(6502):466–71. Available from: http://science.sciencemag.org/

7. Cadart C, Monnier S, Grilli J, Sáez PJ, Srivastava N, Attia R, et al. Size control in mammalian cells involves modulation of both growth rate and cell cycle duration. Nat Commun [Internet]. 2018;9(1). Available from: 10.1038/s41467-018-05393-0

8. Ginzberg MB, Chang N, Kafri R, Kirschner MW. Cell size sensing in animal cells coordinates anabolic growth rates and cell cycle progression to maintain cell size uniformity. Elife [Internet]. 2018;(1):123851+. Available from: 10.1101/123851

9. Zhang S, Zatulovskiy E, Arand J, Sage J, Skotheim JM. The cell cycle inhibitor RB is diluted in G1 and contributes to controlling cell size in the mouse liver. Front Cell Dev Biol [Internet]. 2022 Aug 25;10. Available from: https://www.frontiersin.org/articles/10.3389/fcell.2022.965595/full

10. Lanz MC, Zatulovskiy E, Swaffer MP, Zhang L, Ilerten I, Zhang S, et al. Increasing cell size remodels the proteome and promotes senescence. Mol Cell [Internet]. 2022 Sep;82(17):3255–3269.e8. Available from: https://linkinghub.elsevier.com/retrieve/pii/S1097276522007134

11. Tan C, Ginzberg MB, Webster R, Iyengar S, Liu S, Papadopoli D, et al. Cell size homeostasis is maintained by CDK4-dependent activation of p38 MAPK. Dev Cell [Internet]. 2021 May;1–14. Available from: 10.1016/j.devcel.2021.04.030

12. Liu S, Ginzberg MB, Patel N, Hild M, Leung B, Li Z, et al. Size uniformity of animal cells is actively maintained by a p38 MAPK-dependent regulation of G1-length. Elife [Internet]. 2018 Mar 29;7(April):1–27. Available from: https://elifesciences.org/articles/26947

13. Chen Y, Zhao G, Zahumensky J, Honey S, Futcher B. Differential Scaling of Gene Expression with Cell Size May Explain Size Control in Budding Yeast. Mol Cell [Internet]. 2020;78(2):359–370.e6. Available from: 10.1016/j.molcel.2020.03.012

14. Zetterberg A, Larsson O. Kinetic analysis of regulatory events in G1 leading to proliferation or quiescence of Swiss 3T3 cells. Proc Natl Acad Sci U S A [Internet]. 1985 Aug 1;82(16):5365–9. Available from: http://www.pubmedcentral.nih.gov/articlerender.fcgi?artid=390569&tool=pmcentrez&renderty pe=abstract

15. Araujo AR, Gelens L, Sheriff RSM, Santos SDM. Positive Feedback Keeps Duration of Mitosis Temporally Insulated from Upstream Cell-Cycle Events. Mol Cell [Internet]. 2016;64(2):362–75. Available from: 10.1016/j.molcel.2016.09.018

16. Garmendia-torres C, Tassy O, Matifas A. Multiple inputs ensure yeast cell size homeostasis during cell cycle progression. 2018;1–27.

17. Sveiczer a, Novak B, Mitchison JM. The size control of fission yeast revisited. J Cell Sci [Internet]. 1996 Dec;109 (Pt 1:2947–57. Available from: http://www.ncbi.nlm.nih.gov/pubmed/9013342

18. Turner JJ, Ewald JC, Skotheim JM. Cell Size Control in Yeast. Curr Biol [Internet]. 2012 May [cited 2012 May 7];22(9):R350–9. Available from: http://linkinghub.elsevier.com/retrieve/pii/S0960982212001923

19. Zatulovskiy E, Skotheim JM. On the Molecular Mechanisms Regulating Animal Cell Size Homeostasis. Trends Genet [Internet]. 2020;36(5):360–72. Available from: 10.1016/j.tig.2020.01.011

20. Neurohr GE, Terry RL, Lengefeld J, Bonney M, Brittingham GP, Moretto F, et al. Excessive Cell Growth Causes Cytoplasm Dilution And Contributes to Senescence. Cell [Internet]. 2019 Feb;176(5):1083–1097.e18. Available from: http://www.ncbi.nlm.nih.gov/pubmed/30739799

21. Si F, Le Treut G, Sauls JT, Vadia S, Levin PA, Jun S. Mechanistic Origin of Cell-Size Control and Homeostasis in Bacteria. Curr Biol [Internet]. 2019;29(11):1760–1770.e7. Available from: 10.1016/j.cub.2019.04.062

22. Zlotek-Zlotkiewicz E, Monnier S, Cappello G, Le Berre M, Piel M. Optical volume and mass measurements show that mammalian cells swell during mitosis. J Cell Biol [Internet]. 2015;211(4):765–74. Available from: http://jcb.rupress.org/content/211/4/765.abstract

23. Cooper KL, Oh S, Sung Y, Dasari RR, Kirschner MW, Tabin CJ. Multiple phases of chondrocyte enlargement underlie differences in skeletal proportions. Nature [Internet]. 2013 Mar 13 [cited 2013 Mar 14];495(7441):375–8. Available from: http://www.nature.com/doifinder/10.1038/nature11940

24. Son S, Kang JH, Oh S, Kirschner MW, Mitchison TJ, Manalis S. Resonant microchannel volume and mass measurements show that suspended cells swell during mitosis. J Cell Biol [Internet]. 2015;211(4):757–63. Available from: http://www.jcb.org/cgi/doi/10.1083/jcb.201505058

25. Venkova L, Vishen AS, Lembo S, Srivastava N, Duchamp B, Ruppel A, et al. A mechano-osmotic feedback couples cell volume to the rate of cell deformation. Elife [Internet]. 2022;11:2021.06.08.447538. Available from: 10.1101/2021.06.08.447538%0Ahttps://www.biorxiv.org/content/10.1101/2021.06.08.447538v2%0Ahttps://www.biorxiv.org/content/10.1101/2021.06.08.447538v2.abstract

26. Zangle TA, Teitell MA. Live-cell mass profiling: an emerging approach in quantitative biophysics. Nat Methods [Internet]. 2014;11(12):1221–8. Available from: http://www.nature.com/doifinder/10.1038/nmeth.3175

27. Popescu G, Park K, Mir M, Bashir R. New technologies for measuring single cell mass. Lab Chip. 2014;14(4):646–52.

28. Liu X, Oh S, Peshkin L, Kirschner MW. Computationally enhanced quantitative phase microscopy reveals autonomous oscillations in mammalian cell growth. Proc Natl Acad Sci [Internet]. 2020 Nov 3;117(44):27388–99. Available from: http://www.pnas.org/lookup/doi/10.1073/pnas.2002152117

29. Huh D, Paulsson J. Random partitioning of molecules at cell division. Proc Natl Acad Sci. 2011;108(36):15004–9.

30. Scott SJ, Suvarna KS, D’Avino PP. Synchronization of human retinal pigment ephitilial-1 (RPE-1) cells in mitosis. J Cell Sci [Internet]. 2020 Jan 1; Available from: https://journals.biologists.com/jcs/article/doi/10.1242/jcs.247940/266615/Synchronization-of-human-retinal-pigment

31. Sung Y, Tzur A, Oh S, Choi W, Li V, Dasari RR, et al. Size homeostasis in adherent cells studied by synthetic phase microscopy. Proc Natl Acad Sci U S A [Internet]. 2013 Oct 8 [cited 2014 Mar 21];110(41):16687–92. Available from: http://www.ncbi.nlm.nih.gov/pubmed/24065823

32. Fry DW, Harvey PJ, Keller PR, Elliott WL, Meade M, Trachet E, et al. Specific inhibition of cyclin-dependent kinase 4/6 by PD 0332991 and associated antitumor activity in human tumor xenografts. Mol Cancer Ther [Internet]. 2004 Nov;3(11):1427–38. Available from: http://www.ncbi.nlm.nih.gov/pubmed/15542782

33. Morgan D. The cell cycle: principles of control. Oxford Univesity Press; 2007.

34. Li J, Kim SG, Blenis J. Rapamycin: One drug, many effects. Cell Metab [Internet]. 2014;19(3):373–9. Available from: 10.1016/j.cmet.2014.01.001

35. Saxton RA, Sabatini DM. mTOR Signaling in Growth, Metabolism, and Disease. Vol. 168, Cell. Cell Press; 2017. p. 960–76.

36. Sakaue-Sawano A, Kurokawa H, Morimura T, Hanyu A, Hama H, Osawa H, et al. Visualizing spatiotemporal dynamics of multicellular cell-cycle progression. Cell [Internet]. 2008 Feb 8 [cited 2013 Feb 10];132(3):487–98. Available from: http://www.ncbi.nlm.nih.gov/pubmed/18267078

37. Kafri R, Levy J, Ginzberg MB, Oh S, Lahav G, Kirschner MW. Dynamics extracted from fixed cells reveal feedback linking cell growth to cell cycle. Nature [Internet]. 2013 Feb 27 [cited 2013 Feb 27];494(7438):480–3. Available from: http://www.nature.com/doifinder/10.1038/nature11897

38. Bajar BT, Lam AJ, Badiee RK, Oh YH, Chu J, Zhou XX, et al. Fluorescent indicators for simultaneous reporting of all four cell cycle phases. Nat Methods. 2016;13(12):993–6.

39. McGarry TJ, Kirschner MW. Geminin, an inhibitor of DNA replication, is degraded during mitosis. Cell. 1998;93(6):1043–53.

40. Whitfield M, Zheng L, Baldwin A, Ohta T, Hurt M, Marzluff W. Stem-loop binding protein, the protein that binds the 3′ end of histone mRNA, is cell cycle regulated by both translational and posttranslational mechanisms. … Cell Biol [Internet]. 2000 [cited 2014 Jan 15];20:4188–98. Available from: http://mcb.asm.org/content/20/12/4188.short

41. Leonhardt H, Rahn H-P, Weinzierl P, Sporbert A, Cremer T, Zink D, et al. Dynamics of DNA Replication Factories in Living Cells. J Cell Biol [Internet]. 2000 Apr 17;149(2):271–80. Available from: https://rupress.org/jcb/article/149/2/271/32118/Dynamics-of-DNA-Replication-Factories-in-Living

42. Cardoso MC, Joseph C, Rahn HP, Reusch R, Nadal-Ginard B, Leonhardt H. Mapping and use of a sequence that targets DNA ligase I to sites of DNA replication in vivo. J Cell Biol. 1997;139(3):579– 87.

43. Xie S, Swaffer M, Skotheim JM. Eukaryotic Cell Size Control and Its Relation to Biosynthesis and Senescence. Annu Rev Cell Dev Biol. 2022;38(1):1–29.

44. Cooper S. Control and maintenance of mammalian cell size. BMC Cell Biol. 2004;5:1–21.

45. Godin M, Delgado FF, Son S, Grover WH, Bryan AK, Tzur A, et al. Using buoyant mass to measure the growth of single cells. Nat Methods [Internet]. 2010 [cited 2013 Jan 10];7(5):387–90. Available from: http://www.nature.com/nmeth/journal/vaop/ncurrent/full/nmeth.1452.html

46. Scott M, Hwa T. Bacterial growth laws and their applications. Curr Opin Biotechnol [Internet]. 2011 May 16 [cited 2011 Jun 27];22(4):559–65. Available from: http://www.ncbi.nlm.nih.gov/pubmed/21592775

47. Son S, Tzur A, Weng Y, Jorgensen P, Kim J, Kirschner MW, et al. Direct observation of mammalian cell growth and size regulation. Nat Methods [Internet]. 2012 Sep [cited 2012 Nov 5];9(9):910–2. Available from: http://www.ncbi.nlm.nih.gov/pubmed/22863882

48. Mu L, Kang JH, Olcum S, Payer KR, Calistri NL, Kimmerling RJ, et al. Mass measurements during lymphocytic leukemia cell polyploidization decouple cell cycle-And cell size-dependent growth. Proc Natl Acad Sci U S A [Internet]. 2020 Jul 7;117(27):15659–65. Available from: http://biorxiv.org/cgi/content/short/2019.12.17.879080v1?rss=1&utm_source=researcher_app& utm_medium=referral&utm_campaign=RESR_MRKT_Researcher_inbound

49. Liu S, Tan C, Melo-gavin C, Mark KG, Ginzberg MB, Blutrich R, et al. Large cells activate global protein degradation to maintain cell size homeostasis. bioRxiv. 2021;1–31.

50. Miettinen TP, Kang JH, Yang LF, Manalis SR. Mammalian cell growth dynamics in mitosis. Elife. 2019;8:1–29.

51. Ghenim L, Allier C, Obeid P, Hervé L, Fortin J-Y, Balakirev M, et al. A new ultradian rhythm in mammalian cell dry mass observed by holography. Sci Rep [Internet]. 2021;11(1):1–12. Available from: 10.1038/s41598-020-79661-9

52. Kar P, Tiruvadi-Krishnan S, Männik J, Männik J, Amir A. Distinguishing different modes of growth using single-cell data. Elife [Internet]. 2021 Dec 2;10. Available from: https://elifesciences.org/articles/72565

53. Amir A. Cell size regulation in bacteria. Phys Rev Lett. 2014;112(20):1–5.

54. Thomas P. Analysis of cell size homeostasis at the single-cell and population level. Front Phys. 2018;6(June).

55. Vargas-Garcia CA, Björklund M, Singh A. Modeling homeostasis mechanisms that set the target cell size. Sci Rep. 2020 Dec 1;10(1).

56. Ho P-Y, Lin J, Amir A. Modeling cell size regulation: From single-cell level statistics to molecular mechanisms and population level effects. 2018;

57. Thomas P, Terradot G, Danos V, Weiße AY. Sources, propagation and consequences of stochasticity in cellular growth. Nat Commun [Internet]. 2018;9(1):1–11. Available from: 10.1038/s41467-018-06912-9

58. Cook M, Tyers M. Size control goes global. Curr Opin Biotechnol [Internet]. 2007 Aug [cited 2012 Nov 5];18(4):341–50. Available from: http://www.ncbi.nlm.nih.gov/pubmed/17768045

59. Echave P, Conlon IJ, Lloyd AC. Cell Size Regulation in Mammalian Cells. Cell Cycle [Internet]. 2007 Oct 28 [cited 2015 Feb 16];6(2):218–24. Available from: http://www.tandfonline.com/doi/abs/10.4161/cc.6.2.3744

60. Jorgensen P, Tyers M. How cells coordinate growth and division. Curr Biol [Internet]. 2004 Dec 14 [cited 2012 Mar 6];14(23):R1014-27. Available from: http://www.ncbi.nlm.nih.gov/pubmed/15589139

61. Umen JG. The elusive sizer. Curr Opin Cell Biol [Internet]. 2005 Aug [cited 2012 May 21];17(4):435–41. Available from: http://www.ncbi.nlm.nih.gov/pubmed/15978795

62. Liu S, Tan C, Tyers M, Zetterberg A, Kafri R. What programs the size of animal cells? Front Cell Dev Biol. 2022;10(November):1–19.

63. Taubenberger A V., Baum B, Matthews HK. The Mechanics of Mitotic Cell Rounding. Front Cell Dev Biol. 2020;8(August):1–16.

64. Voldner N, Frey Frøslie K, Godang K, Bollerslev J, Henriksen T. Determinants of birth weight in boys and girls. human_ontogenetics [Internet]. 2009 Mar 18;3(1):7–12. Available from: https://onlinelibrary.wiley.com/doi/10.1002/huon.200900001

65. Amir A. Is cell size a spandrel? Elife. 2017;6:1–8.

66. ElGamel M, Mugler A. Effects of molecular noise on cell size control. 2023;(1). Available from: http://arxiv.org/abs/2303.15232

67. Liu X, Oh S, Kirschner MW. The uniformity and stability of cellular mass density in mammalian cell culture. Front Cell Dev Biol [Internet]. 2022 [cited 2022 Oct 14]; Available from: https://internal-journal.frontiersin.org/articles/10.3389/fcell.2022.1017499/full

68. Cadart C, Venkova L, Piel M, Cosentino Lagomarsino M. Volume growth in animal cells is cell cycle dependent and shows additive fluctuations. Elife. 2022;11.

69. Cadart C, Venkova L, Recho P, Lagomarsino MC, Piel M. The physics of cell-size regulation across timescales. Nat Phys [Internet]. 2019;15(10):993–1004. Available from: 10.1038/s41567-019-0629-y

70. Björklund M. Cell size homeostasis: Metabolic control of growth and cell division. Biochim Biophys Acta - Mol Cell Res [Internet]. 2019 Mar;1866(3):409–17. Available from: 10.1016/j.bbamcr.2018.10.002

71. Cadart C, Heald R. Scaling of biosynthesis and metabolism with cell size. Mol Biol Cell. 2022;33(9):1–6.

72. Turner JJ, Ewald JC, Skotheim JM. cell size control in yeast. Curr Biol. 2012;29(6):997–1003.

73. Navarro FJ, Nurse P. A systematic screen reveals new elements acting at the G2/M cell cycle control. Genome Biol. 2012;13(5).

74. Keifenheim D, Sun XM, D’Souza E, Ohira MJ, Magner M, Mayhew MB, et al. Size-Dependent Expression of the Mitotic Activator Cdc25 Suggests a Mechanism of Size Control in Fission Yeast. Curr Biol [Internet]. 2017;27(10):1491–1497.e4. Available from: 10.1016/j.cub.2017.04.016

75. Donzelli M, Draetta GF. Regulating mammalian checkpoints through Cdc25 inactivation. EMBO Rep. 2003;4(7):671–7.

76. Zhao RY, Elder RT. Viral infections and cell cycle G2/M regulation. Cell Res. 2005;15(3):143–9.

77. Gemble S, Bernhard SV, Srivastava N, Wardenaar R, Nano M, Macé A-S, et al. Mechanisms of genetic instability in a single S-phase following whole genome doubling. bioRxiv [Internet]. 2021;2021.07.16.452672. Available from: https://www.biorxiv.org/content/10.1101/2021.07.16.452672v1*0Ahttps://www.biorxiv.org/content/10.1101/2021.07.16.452672v1.abstract

78. Mei L, Cook JG. Efficiency and equity in origin licensing to ensure complete DNA replication. Biochem Soc Trans [Internet]. 2021 Nov 1;49(5):2133–41. Available from: https://portlandpress.com/biochemsoctrans/article/49/5/2133/229829/Efficiency-and-equity-in-origin-licensing-to

79. Maya-Mendoza A, Moudry P, Merchut-Maya JM, Lee M, Strauss R, Bartek J. High speed of fork progression induces DNA replication stress and genomic instability. Nature [Internet]. 2018;559(7713):279–84. Available from: 10.1038/s41586-018-0261-5

80. Arora M, Moser J, Phadke H, Basha AA, Spencer SL. Endogenous Replication Stress in Mother Cells Leads to Quiescence of Daughter Cells. Cell Rep [Internet]. 2017;19(7):1351–64. Available from: 10.1016/j.celrep.2017.04.055

81. Chen Y, Futcher B. Scaling gene expression for cell size control and senescence in Saccharomyces cerevisiae. Current Genetics. Springer Science and Business Media Deutschland GmbH; 2020.

82. Lin J, Amir A. Homeostasis of protein and mRNA concentrations in growing cells. Nat Commun [Internet]. 2018 Dec 29;9(1):4496. Available from: https://www.biorxiv.org/content/early/2018/01/29/255950

83. Harris LK, Theriot JA. Relative rates of surface and volume synthesis set bacterial cell size. Cell. 2016;165(6):1479–92.

84. Rishal I, Kam N, Perry RBT, Shinder V, Fisher EMCC, Schiavo G, et al. A Motor-Driven Mechanism for Cell-Length Sensing. Cell Rep [Internet]. 2012;1(6):608–16. Available from: 10.1016/j.celrep.2012.05.013

85. Laplante M, Sabatini DM. mTOR signaling in growth control and disease. Cell [Internet]. 2012 Apr 13 [cited 2014 Jul 9];149(2):274–93. Available from: http://www.pubmedcentral.nih.gov/articlerender.fcgi?artid=3331679&tool=pmcentrez&rendertype=abstract

86. Kumari R, Jat P. Mechanisms of Cellular Senescence: Cell Cycle Arrest and Senescence Associated Secretory Phenotype. Front Cell Dev Biol. 2021 Mar 29;9:485.

87. Ginzberg MB. Size control and uniformity in animal cells. Harvard University; 2015.

88. Burnham, Kenneth P., Anderson DR. Model Selection and Multimodel Inference [Internet]. Burnham KP, Anderson DR, editors. New York, NY: Springer New York; 2004. Available from: http://link.springer.com/10.1007/b97636

89. Abdi H. Kendall Rank Correlation Coefficient. Concise Encycl Stat. 2008;278–81.

