## Supplementary Information for "Beyond G1/S regulation: How cell size homeostasis is tightly controlled throughout the cell cycle?"

SI Text

Figure S1 to S13

Table S1 to S10

Legend for Movie S1

**SI Text**

### 1. An abstract model of cell mass homeostasis

In this section, we outline the abstract model employed to simulate the results in Figure 1A-C. The purpose of developing this model is to investigate the effects of different G1 regulation strengths, which we characterize as the slope of the correlation between birth mass and G1 length. In this stochastic model, we make the following assumptions: all cells accumulate mass exponentially, the length of the nonG1 phase remains constant, and the G1 length regulation is the only means to control cell size.

The G1 duration is determined by birth mass

$$G1 = (a_{G1}m_b + b_{G1})(m_b \leq x0_{length\_G1}) + (a_{G1}x0_{length\_G1} + b_{G1})(m_b > x0_{length\_G1}),$$

where $m_b$ denotes birth mass, $a_{G1}$, $b_{G1}$, and $x0_{length\_G1}$ are parameters reflecting the strength of the G1 length regulation. By varying these parameters, we simulate the behaviors of cell populations with different G1 regulation strengths.

For simplicity, we introduced Gaussian noise to the division mass at the initiation of each simulation, the cell cycle length, and the exponent of exponential growth, with their means and coefficients of variation (CVs) summarized in the Table in this section. In each iteration, we simulated a population of 1000 cells for 20 generations.

At the end of each generation, cells divide their mass in half with Gaussian noise, $d_{CV}$, to represent asymmetrical partition. Only one of the two daughter cells is retained for the subsequent generation. We also implemented a cutoff for the entire cell cycle length, $T_{max}$. If the sum of the G1 and nonG1 lengths of a cell exceeds $T_{max}$, its cell cycle length is truncated at $T_{max}$.

The cell mass of cell $i$ in generation $k$ at the time step $j$, $m_j^{i,k}$, is determined by

$$m_j^{i,k} = m_{j-1}^{i,k} + m_{j-1}^{i,k}\alpha^{i,k}dt, \text{ where } j = 2, \ldots, [\frac{T^{i,k}}{dt}].$$

where $\alpha^{i,k}$ and $T^{i,k}$ are the growth exponent and the cell cycle length of cell $i$ in generation $k$, respectively.

The simulation results are presented in Fig. 1A-C.

**Table. The meanings and values of parameters used to simulate the results in Figure 1A-C.**

| Parameter | Meaning | Blue No G1 length regulation | Red Strong G1 length regulation | Yellow Weak G1 length regulation |
|---|---|---|---|---|
| $\mu_{md}$ | The mean cell mass at division within the population at the initiation of each simulation | 1000 | - | - |
| $CV_{md}$ | The CV of the cell mass at division within the population at the initiation of each simulation | 0.15 | - | - |
| $d_{CV}$ | The CV of the partition error | 0.05 | - | - |
| $T_{max}$ | The maximum cell cycle length | 48 | - | - |
| $\mu_{nonG1}$ | The mean nonG1 length | 12.5 | - | - |
| $CV_T$ | The CV of the cell cycle length | 0.18 | - | - |
| $dt$ | The time step of simulations | 0.5 | - | - |
| $\mu_\alpha$ | The mean growth exponent | 0.0277 | | |
| $CV_\alpha$ | The CV of the growth exponent | 0.15 | - | - |
| $a_{G1}$ | Parameter of the G1 regulation | 0 | -0.053 | -0.017 |
| $b_{G1}$ | Parameter of the G1 regulation | 12.5 | 39 | 21 |
| $x0_{length\_G1}$ | Parameter of the G1 regulation | Inf | 717 | 1176.5 |

### 2. Simulation of the impact of minimal cell cycle length on cell mass variation

In this section, we describe the stochastic model we used for generating the results in Fig. S6.

To investigate the impact of minimal cell cycle length on cell mass variation, we simulated cell populations across multiple generations, assuming exponential growth in individual cell masses. The birth mass of cells in the first generation, $mb_i^1$, follows a normal distribution with a mean of $\mu_{mb}$ and a standard deviation of $\mu_{mb} * CV_{mb}$. Here, $i = 1, \ldots, 10000$ is the ID of the cells.

$$mb_i^1 \sim N(\mu_{mb}, (\mu_{mb} * CV_{mb})^2) .$$

The cell cycle length of cell $i$ in generation $j$, $T_i^j$, is determined by the birth mass, $mb_i^j$, according to a bilinear function

$$T_i^j = ((a \cdot mb_i^j + b)(mb_i^j \leq m_{thresh}) + (a \cdot m_{thresh} + b)(mb_i^j > m_{thresh}))(1 + \delta_t),$$

where $m_{thresh}$ represents the 100$^{th}$, 90$^{th}$, 80$^{th}$, 70$^{th}$, 60$^{th}$, or 50$^{th}$ percentile of the $mb_i^1$ distribution. $\delta_t$ accounts for noise in the cell cycle length, which follows a normal distribution, $\delta_t \sim N(0, CV_t^2)$.

The cell mass at division, $md_i^j$, is determined by

$$md_i^j = mb_i^j e^{\alpha T_i^j},$$

where $\alpha$ represents the exponent of exponential growth.

The birth mass of the subsequent generation is determined by

$$mb_i^{j+1} = md_i^j (0.5 + \delta_p),$$

where $\delta_p$ represents the noise in the symmetric partition and follows a normal distribution, $\delta_p \sim N(0, p\_err^2)$.

The simulation was conducted for 100 generations.

**Table. Parameter values used for generating results in Fig. S6.**

| | |
|---|---|
| $\mu_{mb}$ | 484 |
| $CV_{mb}$ | 0.18 |
| $p\_err$ | 0.051 |
| $\alpha$ | 0.0248 |
| $CV_t$ | 0.15 |
| $a$ | -0.0213 |
| $b$ | 38.3 |

The parameter values of $\mu_{mb}$, $CV_{mb}$, $\alpha$, $CV_t$, $a$, $b$, and $p\_err$ were derived from the HeLa cell data.

### 3. The impact of different forms of growth modulation on cell mass variation

In this section, we examined the impact of growth rate modulation and growth rate noise on the coefficient of variation in cell mass over a single cell cycle. We assumed that all cells divided at the same cell cycle length and that all populations started with the same variation in birth mass. To address this problem, we utilized a combination of analytical solutions and numerical simulations.

#### 3.1 Sub-exponential growth rate modulation

When growth rate modulation is in the Sub-exponential (SE) form, the correlation between cell mass and growth rate can be expressed as

$$\frac{dm}{dt} = \alpha m + \beta. \qquad \text{(Eq. 2)}$$

Equation (2) has the following solution with initial condition $m(0) = m_0$:

$$m(t) = m_0 e^{\alpha t} + \beta(e^{\alpha t} - 1)/\alpha. \qquad \text{(Eq. 3)}$$

Here, $m(t)$, $m_0$, $\alpha$, $\beta$, and $t$ are normalized by the mean birth mass and the mean cell cycle length.

##### 3.1.1 Analytical solutions

We are interested in understanding how the coefficient of variation (CV) of $m(t)$ changes, particularly in identifying the parameters that lead to a decrease in CV over time.

When $\beta = 0$, cell mass accumulates exponentially.

We have

$$CV_{m(t)}^2 = \frac{\langle m(t)^2 \rangle - \langle m(t) \rangle^2}{\langle m(t) \rangle^2}.$$

The variability in $\alpha$ arises from two sources: the stochastic partitioning of cellular contents during cell division and intrinsic fluctuations in biochemical reactions (1). For the sake of mathematical simplicity, we assume that the mean and CV of $\alpha$ do not change with time and are independent of $m$. The assumption that CV of $\alpha$ is independent of cell mass is supported by the experimental data in Fig. S11A-B. Both $m_b$ and $\alpha$ follow normal distributions. Then, by using the property of moment-generating function, the CV of $m(t)$, $CV_{m(t)}^2$, can be calculated analytically as:

$$CV_{m(t)}^2 = \frac{\langle m_0^2 \rangle \langle e^{2\alpha t} \rangle - \langle m_0 \rangle^2 \langle e^{\alpha t} \rangle^2}{\langle m_0 \rangle^2 \langle e^{\alpha t} \rangle^2} = \frac{(\sigma_m^2 + \mu_m^2) e^{2\mu_\alpha t + 2\sigma_\alpha^2 t^2} - \mu_m^2 e^{2\mu_\alpha t + \sigma_\alpha^2 t^2}}{\mu_m^2 e^{2\mu_\alpha t + \sigma_\alpha^2 t^2}},$$

$$CV_{m(t)}^2 = \left(CV_{m_0}^2 + 1\right) e^{\sigma_\alpha^2 t^2} - 1\,,$$

which shows that <u>the CV of $m(t)$ increases super-exponentially with cell cycle progression if there were no control mechanisms.</u>

When $\beta \neq 0$, the CV of $m(t)$ becomes too complex to obtain a closed-form solution, but we can still apply a Taylor expansion to the solution to determine how CV changes at the beginning of the cell cycle:

$e^{\alpha t} = \sum_{n=0}^{\infty} \frac{(\alpha t)^n}{n!}$, and $(e^{\alpha t} - 1)/\alpha = t \sum_{n=0}^{\infty} \frac{(\alpha t)^n}{(n+1)!}$

Generically, we have:

$$\langle m(t)^2 \rangle = \langle m_0^2 \rangle \langle e^{2\alpha t} \rangle + 2\langle m_0 \rangle \langle \beta e^{\alpha t}(e^{\alpha t}-1)/\alpha \rangle + \langle m_0 \rangle^2 \langle \beta^2 (e^{\alpha t}-1)^2/\alpha^2 \rangle$$

$$= \langle m_0^2 \rangle \langle e^{2\alpha t} \rangle + 2t \sum_{n,m=0}^{\infty} \frac{\langle \alpha^{n+m}\beta \rangle}{n!(m+1)!} t^{n+m} + t^2 \sum_{n,m=0}^{\infty} \frac{\langle \alpha^{n+m}\beta^2 \rangle}{(n+1)!(m+1)!} t^{n+m},$$

$$\langle m(t) \rangle^2 = \langle m_0 \rangle^2 \langle e^{\alpha t} \rangle^2 + 2\langle m_0 \rangle \langle e^{\alpha t} \rangle \langle \beta(e^{\alpha t}-1)/\alpha + \langle m_0 \rangle^2 \langle \beta(e^{\alpha t}-1)/\alpha \rangle^2$$

$$= \langle e^{\alpha t} \rangle^2 + 2t \sum_{n,m=0}^{\infty} \frac{\langle \alpha^n \rangle \langle \alpha^m \beta \rangle}{n!(m+1)!} t^{n+m} + t^2 \sum_{n,m=0}^{\infty} \frac{\langle \alpha^n \beta \rangle \langle \alpha^m \beta \rangle}{(n+1)!(m+1)!} t^{n+m},$$

where $\langle m_0 \rangle = 1$ due to the normalization.

Therefore if we only keep lower-order terms, we have:

$$\langle m(t)^2 \rangle \approx (\sigma_m^2 + 1) + 2\left[(\sigma_m^2+1)\langle \alpha \rangle + \langle \beta \rangle\right] + \left[2(\sigma_m^2+1)\langle \alpha^2 \rangle + 3\langle \alpha\beta \rangle + \langle \beta^2 \rangle\right] t^2,$$

$$\langle m(t) \rangle^2 \approx 1 + 2\left[\langle \alpha \rangle + \langle \beta \rangle\right] t$$
$$+\left[\left(\langle \alpha^2 \rangle + \langle \alpha \rangle^2\right) + \left(\langle \alpha\beta \rangle + 2\langle \alpha \rangle \langle \beta \rangle\right) + \langle \beta \rangle^2\right] t^2.$$

So the derivatives are approximately:

$$\frac{d}{dt} \langle m(t)^2 \rangle \approx 2\left[(\sigma_m^2+1)\langle \alpha \rangle + \langle \beta \rangle\right] + 2\left[2(\sigma_m^2+1)\langle \alpha^2 \rangle + 3\langle \alpha\beta \rangle + \langle \beta^2 \rangle\right] t,$$

$$\frac{d}{dt} \langle m(t) \rangle^2 \approx 2\left[\langle \alpha \rangle + \langle \beta \rangle\right]$$
$$+2\left[\left(\langle \alpha^2 \rangle + \langle \alpha \rangle^2\right) + \left(\langle \alpha\beta \rangle + 2\langle \alpha \rangle \langle \beta \rangle\right) + \langle \beta \rangle^2\right] t.$$

Since we are mostly interested in whether the CV is increasing or decreasing, i.e., the sign of $d_t CV$, we only need to compute the numerator part of the derivative. Combining, we have:

$$\frac{dCV_{m(t)}^2}{dt} \propto \frac{d\langle m(t)^2 \rangle}{dt} \langle m(t) \rangle^2 - \langle m(t)^2 \rangle \frac{d\langle m(t) \rangle^2}{dt}$$

$$\approx 2\left[(\sigma_m^2+1)\langle \alpha \rangle + \langle \beta \rangle\right] - 2(\sigma_m^2+1)\left[\langle \alpha \rangle + \langle \beta \rangle\right]$$

$$+4\left[(\sigma_m^2+1)\langle \alpha \rangle + \langle \beta \rangle\right]\left[\langle \alpha \rangle + \langle \beta \rangle\right] t - 4\left[\langle \alpha \rangle + \langle \beta \rangle\right]\left[(\sigma_m^2+1)\langle \alpha \rangle + \langle \beta \rangle\right] t$$

$$+2\left[2(\sigma_m^2+1)\langle \alpha^2 \rangle + 3\langle \alpha\beta \rangle + \langle \beta^2 \rangle\right] t$$

$$-2(\sigma_m^2+1)\left[\left(\langle\alpha^2\rangle + \langle\alpha\rangle^2\right)+\left(\langle\alpha\beta\rangle + 2\langle\alpha\rangle\langle\beta\rangle\right)+\langle\beta\rangle^2\right]t$$

$$\approx -2\sigma_m^2\mu_\beta + 2\left[(\sigma_m^2+1)\sigma_\alpha^2 + \sigma_\beta^2 + (2-\sigma_m^2)\sigma_{\alpha\beta} - \sigma_m^2{\mu_\beta}^2 - 3\sigma_m^2\mu_\alpha\mu_\beta\right]t.$$

Given our assumption of $\mu_\beta \geq 0$, the 0th order is always non-positive. Thus, when $\beta \neq 0$, the cell mass CV always decreases at the beginning of the cell cycle. Additionally, the linear term suggests that if $\beta$ is not large enough, the cell mass CV could eventually increase after an initial brief decline.

The results above show that the cell size variation consistently decreases to a certain extent at the beginning of the cell cycle. However, they do not indicate whether the overall cell size variation can be maintained or reduced after an entire cell cycle ($t = 1$):

$$CV_{m(1)} \leq CV_{m(0)}.$$

Because of the nature of Taylor expansion, the results above cannot be readily extended to the scenario where $t = 1$. Therefore, rather than presenting an exact analytical solution, we will now derive a lower bound for $\mu_\beta$ that leads to a reduction in cell size variation.

First, we notice that $CV_{m(1)} \leq CV_{m(0)}$ can be rephrased as determining the condition under which the following inequality holds:

$$\frac{\langle m(1)^2\rangle - \langle m(1)\rangle^2}{\langle m(1)\rangle^2} \leq \frac{\langle m_0^2\rangle - \langle m_0\rangle^2}{\langle m_0\rangle^2}.$$

Since we also require that cells can maintain the same division size, we have the condition:

$$\langle m(1)\rangle = 2\langle m_0\rangle = 2.$$

The inequality above simply becomes:

$$\langle m(1)^2\rangle \leq 4(\sigma_m^2+1).$$

From above we find that

$$\langle m(1)^2\rangle = \langle m_0^2\rangle\langle e^{2\alpha}\rangle + 2\sum_{n,m=0}^{\infty}\frac{\langle\alpha^{n+m}\beta\rangle}{n!\,(m+1)!} + \sum_{n,m=0}^{\infty}\frac{\langle\alpha^{n+m}\beta^2\rangle}{(n+1)!\,(m+1)!}$$

$$< \langle m_0^2\rangle\langle e^{2\alpha}\rangle + 2\sum_{n,m=0}^{\infty}\frac{\langle\alpha^{n+m}\beta\rangle}{n!\,m!} + \sum_{n,m=0}^{\infty}\frac{\langle\alpha^{n+m}\beta^2\rangle}{n!\,m!}$$

$$= (\sigma_m^2+1)\langle e^{2\alpha}\rangle + 2\langle\beta e^{2\alpha}\rangle + \langle\beta^2 e^{2\alpha}\rangle,$$

$$\langle\beta e^{2\alpha}\rangle = \langle\beta e^{2\sigma_{\alpha\beta}\frac{\beta-\mu_\beta}{\sigma_\beta^2}}\rangle e^{2\mu_\alpha+2\sigma_\alpha^2(1-\rho^2)} = (\mu_\beta + 2\sigma_{\alpha\beta})e^{2\sigma_\alpha^2\rho^2+2\mu_\alpha+2\sigma_\alpha^2(1-\rho^2)}$$

$$= (\mu_\beta + 2\sigma_{\alpha\beta})e^{2(\mu_\alpha+\sigma_\alpha^2)},$$

$$\langle \beta^2 e^{2\alpha} \rangle = \langle \beta^2 e^{2\sigma_{\alpha\beta}\frac{\beta-\mu_\beta}{\sigma_\beta^2}} \rangle e^{2\mu_\alpha+2\sigma_\alpha^2(1-\rho^2)} = \left[\sigma_\beta^2 + \left(\mu_\beta + 2\sigma_{\alpha\beta}\right)^2\right] e^{2(\mu_\alpha+\sigma_\alpha^2)},$$

$$\langle m(1)^2 \rangle < \left[\sigma_m^2 + 1 + 2\left(\mu_\beta + 2\sigma_{\alpha\beta}\right) + \left(\mu_\beta + 2\sigma_{\alpha\beta}\right)^2 + \sigma_\beta^2\right] e^{2(\mu_\alpha+\sigma_\alpha^2)}.$$

Therefore, we obtained an upper bound on $\mu_\alpha$, $\mu_\beta$, $\sigma_\alpha^2$,$\sigma_\beta^2$, and $\sigma_{\alpha\beta}$ to achieve a reduction in cell size variation:

$$1 + \frac{2(\mu_\beta+2\sigma_{\alpha\beta})+(\mu_\beta+2\sigma_{\alpha\beta})^2+\sigma_\beta^2}{\sigma_m^2+1} \le 4e^{-2(\mu_\alpha+\sigma_\alpha^2)}.$$

Similarly, the constraint of $\langle m(1) \rangle = 2 \langle m_0 \rangle = 2$ leads to a lower bound:

$$2 \le e^{\mu_\alpha+\frac{1}{2}\sigma_\alpha^2} + \langle \beta e^{\sigma_{\alpha\beta}\frac{\beta-\mu_\beta}{\sigma_\beta^2}} \rangle e^{\mu_\alpha+\frac{1}{2}\sigma_\alpha^2(1-\rho^2)} = \left(1 + \mu_\beta + \sigma_{\alpha\beta}\right) e^{\mu_\alpha+\frac{1}{2}\sigma_\alpha^2}.$$

Together, the parameter regions required to achieve cell size CV reduction and double the average cell size are as follows:

$$1 + \frac{2(\mu_\beta+2\sigma_{\alpha\beta})+(\mu_\beta+2\sigma_{\alpha\beta})^2+\sigma_\beta^2}{\sigma_m^2+1} \le 4e^{-2(\mu_\alpha+\sigma_\alpha^2)},$$

$$\left(1 + \mu_\beta + \sigma_{\alpha\beta}\right) \ge 2e^{-\mu_\alpha-\frac{1}{2}\sigma_\alpha^2}.$$

Combining these two inequalities, we have:

$$\frac{(\sigma_m^2+1)(1+\mu_\beta+\sigma_{\alpha\beta})^2}{(\sigma_m^2+\sigma_\beta^2)+(1+\mu_\beta+2\sigma_{\alpha\beta})^2} \ge e^{\sigma_\alpha^2}.$$

For simplicity, we assume $\sigma_{\alpha\beta} = 0$, then:

$$\left[\sigma_m^2 - \left(e^{\sigma_\alpha^2} - 1\right)\right]\left(1 + \mu_\beta\right)^2 \ge e^{\sigma_\alpha^2}\left(\sigma_m^2 + \sigma_\beta^2\right).$$

This leads to the conditions:

$$\mu_\beta \ge e^{\frac{1}{2}\sigma_\alpha^2}\sqrt{1 + \sigma_\beta^2/\sigma_m^2} - 1 \text{ and } \sigma_\alpha^2 \le log(1 + \sigma_m^2).$$

In conclusion, in order to maintain or reduce cell size variation throughout the cell cycle, two conditions must be met: the mean of $\beta$ should not be too small, and the variation of $\alpha$ should not be too large.

#### 3.1.2 Numerical simulations.

We also validated the conclusions drawn from the analytical solutions through numerical simulations.

For universality, we normalized the correlation between growth rate and cell mass using the means of birth mass and cell cycle length as follows:

$$\frac{dm'}{dt'} = \alpha' m' + \beta',$$

where $m' = \frac{m}{<m_0>}$, $t' = \frac{t}{<T>}$, $\alpha' = \alpha < T >$, $\beta' = \beta \frac{<T>}{<m_0>}$.

For simplicity, we assumed that $\frac{\sigma_\beta}{<\beta>} = \frac{\sigma_\alpha}{<\alpha>}$, $\sigma_{\alpha\beta} = 0$, and that cells could maintain the same division size. Thus from Eq. 3, we have:

$$2 =< e^{\alpha'} + \frac{\beta'}{\alpha'}(e^{\alpha'} - 1) >.$$

From this, we derive:

$$< \beta' >= \frac{2-e^{<\alpha'>+\frac{1}{2}Var(\alpha')}}{e^{<\alpha'>+\frac{1}{2}Var(\alpha')}-1} < \alpha' >. \quad \text{(Eq. 4)}$$

As a result, the CV of $m(t)$ only depends on the mean and variation of $\alpha'$. In our simulations, we varied the mean of $\alpha'$ within the range of 0 to ln(2). Previous studies in the literature have reported that the CV of cell mass growth rate typically falls between 10% and 25%(2–6). Therefore we considered a range of CV for $\alpha'$ from 0 to 40%. Since all the cell lines we investigated demonstrated similar variability in birth mass (Fig. 1E), we adopted a constant birth mass CV, $\sigma_{m_0} = 0.22$. Thus the birth mass, $m_i'(0)$, follows a normal distribution:

$$m_i'(0) \sim N(1, (\sigma_{mb})^2).$$

For cell $i$, we have:

$$m_i(t' + \Delta t') = m_i(t') + (\alpha_i' m_i'(t') + \beta_i{}')\Delta t',$$

with both $\alpha_i'$ and $\beta_i'$ follow normal distributions:

$$\alpha_i' \sim N(\mu_{\alpha'}, (\mu_{\alpha'} CV_{\alpha'})^2),$$

$$\beta_i' \sim N\left(\mu_{\beta'}, \left(\mu_{\beta'} CV_{\alpha'}\right)^2\right),$$

where $\mu_{\beta'}$ is determined by Eq. 4.

The simulation results were plotted in Fig. S10A-B and Fig. 4G, confirming the findings derived from the analytical solutions:

1. The cell mass CV decreases at the beginning of the cell cycle, with the rate of reduction inversely related to the mean of $\alpha'$ and unaffected by the CV of $\alpha'$ (Fig. S10A).
2. The rate of mass CV reduction diminishes as cell cycle progresses and can potentially turn positive towards the latter stages of the cell cycle (Fig. S10B).
3. The overall change in cell mass CV over the entire cell cycle is contingent on both the mean and CV of $\alpha'$.
4. Smaller mean and CV for $\alpha'$ correspond to more significant reductions in cell mass CV (Fig. 4G).

We further examined scenarios where growth rate variation was introduced solely to $\alpha'$ or $\beta'$. Importantly, these variations did not significantly impact our major conclusions (Fig. S10C-H).

#### 3.2 Bilinear growth rate modulation

When growth rate modulation follows a bilinear (BI) form, the correlation between cell mass and growth rate can be expressed as

$$\frac{dm}{dt} = \alpha m(m < m_\tau) + (\gamma m + \alpha m_\tau - \gamma m_\tau)(m \geq m_\tau).$$

We normalized this expression by the means of birth mass and cell cycle length:

$$\frac{dm'}{dt'} = \alpha' m'(m' < m'_\tau) + (\gamma' m' + \alpha' m'_\tau - \gamma' m'_\tau)(m' \geq m'_\tau),$$

where $m' = \frac{m}{<m_b>}, t' = \frac{t}{<T>}, \alpha' = \alpha < T >, < \alpha' >= ln2,\ \gamma' = \gamma < T >, and\ m'_\tau = \frac{m_\tau}{<m_b>}$.

Due to the complexity of the expression above, we turned to numerical simulations to explore the change in cell mass CV throughout the cell cycle. For each cell $i$, we have:

$$m_i(t' + \Delta t') = m_i(t') + [\alpha_i{}' m_i{}'(m_i{}' < m'_{\tau i}) + (\gamma_i{}' m_i{}' + \alpha_i{}' m'_{\tau i} - \gamma_i{}' m'_{\tau i})(m_i{}' \geq m'_{\tau i})]t'.$$

For simplicity, we made the assumption that $\alpha'$, $\gamma'$, and $m_\tau{}'$ are independent Gaussian variables. We investigated the impact of the CV in each of these variables and compared the results to the scenario where all three variables share an equal CV (Fig. S12 A-D). We found that the primary factor driving an increase in the cell mass CV is the CV associated with $\alpha'$, which represents the exponential portion of the mass vs. growth correlation. Conversely, the impact of the CV in $\gamma'$ and $m_\tau{}'$ is relatively minor. Moreover, we examined the impact of the means of $\gamma'$ and $m_\tau{}'$ on the change in cell mass CV. We found that a smaller mean value of $\gamma'$, indicating a more pronounced growth rate modulation, and a smaller mean value of $m_\tau{}'$, signifying more cells being affected by the growth rate modulation, resulting in a more significant reduction in the cell mass CV (Fig. 4H-I, Fig. S12E-H).

4. **Simulations of the contribution of each control mechanism on cell mass variation in untreated HeLa and RPE-1 cells, as well as RPE-1 cells treated with palbociclib or rapamycin.**

We adopted parameters derived from experimental data to simulate the contribution of each control mechanism to cell mass variation. The meanings and values of these parameters are summarized in the Table in this section.

The CV of the cell cycle length, $CV_T$, was estimated from the averaged CV within each bin of the cell mass vs. cell cycle length correlation. For simplification, we only considered the extrinsic noise of growth rate fluctuation (Fig. S11C). We utilized the averaged slope and intercept of the cell mass vs. growth rate correlations for both the G1 and nonG1 phases to simulate size-dependent growth rate regulation across the entire cell cycle.

At the initiation of each simulation run, we generated the division mass distribution for the cell population, which followed a normal distribution, $md_{i,i=1,\dots,evt} \sim N(\mu_{md}, (\mu_{md} * CV_{md})^2)$. Subsequently, we conducted simulations to track the change in cell mass variation over the course of the cell cycle, culminating in the division of the subsequent generation. These simulations encompassed various scenarios designed to explore different conditions and factors influencing cell mass variation.

In the simulations, we implemented a cutoff for the entire cell cycle length, $T_{max}$. If the cell cycle length of a cell exceeds $T_{max}$, its cell cycle length is truncated at $T_{max}$.

<u>Scenario I. Without noise or control mechanisms</u>

The birth mass of a cell $i$ is determined by half of the mother cell division mass,

$$m_i^{j=1} = md_i/2.$$

The growth rate is represented as

$$gr = ln2/T,$$

where $T$ is the average cell cycle length.

Thus cell mass at any given point of the cell cycle can be calculated as

$$m_i^{j+1} = m_i^j + m_i^j gr \cdot dt,$$

where $j = 1, \dots, \frac{T}{dt} - 1$ is the simulation step.

<u>Scenario II. With partition error, without control mechanisms</u>

We used $\delta_p$ to represent the noise in cell partition, which follows a normal distribution, $\delta_p \sim N(0, {DA_{std}}^2)$.

As a result,

$$m_i^{j=1} = md_i(0.5 + \delta_p).$$

Similarly to Scenario I,

$$gr = ln2/T,$$

$$m_i^{j+1} = m_i^j + m_i^j gr \cdot dt,$$

$$j = 1, \ldots, \frac{T}{dt} - 1.$$

<u>Scenario III. With cell cycle variation, without control mechanisms</u>

$$m_i^{j=1} = md_i/2,$$

$$gr = ln2/T,$$

We add Gaussian noise to the G1 and nonG1 lengths, with the noise CV equal to $CV_T$,

$$G1_i = G1(1 + \delta_{G1}), \delta_{G1} \sim N(0, CV_T^2),$$

$$nonG1_i = nonG1(1 + \delta_{nonG1}), \delta_{nonG1} \sim N(0, CV_T^2).$$

The whole cell cycle length of cell $i$ is the sum of the G1 and nonG1 lengths:

$$T_i = G1_i + nonG1_i.$$

$$m_i^{j+1} = m_i^j + m_i^j gr \cdot dt,$$

$$j = 1, \ldots, \left\lceil \frac{T_i}{dt} \right\rceil - 1.$$

<u>Scenario IV. With growth rate variation, without control mechanisms</u>

$$m_i^1 = md_i/2,$$

We introduce Gaussian noise to growth rate, with the noise CV equal to $CV_{gr}$. Then the growth rate of cell $i$ is determined by

$$gr_i = ln2/T(1 + \delta_{gr}), \delta_{gr} \sim N(0, CV_{gr}^2).$$

$$m_i^{j+1} = m_i^j + m_i^j gr_i \cdot dt,$$

$$j = 1, \ldots, \frac{T}{dt} - 1.$$

<u>Scenario V. With all noise, without control mechanisms</u>

As described in Scenario II-IV, we incorporate Gaussian noise in cell partition, cell cycle length, and cell growth rate. For cell $i$, we have

$$m_i^{j=1} = md_i(0.5 + \delta_p), \delta_p \sim N(0, {DA_{std}}^2),$$

$$gr_i = ln2/T(1 + \delta_{gr}), \delta_{gr} \sim N(0, CV_{gr}^2),$$

$$G1_i = G1(1 + \delta_{G1}), \delta_{G1} \sim N(0, CV_T^2),$$

$$nonG1_i = nonG1(1 + \delta_{nonG1}), \delta_{nonG1} \sim N(0, CV_T^2),$$

$$T_i = G1_i + nonG1_i.$$

Thus cell mass at any given point of the cell cycle is determined by

$$m_i^{j+1} = m_i^j + m_i^j gr_i \cdot dt,$$

$$j = 1, \dots, \left\lceil \frac{T_i}{dt} \right\rceil - 1.$$

<u>Scenario VI. With all noise and G1 length control</u>

Similarly to Scenario V,

$$m_i^{j=1} = md_i\left(0.5 + \delta_p\right), \delta_p \sim N(0, {DA_{std}}^2),$$

$$gr_i = ln2/T(1 + \delta_{gr}), \delta_{gr} \sim N(0, CV_{gr}^2),$$

The G1 length is determined by a bilinear function:

$$G1_i = [\left(m_i^{j=1} \le m_{G1}\right) + (a_{G1}m_{G1} + b_{G1})\left(m_i^{j=1} > m_{G1}\right)](1 + \delta_{G1}),$$

where $a_{G1}$, $b_{G1}$, and $m_{G1}$ are parameters of the bilinear function, and $\delta_{G1}$ is the noise term, $\delta_{G1} \sim N(0, CV_T^2)$.

The nonG1 length follows a normal distribution with a mean of $nonG1$ and a CV of $CV_T$,

$$nonG1_i = nonG1(1 + \delta_{nonG1}), \delta_{nonG1} \sim N(0, CV_T^2),$$

The cell cycle length is the sum of the G1 and nonG1 lengths,

$$T_i = G1_i + nonG1_i.$$

$$m_i^{j+1} = m_i^j + m_i^j gr_i \cdot dt,$$

$$j = 1, \dots, \left\lceil \frac{T_i}{dt} \right\rceil - 1.$$

<u>Scenario VII. with all noise and nonG1 length control</u>

Similarly to Scenario V,

$$m_i^{j=1} = md_i\left(0.5 + \delta_p\right), \delta_p \sim N(0, {DA_{std}}^2),$$

$$gr_i = ln2/T(1 + \delta_{gr}), \delta_{gr} \sim N(0, {gr_{CV}}^2).$$

The G1 length follows a normal distribution with a mean of $G1$ and a CV of $CV_T$,

$$G1_i = G1(1 + \delta_{G1}), \delta_{G1} \sim N(0, {CV_T}^2).$$

Thus the cell mass at any given point of the G1 phase is determined by

$$m_i^{j_1+1} = m_i^{j_1} + m_i^{j_1} gr_i \cdot dt,$$

$$j_1 = 1, \dots, \left\lceil \frac{G1_i}{dt} \right\rceil - 1.$$

The cell mass at the G1/S transition is

$m_i^{G1\_S} = m_i^{\left\lceil \frac{G1_i}{dt} \right\rceil}$.

The nonG1 length is determined by a bilinear function,

$$nonG1_i = [(m_i^{G1\_S} \le m_{nonG1}) + (a_{nonG1}m_{nonG1} + b_{nonG1})(m_i^{G1\_S} > m_{nonG1})](1 + \delta_{nonG1}),$$

where $a_{nonG1}$, $b_{nonG1}$, and $m_{nonG1}$ are parameters of the bilinear function, and $\delta_{nonG1}$ is the noise term, $\delta_{nonG1} \sim N(0, CV_T^2)$.

The cell mass during the nonG1 phase is determined by

$$m_i^{\left\lceil \frac{G1_i}{dt} \right\rceil + j_2 + 1} = m_i^{\left\lceil \frac{G1_i}{dt} \right\rceil + j_2} + m_i^{\left\lceil \frac{G1_i}{dt} \right\rceil + j_2} gr_i \cdot dt,$$

$$j_2 = 1, \ldots, \left\lceil \frac{nonG1_i}{dt} \right\rceil - 1.$$

<u>Scenario VIII. With all noise and growth rate control</u>

Similarly to Scenario V, we incorporate Gaussian noise in cell partition and cell cycle length:

$$m_i^{j=1} = md_i(0.5 + \delta_p), \delta_p \sim N(0, {DA_{std}}^2),$$

$$G1_i = G1(1 + \delta_{G1}), \delta_{G1} \sim N(0, {CV_T}^2),$$

$$nonG1_i = nonG1(1 + \delta_{nonG1}), \delta_{nonG1} \sim N(0, {CV_T}^2),$$

$$T_i = G1_i + nonG1_i.$$

The cell growth rate is determined by a subexponential correlation between cell mass and growth rate, and we introduce Gaussian noise to both the slope and the intercept terms of the correlation:

$$m_i^{j+1} = m_i^j + (m_i^j \alpha_i + \beta_i) \cdot dt,$$

with $\alpha_i = \alpha(1 + \delta_{gr}), \delta_{gr} \sim N(0, CV_{gr}^2)$,

and $\beta_i = \beta(1 + \delta_{gr}), \delta_{gr} \sim N(0, CV_{gr}^2)$.

$$j = 1, \ldots, \left\lceil \frac{T_i}{dt} \right\rceil - 1.$$

<u>Scenario IX. With all noise and all control mechanisms</u>

As described in previous scenarios, we introduce Gaussian noise to cell partition:

$$m_i^{j=1} = md_i(0.5 + \delta_p), \delta_p \sim N(0, {DA_{std}}^2).$$

The G1 length is determined by a bilinear function,

$$G1_i = [(m_i^1 \le m_{G1}) + (a_{G1}m_{G1} + b_{G1})(m_i^1 > m_{G1})](1 + \delta_{G1}),$$

where $a_{G1}$, $b_{G1}$, and $m_{G1}$ are parameters of the bilinear function, and $\delta_{G1}$ is the noise term, $\delta_{G1} \sim N(0, CV_T^2)$.

Similarly, the nonG1 length is determined by another bilinear function,

$$nonG1_i = [(m_i^{G1\_S} \leq m_{nonG1}) + (a_{nonG1}m_{nonG1} + b_{nonG1})(m_i^{G1\_S} > m_{nonG1})](1 + \delta_{nonG1}),$$
$$\delta_{nonG1} \sim N(0, {CV_T}^2),$$

where $a_{nonG1}$, $b_{nonG1}$, and $m_{nonG1}$ are parameters of the bilinear function; $m_i^{G1\_S}$ is the cell mass at the G1/S transition, $m_i^{G1\_S} = m_i^{\left\lceil \frac{G1_i}{dt} \right\rceil}$; and $\delta_{nonG1}$ is the noise term, $\delta_{nonG1} \sim N(0, CV_T^2)$.

The cell growth rate is determined by a subexponential correlation between cell mass and growth rate, and we introduce Gaussian noise to both the slope and the intercept terms of the correlation.

Thus, the cell mass during the G1 phase is determined by

$$m_i^{j_1+1} = m_i^{j_1} + (m_i^{j_1}\alpha_i + \beta_i) \cdot dt,$$

$$j_1 = 1, \dots, \left\lceil \frac{G1_i}{dt} \right\rceil - 1;$$

whereas the cell mass during the nonG1 phase is determined by

$$m_i^{\left\lceil \frac{G1_i}{dt} \right\rceil + j2 + 1} = m_i^{\left\lceil \frac{G1_i}{dt} \right\rceil + j2} + (m_i^{\left\lceil \frac{G1_i}{dt} \right\rceil + j2}\alpha_i + \beta_i) \cdot dt,$$

$$j_2 = 1, \dots, \left\lceil \frac{nonG1_i}{dt} \right\rceil - 1,$$

where $\alpha_i$ and $\beta_i$ are parameters of the cell mass vs. growth rate correlation for cell $i$, drawn from two normal distributions, respectively. One distribution has a mean of $\alpha$ and a CV of $CV_{gr}$; the other distribution has a mean of $\beta$ and the same CV of $CV_{gr}$:

$$\alpha_i = \alpha(1 + \delta_{gr}), \delta_{gr} \sim N(0, CV_{gr}^2),$$

$$\beta_i = \beta(1 + \delta_{gr}), \delta_{gr} \sim N(0, CV_{gr}^2).$$

The simulated division mass CVs for each scenario under different treatment conditions were summarized in Table S8.

Table. Parameters used in the simulations in this section.

| | meaning | HeLa | RPE | RPE Palb | RPE Rapa |
|---|---|---|---|---|---|
| $evt$ | Number of cells | 1000 | 1000 | 1000 | 1000 |
| $T$ | Average cell cycle length | 28.0 | 20.0 | 24.2 | 33.5 |
| $G1$ | Average G1 length | 11.1 | 7.8 | 12.3 | 17.4 |
| $nonG1$ | Average nonG1 length | 17.1 | 12.3 | 11.9 | 16.1 |
| $dt$ | Simulation step size | 0.5 | 0.5 | 0.5 | 0.5 |
| $CV_T$ | CV of cell cycle phase length | 0.25 | 0.25 | 0.25 | 0.25 |
| $T_{max}$ | Maximum cell cycle length allowed in the simulation | 48 | 48 | 48 | 72 |
| $\mu_{md}$ | Average division mass | 897 | 785 | 1139 | 556 |
| $CV_{md}$ | Division mass CV at the initiation of the simulation | 0.18 | 0.23 | 0.21 | 0.20 |
| $D.A._{std}$ | Standard deviation of Division Asymmetry | 0.053 | 0.051 | 0.055 | 0.050 |
| $CV_{gr}$ | CV of growth rate | 0.23 | 0.33 | 0.43 | 0.30 |
| $a_{G1}$ | Slope of the birth mass vs. G1 length correlation | -0.0165 | -0.0132 | -0.0067 | -0.0889 |
| $b_{G1}$ | Intercept of the birth mass vs. G1 length correlation | 19.2 | 12.2 | 16.0 | 40.9 |
| $m_{G1}$ | Transition mass of the birth mass vs. G1 length correlation | 595 | 483 | Inf | 334 |
| $a_{nonG1}$ | Slope of the G1/S mass vs. nonG1 length correlation | -0.0237 | -0.0152 | -0.0294 | -0.0141 |
| $b_{nonG1}$ | Intercept of the G1/S mass vs. nonG1 length correlation | 30.9 | 18.6 | 31.95 | 21.2 |
| $m_{nonG1}$ | Transition of the G1/S mass vs. nonG1 length correlation | 699 | 459 | 701 | 387 |
| $\alpha$ | Slope of the cell mass vs. growth rate correlation | 0.0177 | 0.0141 | -0.00007 | 0.0158 |
| $\beta$ | Intercept of the cell mass vs. growth rate correlation | 6.87 | 9.74 | 22.20 | 1.47 |

**References**

1. Thomas P, Terradot G, Danos V, Weiße AY. Sources, propagation and consequences of stochasticity in cellular growth. Nat Commun [Internet]. 2018;9(1):1–11. Available from: http://dx.doi.org/10.1038/s41467-018-06912-9

2. Son S, Tzur A, Weng Y, Jorgensen P, Kim J, Kirschner MW, et al. Direct observation of mammalian cell growth and size regulation. Nat Methods [Internet]. 2012 Sep [cited 2012 Nov 5];9(9):910–2. Available from: http://www.ncbi.nlm.nih.gov/pubmed/22863882

3. Mu L, Kang JH, Olcum S, Payer KR, Calistri NL, Kimmerling RJ, et al. Mass measurements during lymphocytic leukemia cell polyploidization decouple cell cycle- And cell size-dependent growth. Proc Natl Acad Sci U S A [Internet]. 2020 Jul 7;117(27):15659–65. Available from: http://biorxiv.org/cgi/content/short/2019.12.17.879080v1?rss=1&utm_source=researcher_app&utm_medium=referral&utm_campaign=RESR_MRKT_Researcher_inbound

4. Godin M, Delgado FF, Son S, Grover WH, Bryan AK, Tzur A, et al. Using buoyant mass to measure the growth of single cells. Nat Methods [Internet]. 2010 [cited 2013 Jan 10];7(5):387–90. Available from: http://www.nature.com/nmeth/journal/vaop/ncurrent/full/nmeth.1452.html

5. Miettinen TP, Ly KS, Lam A, Manalis SR. Single-cell monitoring of dry mass and dry mass density reveals exocytosis of cellular dry contents in mitosis. Elife. 2022;11:1–20.

6. Mir M, Wang Z, Shen Z, Bednarz M, Bashir R, Golding I, et al. Optical measurement of cycle-dependent cell growth. Proc Natl Acad Sci U S A [Internet]. 2011 Aug 9 [cited 2013 May 22];108(32):13124–9. Available from: http://www.pubmedcentral.nih.gov/articlerender.fcgi?artid=3156192&tool=pmcentrez&rendertype=abstract

7. Liu X, Oh S, Peshkin L, Kirschner MW. Computationally enhanced quantitative phase microscopy reveals autonomous oscillations in mammalian cell growth. Proc Natl Acad Sci [Internet]. 2020 Nov 3;117(44):27388–99. Available from: http://www.pnas.org/lookup/doi/10.1073/pnas.2002152117

8. Kafri R, Levy J, Ginzberg MB, Oh S, Lahav G, Kirschner MW. Dynamics extracted from fixed cells reveal feedback linking cell growth to cell cycle. Nature [Internet]. 2013 Feb 27 [cited 2013 Feb 27];494(7438):480–3. Available from: http://www.nature.com/doifinder/10.1038/nature11897

9. Kar P, Tiruvadi-Krishnan S, Männik J, Männik J, Amir A. Distinguishing different modes of growth using single-cell data. Elife [Internet]. 2021 Dec 2;10. Available from: https://elifesciences.org/articles/72565

**Supplementary figures**

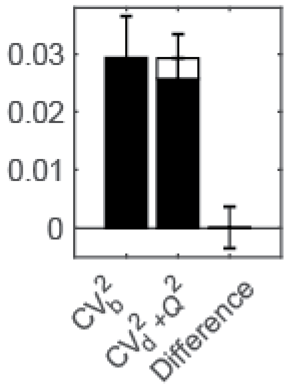

**Figure S1. The left- and right-hand sides of Eq. 1 and their difference quantified in HeLa cells.** In the term, $\mathrm{CV}_d^2 + Q^2$, $\mathrm{CV}_d^2$ is indicated in black, $Q^2$ is indicated in white; error bars are the standard deviation of 8 experiments.

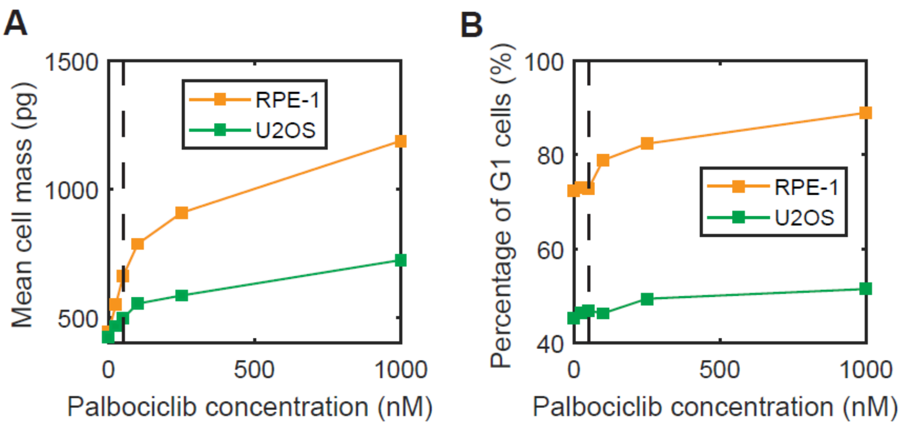

**Figure S2. RPE-1 and U2OS sensitivity to palbociclib.** The mean cell mass of the population (A) and the percentage of G1 cells quantified by low Geminin expression (B) after being treated in palbociclib at the indicated concentrations for 2 days. Dashed black lines show the concentration (50 nM) chosen for the analyses in this study.

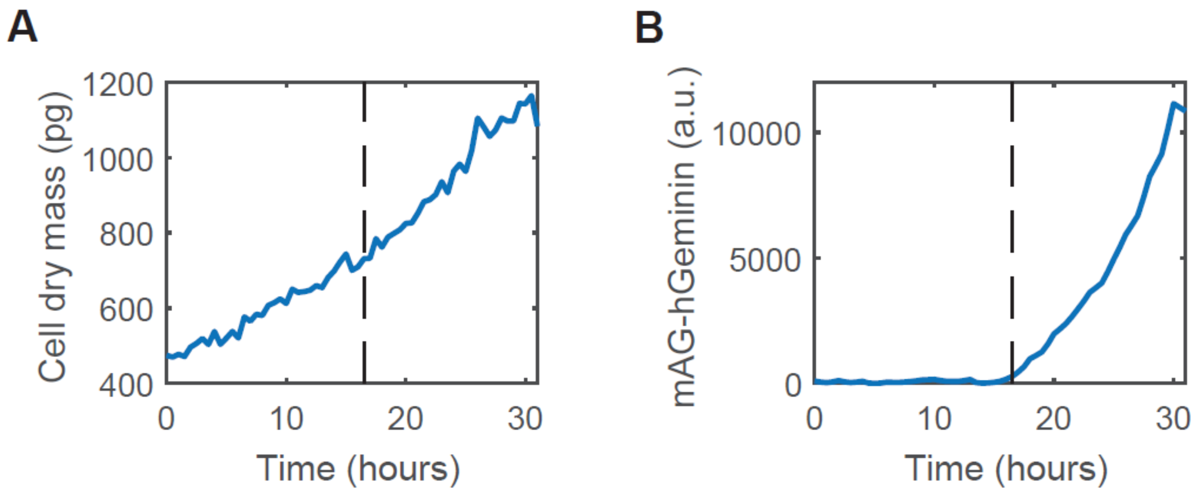

**Figure S3. Representative cell mass (A) and mAG-hGeminin (B) trajectories of the same HeLa cell.** Dashed lines denote the timing of the G1/S transition identified by the initiation of geminin accumulation.

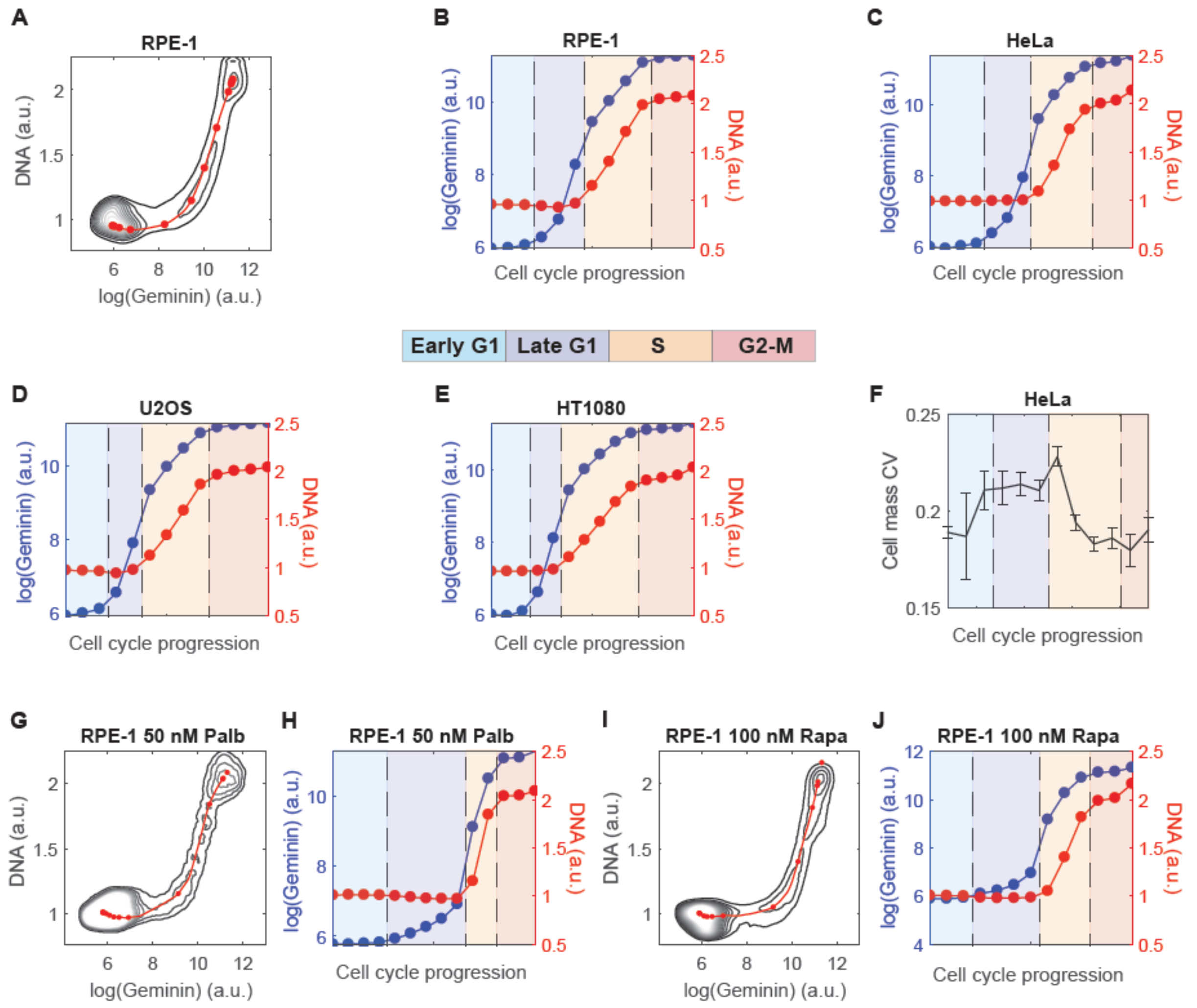

**Figure S4. Segregation of cells into stages along the cell cycle mean path.** (A) The 2D plane of the logarithmic scale of mAG-hGeminin intensity, log(Geminin), and the intensity of Hoechst fluorescence, DNA, in asynchronous RPE-1 cells. Black contours indicate cell number density; the solid red line is the cell cycle mean path; filled red circles show the centroids of the chosen stages along the mean path; the stages are evenly separated in the time axis computed by the Ergodic Rate Analysis (ERA) method(8). (B-E) The average of log(Geminin) (blue) and DNA content (red) change with cell cycle progression in different cell lines. X-axes are calculated by the ERA method(8). The cell cycle is segregated into four phases indicated by color-shaded areas: the early G1 phase from birth to the onset of geminin accumulation, the late G1 phase from the initiation of geminin accumulation to the onset of DNA replication, the S phase covering DNA replication, and the G2-M phase where geminin and DNA accumulation plateau. (F) Error in computed cell mass CV caused by inaccurate cell cycle stage identification. The cell dry mass and cell cycle markers data were from Fig. 2D. We added 10% random Gaussian noise to each cell’s position in the log(Geminin)-DNA plane. The cells were reassigned to cell cycle stages according to their new positions, and the cell mass CV of each stage was computed. The solid black line and error bars indicate the mean and standard deviation of computed cell mass CVs of 100 simulations; the first and last stages are truncated due to having much higher cell numbers and variations than other stages. (G, I) The 2D planes of log(Geminin) and DNA content in RPE-1 cells in 50 nM palbociclib (G) or 100 nM rapamycin (I). The red line and filled circles are the cell cycle mean path and centroids of stages calculated from the treated cells. (H, J) The average of log(Geminin) (blue) and DNA content (red) change with cell cycle progression in RPE-1 cells in 50 nM palbociclib (H) or 100 nM rapamycin (J).

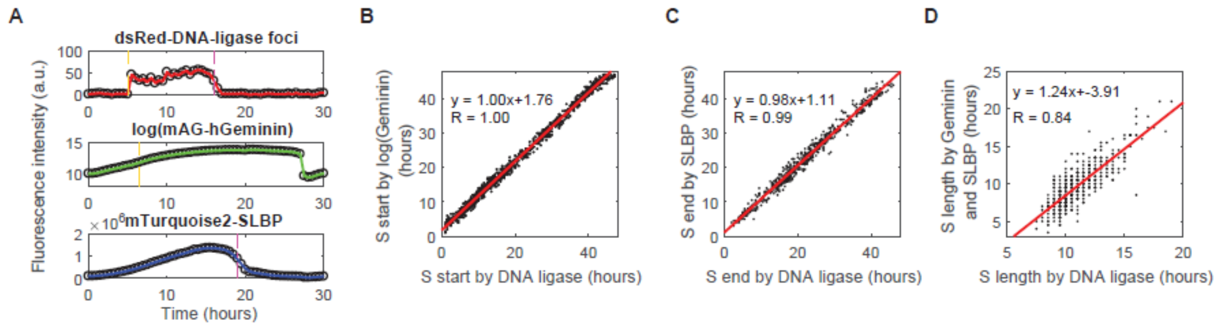

**Figure S5. The geminin and SLBP markers faithfully report the timing and duration of S phase.** (A) The trajectories of dsRed-DNA-ligaseI foci, mAG-hGeminin, and mTurquoise2-SLBP in a representative HeLa cell. Open circles are the raw data; solid colored lines are the spline interpolations; dashed yellow and pink lines mark the S phase start and end, respectively. (B-D) Correlations between the S phase start (B), end (C), and duration (D) identified by the dsRed-DNA-ligase foci or mAG-hGeminin and mTurquoise2-SLBP combined. Each black dot is one observation; Solid red lines are the best linear fit.

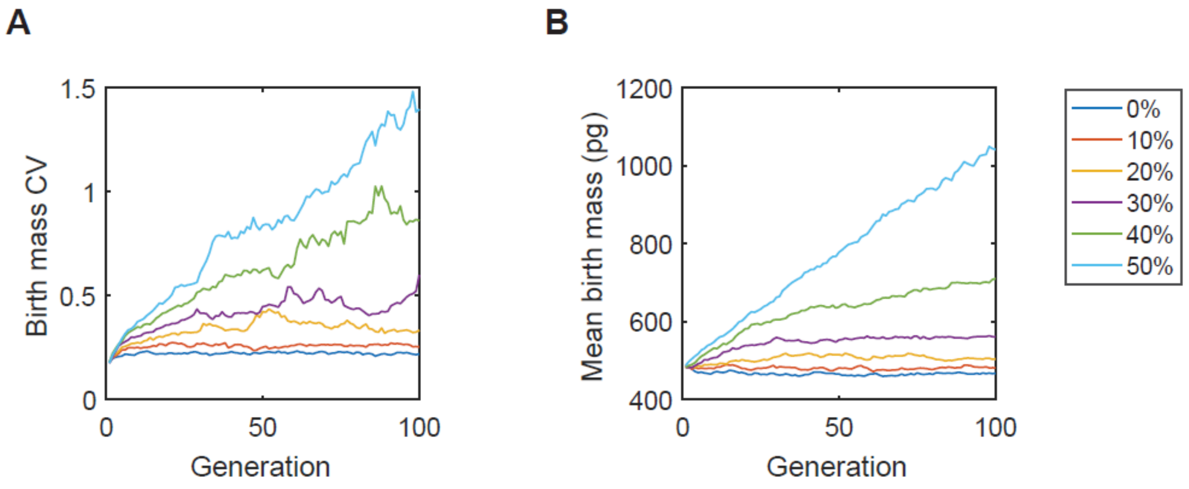

**Figure S6. The impact of minimal cell cycle length on cell mass homeostasis, indicated by the birth mass CV (A) and mean birth mass (B) changing with simulated generations.** Different colors show the percentage of cells affected by the minimal cell cycle length in the population of the first generation of simulations.

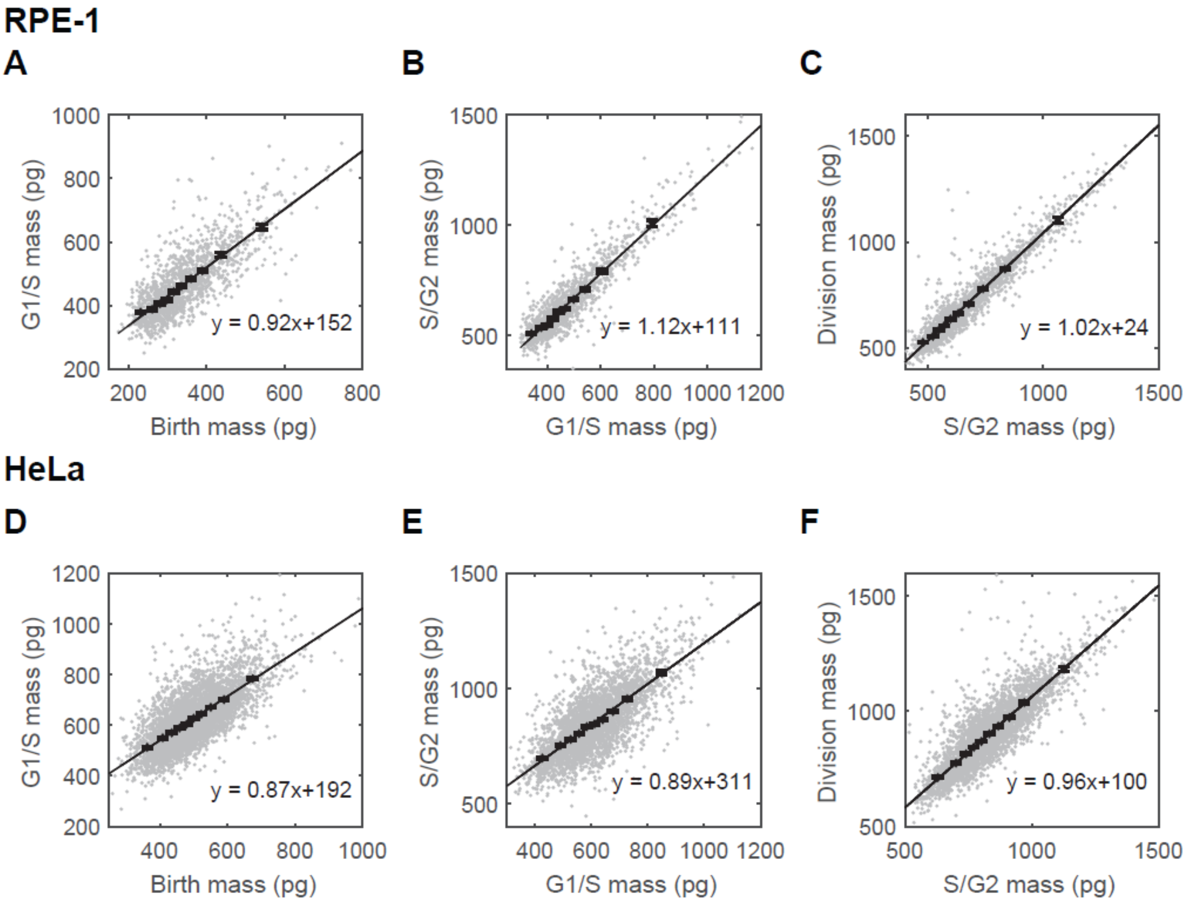

**Figure S7. The sequential adder behavior in RPE-1 and HeLa cells.** (A, D) The correlations between birth mass and mass at G1/S in RPE-1 (A) and HeLa (D) cells. (B, E) The correlations between mass at G1/S and mass at S/G2 in RPE-1 (B) and HeLa (E) cells. (C, F) The correlations between mass at S/G2 and division mass in RPE-1 (C) and HeLa (F) cells. Each gray dot is an observation; black squares are the average of each cell mass bin; error bars are the standard error of means (SEMs). Solid black lines are the best linear fits of the gray dots; Texts indicate the functions of the solid black lines.

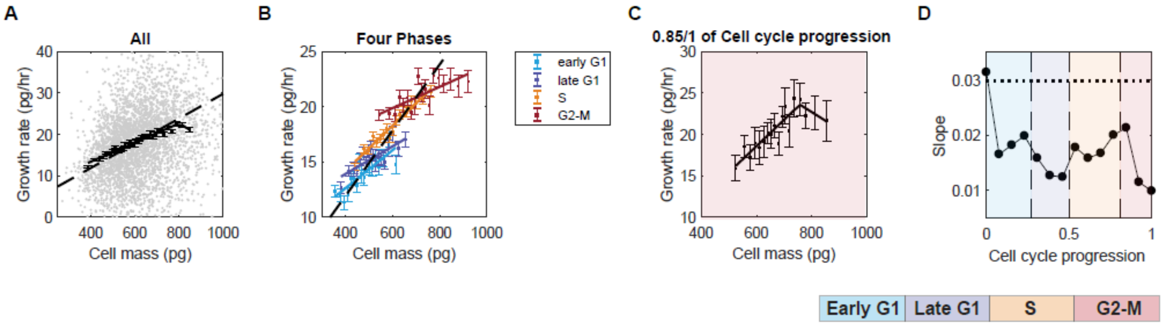

**Figure S8. Growth rate modulation in HeLa cells.** (A) The correlation between cell mass and growth rate in HeLa cells when pooling all the cells together. Each gray dot is an observation in the 3-hour measurements, n = 24,998. Black squares are the median growth rate of each mass bin; error bars are SEMs. The solid black line is the best fit of the black squares (Table S5). The dashed black line indicates exponential growth. (B, C) The correlations between cell mass and growth rate in HeLa cells in four cell cycle phases (B) and one fine stage of the cell cycle (C). The stages were determined by the log(Geminin) and DNA intensity using the ERA method(8), as indicated in Fig. S4C. Filled squares are the median growth rate of each mass bin; error bars are SEMs. The solid lines are the best fit of the filled squares (Table S5). The dashed black line in (B) indicates exponential growth. (D) The slope of the linear relationship between cell mass and growth rate plotted against cell cycle progression. The short dashed line indicates the expected slope for exponential growth.

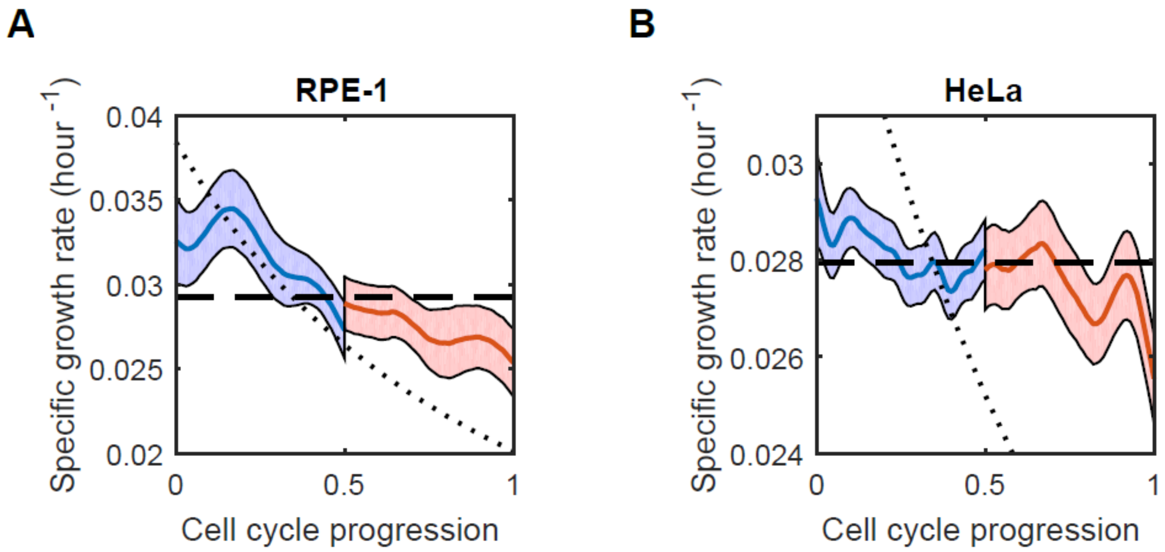

**Figure S9. Specific growth rate changes with cell cycle progression in RPE-1 (A) and HeLa cells (B) in G1 (blue) and nonG1 (red) phases.** Since the binned correlation could be affected by inspection bias(9), we investigated how the specific growth rate (growth rate divided by mass) changed with cell cycle progression from the long-term trajectories as recommended by Kar et al.(9). We arbitrarily assumed the G1 or nonG1 phase each occupies half of the cell cycle when normalizing the length of the growth trajectories. Solid blue and red lines are the mean of the normalized growth trajectories; the shaded areas indicate SEM. Dashed lines are the exponential growth; short dashed lines are the linear growth, assuming the cells behave like an adder.

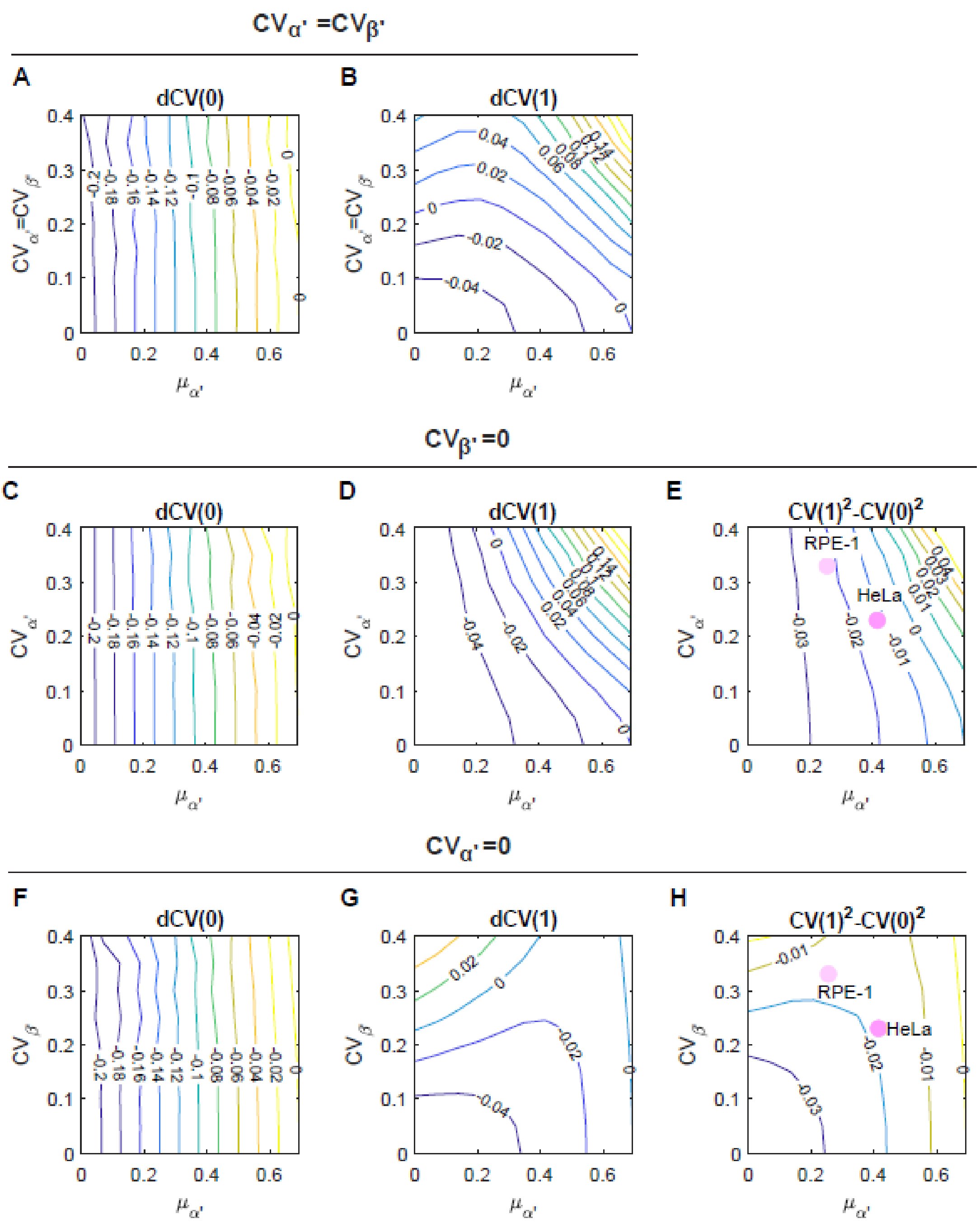

**Figure S10. Simulation results for the Sub-exponential growth rate modulation.** (A, C, F) Contour plots illustrating the rate of change in cell mass CV at the beginning of the cell cycle ($t' = 0$) when assuming $CV_{\alpha'} = CV_{\beta'}$ (A), $CV_{\beta'} = 0$ (C), or $CV_{\alpha'} = 0$ (F), respectively. Here, $\mu_{\alpha'}$ represents the mean of $\alpha'$. (B, D, G) Contour plots illustrating the rate of change in cell mass CV at the end of the cell cycle ($t' = 1$) when assuming $CV_{\alpha'} = CV_{\beta'}$ (B), $CV_{\beta'} = 0$ (D), or $CV_{\alpha'} = 0$ (G), respectively. (E, H) Contour plot illustrating the overall change in cell mass CV throughout the cell cycle when assuming $CV_{\beta'} = 0$ (E), or $CV_{\alpha'} = 0$ (H), respectively. Solid circles indicate the corresponding positions in the contour plots when adopting parameter values from the experimental observations of RPE-1 and HeLa cells.

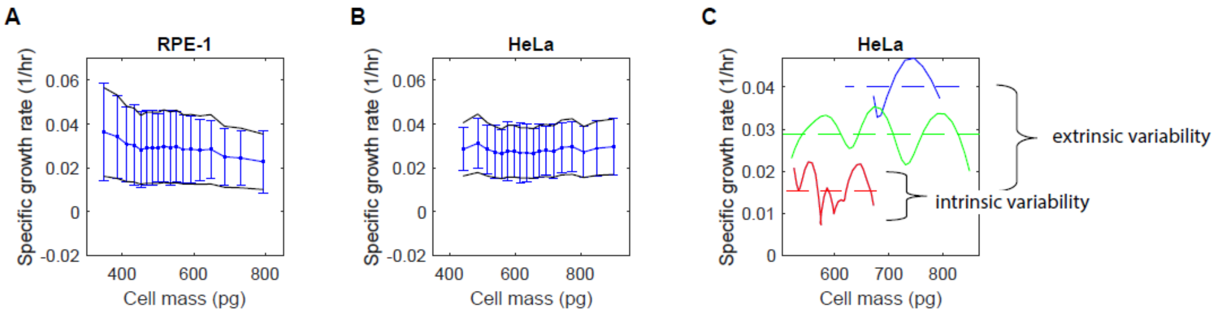

**Figure S11. Estimating the variability in $\alpha'$ for RPE-1 and HeLa cells.** (A-B) The variability of the specific growth rate, defined as the growth rate divided by cell mass, does not change with cell mass for RPE-1 (A) and HeLa (B) cells. Blue squares and lines indicate the means and standard deviations of live cell growth trajectories, which are binned by cell mass. The black lines show $(1 \pm \overline{CV})\overline{gr}_j$, where $\overline{CV}$ is the average CV in specific growth rate for all cell mass bins, and $\overline{gr_j}$ is the average specific growth rate for each bin. (C) Schematic illustrating the definitions of extrinsic and intrinsic variability in $\alpha'$. Solid lines are representative live cell growth trajectories. Dashes lines represent the means of each trajectory. Extrinsic variabilty is defined as the variaiton among the means of each trajectories, while intrinsic variability is defined as the fluctruation within individual trajectories.

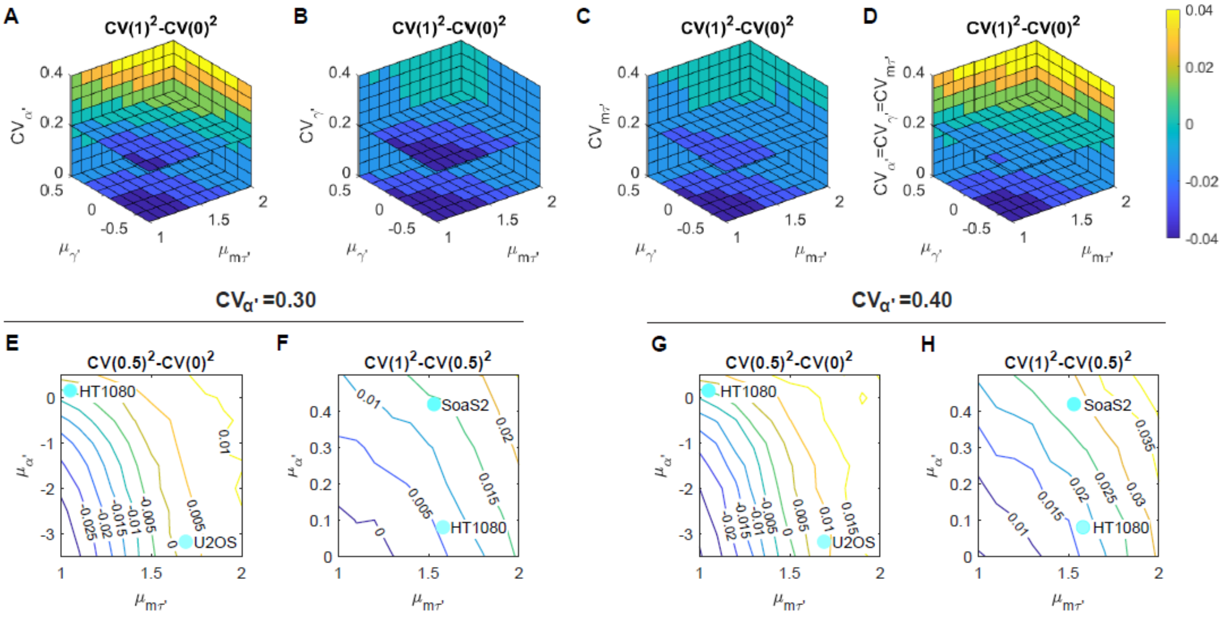

**Figure S12. Simulation results for the Bilinear growth rate modulation.** (A-D) Three-dimensional (3D) volumetric plot showing how the change in cell mass CV throughout the cell cycle responds to the means of $\gamma'$ and $m_\tau'$, represented by $\mu_{\alpha'}$ and $\mu_{m_\tau'}$, when applying variability to only one parameter of $\alpha'$ (A), $\gamma'$ (B), $m_\tau'$ (C), or equal variability to all three parameters (D). The slice planes are orthogonal to the CV axis at CV = 0.2. (E-F) Contour plots illustrating the change in cell mass CV during the G1 (E) and nonG1 phases (F) when assuming a 30% CV in $\alpha'$. (G-H) Contour plots illustrating the change in cell mass CV during the G1 (G) and nonG1 phases (H) when assuming a 40% CV in $\alpha'$. Solid circles in (E-H) indicate the corresponding positions in the contour plots when adopting parameter values from the experimental data.

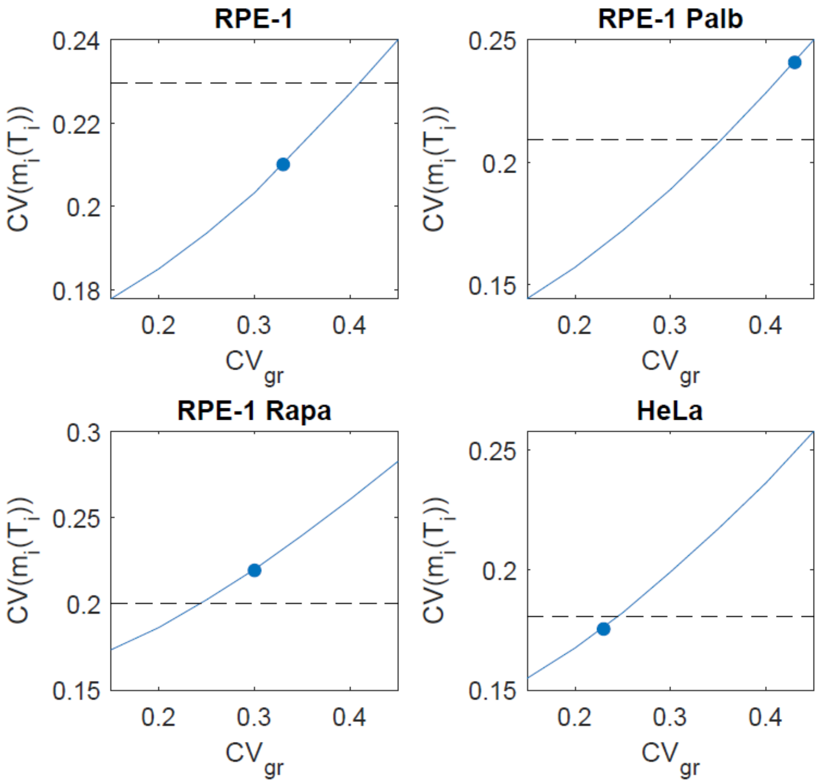

**Figure S13. Impact of growth rate variability on division mass CV, $CV(m_i(T_i))$, in the stochastic model.** The stochastic model is described in SI Text, Section 4, Scenario IX. All parameter values used in this simulation are listed in the Table at the end of Section 4, with the exception of $CV_{gr}$, which is varied in this simulation. Solid blue lines indicate the simulation results. Filled blue circles are the division mass CV when simulated with the $CV_{gr}$ estimated from experimental data. Dashed black lines represent the division mass CV measured in experiments.

**Supplementary tables**

**Table S1. Characteristics of the cell lines used in this study.**

| | RPE-1 | HeLa | U2OS | Saos-2 | HT1080 |
|---|---|---|---|---|---|
| morphology | epithelial | epithelial | epithelial | epithelial | epithelial |
| tissue | Eye; Pigmented epithelium; Retina | Uterus; Cervix | Bone | Bone | Connective tissue |
| disease | Normal | Adenocarcinoma | Osteosarcoma | Osteosarcoma | Fibrosarcoma |
| gender and age | Female, 1 | Female, 31 | Female, 15 | Female, 11 | Male, 35 |
| Karyotype | Modal chromosome number of 46, near-diploid | Modal number = 82, 1.76 fold of RPE-1* | 1.37 fold of RPE-1* | 1.33 fold of RPE-1* | 2.01 fold of RPE-1* |
| G1/S circuitry | Intact | inactivated pRb, p103, p170, p21, p27, p53 [1] | deficient p16, p53, Wip1; intact pRb[2-5] | deleted pRb and p53[2,3,5] | deleted p16; mutated p53 and N-ras; intact pRB[5-7] |

* measured by this study using Hoechst stain.

[1]Moody, Cary A., and Laimonis A. Laimins. "Human papillomavirus oncoproteins: pathways to transformation." Nature Reviews Cancer 10.8 (2010): 550-560.

[2]Diller, Lisa, et al. "p53 functions as a cell cycle control protein in osteosarcomas." Molecular and cellular biology 10.11 (1990): 5772-5781.

[3]Stott, Francesca J., et al. "The alternative product from the human CDKN2A locus, p14ARF, participates in a regulatory feedback loop with p53 and MDM2." The EMBO journal 17.17 (1998): 5001-5014.

[4]Kleiblova, Petra, et al. "Gain-of-function mutations of PPM1D/Wip1 impair the p53-dependent G1 checkpoint." Journal of Cell Biology 201.4 (2013): 511-521.

[5]Moolmuang, Benchamart, and Michael A. Tainsky. "CREG1 enhances p16INK4a-induced cellular senescence." *Cell Cycle* 10.3 (2011): 518-530.

[6]Anderson, Michael J., et al. "Evidence that wild-type TP53, and not genes on either chromosome 1 or 11, controls the tumorigenic phenotype of the human fibrosarcoma HT1080." Genes, Chromosomes and Cancer 9.4 (1994): 266-281.

[7]Brown, Robin, et al. "Mechanism of activation of an N-ras gene in the human fibrosarcoma cell line HT1080." The EMBO journal 3.6 (1984): 1321-1326.

**Table S2. The durations of cell cycle phases for HeLa, RPE-1, RPE-1 in 100 nM rapamycin or 50 nM palbociclib at cell mass homeostasis.** MAD is the median absolute deviation, and nMAD is MAD normalized by the median in robust statistics.

| | | mean (hour) | median (hour) | std (hour) | MAD (hour) | CV | nMAD |
|---|---|---|---|---|---|---|---|
| HeLa | Cell cycle | 28.0 | 27.5 | 4.6 | 3.6 | 0.16 | 0.13 |
| | G1 | 11.1 | 10.5 | 4.3 | 3.2 | 0.38 | 0.30 |
| | S | 11.3 | 11.0 | 4.2 | 3.1 | 0.37 | 0.28 |
| | G2-M | 5.8 | 5.5 | 2.7 | 2.0 | 0.46 | 0.36 |
| RPE-1 | Cell cycle | 18.0 | 18.0 | 3.1 | 2.3 | 0.17 | 0.13 |
| | G1 | 7.8 | 7.5 | 3.2 | 2.4 | 0.41 | 0.32 |
| | S | 10.0 | 9.5 | 3.2 | 2.4 | 0.32 | 0.25 |
| | G2-M | 3.1 | 3.0 | 3.2 | 1.7 | 0.54 | 0.39 |
| RPE-1 Rapa | Cell cycle | 32.6 | 32.0 | 8.2 | 7.0 | 0.25 | 0.22 |
| | G1 | 16.7 | 15.5 | 9.4 | 7.3 | 0.56 | 0.47 |
| | S | 14.1 | 13.5 | 4.4 | 3.2 | 0.31 | 0.24 |
| | G2-M | 3.9 | 3.5 | 2.5 | 1.6 | 0.64 | 0.46 |
| RPE-1 Palb | Cell cycle | 26.0 | 25.3 | 5.3 | 4.1 | 0.20 | 0.16 |
| | G1 | 12.3 | 11.0 | 7.0 | 5.3 | 0.57 | 0.48 |
| | S | 11.9 | 11.0 | 6.8 | 5.0 | 0.58 | 0.45 |
| | G2-M | 2.8 | 2.4 | 2.2 | 1.3 | 0.77 | 0.55 |

**Table S3. Comparing cell cycle phase duration and mass versus phase length correlation with and without the mTurquoise2-SLBP marker in HeLa cells.**

| | | HeLa mAG-hGeminin | HeLa mAG-Geminin mTurq2-SLBP |
|---|---|---|---|
| Median duration (hour) | Cell cycle | 26.0 | 27.5 |
| | G1 | 10.5 | 10.5 |
| | nonG1 | 16.5 | 16.5 |
| Mass-length Pearson correlation | Birth mass-Cell cycle | -0.27 | -0.26 |
| | Birth mass-G1 | -0.19 | -0.20 |
| | G1/S mass-nonG1 | -0.32 | -0.34 |

**Table S4. Comparison of the linear and bilinear fits for the cell mass vs. cell cycle phase length correlations.** The significantly better fits (p_bilinear or p_linear < 0.05) and the significant negative correlations (p<0.05) are highlighted.

| | | AICc_bilinear | AICc_linear | p_bilinear | p_linear | R | p-Value of R |
|---|---|---|---|---|---|---|---|
| HeLa | Birth mass-G1 length | -61.8 | -41.8 | 4.6E-05 | | -0.20 | 1.4E-84 |
| | G1/S mass-S length | -65.4 | -14.4 | 8.3E-12 | | -0.29 | 3.1E-141 |
| | S/G2 mass-G2-M length | -103.1 | -79.4 | 7.1E-06 | | -0.18 | 8.5E-66 |
| | Birth mass-cell cycle length | -22.8 | -6.1 | 2.4E-04 | | -0.26 | 5.8E-61 |
| RPE-1 | Birth mass-G1 length | -45.0 | -39.9 | 0.07 | | -0.25 | 3.6E-55 |
| | G1/S mass-S length | -43.4 | -22.7 | 3.2E-05 | | -0.12 | 3.5E-12 |
| | S/G2 mass-G2-M length | -91.9 | -92.9 | | 0.58 | -0.08 | 2.3E-13 |
| | Birth mass-cell cycle length | -15.2 | -8.0 | 0.03 | | -0.25 | 1.6E-12 |

**Table S5. Comparison of the linear and bilinear fits for the cell mass vs. growth rate correlations.** The significantly better fits (p_bilinear or p_linear < 0.05) are highlighted.

| | | AICc_bilinear | AICc_linear | p_bilinear | p_linear |
|---|---|---|---|---|---|
| HeLa | All | -41.2 | -30.0 | 0.003 | |
| | G1 | -12.3 | -17.2 | | 0.09 |
| | nonG1 | -13.0 | -14.0 | | 0.61 |
| | early G1 | -2.5 | -5.5 | | 0.22 |
| | late G1 | -5.4 | -11.1 | | 0.06 |
| | S | -19.8 | -27.7 | | 0.02 |
| | G2-M | 3.0 | -3.4 | | 0.04 |
| | Stage 0.85/1 | 13.7 | 16.0 | 0.32 | |
| RPE-1 | G1 | -3.1 | -11.4 | | 0.02 |
| | nonG1 | -15.9 | -25.0 | | 0.01 |
| U2OS | G1 | 21.0 | 25.9 | 0.09 | |
| | nonG1 | -6.4 | -11.1 | | 0.09 |
| HT1080 | G1 | 9.7 | 13.2 | 0.18 | |
| | nonG1 | -0.7 | 9.4 | 0.007 | |
| Saos-2 | G1 | -28.5 | -31.0 | | 0.29 |
| | nonG1 | -41.5 | -35.7 | 0.05 | |
| U2OS DMEM | G1 | 26.5 | 29.1 | 0.28 | |
| | nonG1 | -10.8 | -18.9 | | 0.02 |
| U2OS DMEM Rapa | G1 | -0.9 | -8.9 | | 0.02 |
| | nonG1 | -26.6 | -34.5 | | 0.02 |
| U2OS DMEM MG132 | G1 | 16.2 | 4.5 | | 0.003 |
| | nonG1 | -19.1 | -25.4 | | 0.04 |
| U2OS DMEM CHX | G1 | -12.6 | -9.9 | 0.24 | |
| | nonG1 | -22.2 | -31.3 | | 0.01 |
| RPE-1 2% FBS | G1 | 6.4 | -2.1 | | 0.01 |
| | nonG1 | 6.0 | -2.2 | | 0.02 |

**Table S6. The normalized fitting parameters for the cell mass vs. growth rate correlations for different cell lines.** For correlations fitted better by the linear model, $\frac{dm}{dt} = \alpha m + \beta$, the normalized parameters $\alpha'$ and $\beta'$ are listed in the table, with $\alpha' = \alpha < T >$, $\beta' = \beta \frac{<T>}{<m_b>}$, where $T$ and $m_b$ are the cell cycle length and cell birth mass, respectively. For exponential growth, $\alpha' = ln2 = 0.693$. For correlations fitted better by the bilinear model, $\frac{dm}{dt} = (am + b)(m < m_\tau) + (\gamma m + am_\tau + b - \gamma m_\tau)(m \geq m_\tau)$, the normalized parameters $a'$, $b'$, $\gamma'$, and $m_\tau'$ are listed in the table, with $a' = a < T >$, $b' = b \frac{<T>}{<m_b>}$, $\gamma' = \gamma < T >, m_\tau' = \frac{m_\tau}{<m_b>}$. The correlation slopes, $\alpha'$, $a'$, and $\gamma'$, lower than 0.75 or higher than 1.25 fold (arbitrarily chosen thresholds) of $ln2$ were highlighted. SE and BI denote the type of growth rate modulation, where SE stands for Sub-exponential and BI stands for Bilinear.

| | | $\alpha', \beta'$ | $a', b', \gamma', m_\tau'$ | Modulation type |
|---|---|---|---|---|
| RPE-1 | G1 | 0.25, 0.44 | | SE |
| | nonG1 | 0.26, 0.57 | | SE |
| HeLa | G1 | 0.40, 0.36 | | SE |
| | nonG1 | 0.43, 0.43 | | SE |
| U2OS | G1 | | 0.67, -0.16, -3.18, 1.69 | BI |
| | nonG1 | 0.67, 0.15 | | None |
| HT1080 | G1 | | 0.85, -0.10, 0.16, 1.05 | BI |
| | nonG1 | | 0.54, 0.32, 0.08, 1.58 | BI |
| Saos-2 | G1 | 0.84, -0.15 | | None |
| | nonG1 | | 0.96, -0.30, 0.42, 1.53 | BI |

**Table S7. The values of $\lambda'$ and $\alpha'$ used in Fig. 5L, for untreated HeLa and RPE-1 cells, as well as RPE-1 cells treated with 50 nM palbociclib or 100 nM rapamycin.**

| | $\lambda'$ | $\alpha'$ |
|---|---|---|
| HeLa | -0.47 | 0.41 |
| RPE-1 | -0.22 | 0.25 |
| RPE-1 Palb | -0.15 | -0.002 |
| RPE-1 Rapa | -0.45 | 0.55 |

**Table S8: Contribution of each factor to cell mass variation, as indicated by the division mass CV simulated using the stochastic model (SI Text, Section 4).** The values reported in this table are the average division mass CV obtained from 50 simulations.

| | HeLa | RPE | RPE Palb | RPE Rapa |
|---|---|---|---|---|
| I. without noise or control mechanisms | 0.18 | 0.23 | 0.21 | 0.20 |
| II. with partition noise, without control mechanisms | 0.21 | 0.25 | 0.24 | 0.22 |
| III. with cell cycle variation, without control mechanisms | 0.22 | 0.26 | 0.24 | 0.24 |
| IV. with growth rate variation, without control mechanisms | 0.24 | 0.33 | 0.37 | 0.29 |
| V. with all noise, without control mechanisms | 0.30 | 0.38 | 0.43 | 0.35 |
| VI. with all noise and G1 length control | 0.28 | 0.35 | 0.41 | 0.28 |
| VII. with all noise and nonG1 length control | 0.25 | 0.37 | 0.40 | 0.34 |
| VIII. with all noise and growth rate control | 0.23 | 0.24 | 0.26 | 0.28 |
| IX. with all noise and all control mechanisms | 0.18 | 0.21 | 0.24 | 0.22 |

**Table S9. The frequencies of cell death, cell cycle arrest, and cytoplasmic loss observed in the long–term measurements in HeLa, RPE-1, RPE-1 in 100 nM rapamycin, and RPE-1 in 50 nM palbociclib when cells have reached cell mass homeostasis.**

| | HeLa | RPE | RPE Rapa | RPE Palb |
|---|---|---|---|---|
| Death | ~2% | <0.1% | <0.1% | ~2% |
| Cell cycle arrest | ~0.2% | ~0.2% | <0.1% | ~0.3% |
| Cytoplasmic loss during mitosis | <0.1% | <0.1% | <0.1% | ~0.5% |

**Table S10. Birth size CV, division size CV, and DA std. reported in the literature.**

| | Birth size CV | Division size CV | DA std. |
|---|---|---|---|
| *C. crescentus*[1] | 16-18% | 12% | |
| *E. coli* | 12%[1] | 11%[1] | 1%[3] |
| *M. smegmatis*[2] | 19% | | |
| *M. bovis* BCG[2] | 20% | | |
| *H. salinarum*[3] | 16% | 13% | 3% |
| *S. cerevisiae*[4] | 24% | 23% | |
| *S. pombe*[5] | | 6% | 1.6% |
| *Arabidopsis* shoot stem cell[6] | 24% | 13% | |
| L1210 | 25%[7], 8%[9] | 7%[9] | 4.8%[10] |
| MOLT4[7] | 29% | | |
| RBL[8] | 11% | 11% | |
| RAW 264.7[8] | 16% | | |
| FL5.12[9] | 13% | 11.50% | |
| RKO[10] | | | 6.8% |
| HT-29[10] | | | 6.4% |
| L-929[11] | 16% | 12% | 4.8% |

[1]Campos, Manuel, et al. "A constant size extension drives bacterial cell size homeostasis." Cell 159.6 (2014): 1433-1446.
[2]Logsdon, Michelle M., et al. "A parallel adder coordinates mycobacterial cell-cycle progression and cell-size homeostasis in the context of asymmetric growth and organization." Current Biology 27.21 (2017): 3367-3374.
[3]Eun, Ye-Jin, et al. "Archaeal cells share common size control with bacteria despite noisier growth and division." Nature microbiology 3.2 (2018): 148-154.
[4]Barber, Felix, Ariel Amir, and Andrew W. Murray. "Cell-size regulation in budding yeast does not depend on linear accumulation of Whi5." Proceedings of the National Academy of Sciences 117.25 (2020): 14243-14250.
[5]Sveiczer, A., B. Novak, and J. M. Mitchison. "The size control of fission yeast revisited." Journal of cell science 109.12 (1996): 2947-2957.
[6]D'Ario, Marco, et al. "Cell size controlled in plants using DNA content as an internal scale." Science 372.6547 (2021): 1176-1181.
[7]Tzur, Amit, et al. "Cell growth and size homeostasis in proliferating animal cells." Science 325.5937 (2009): 167-171.
[8]Varsano, Giulia, Yuedi Wang, and Min Wu. "Probing mammalian cell size homeostasis by channel-assisted cell reshaping." Cell reports 20.2 (2017): 397-410.
[9]Son, Sungmin, et al. "Direct observation of mammalian cell growth and size regulation." Nature methods 9.9 (2012): 910-912.
[10]Sung, Yongjin, et al. "Size homeostasis in adherent cells studied by synthetic phase microscopy." Proceedings of the National Academy of Sciences 110.41 (2013): 16687-16692.

[11]Killander, D., and A. Zetterberg. "Quantitative cytochemical studies on interphase growth: I. Determination of DNA, RNA and mass content of age determined mouse fibroblasts in vitro and of intercellular variation in generation time." Experimental cell research 38.2 (1965): 272-284.

**Legend for Movie S1**

Time-lapse quantitative phase images of RPE-1 cells in 50nM palbociclib; the time interval is 30 minutes; the yellow arrow indicates the lost cytoplasmic mass of a mitotic cell (red arrow); the scale bar indicates 100 μm.
